# Quantifying the Local Adaptive Landscape of a Nascent Bacterial Community

**DOI:** 10.1101/2022.02.03.475969

**Authors:** Joao A Ascensao, Kelly M Wetmore, Benjamin H Good, Adam P Arkin, Oskar Hallatschek

## Abstract

The fitness effects of all possible mutations available to an organism largely shapes the dynamics of evolutionary adaptation. Tremendous progress has been made in quantifying the strength and abundance of selected mutations available to single microbial species in simple environments, lacking strong ecological interactions. However, the adaptive potential of strains that are part of multi-strain communities remains largely unclear. We sought to fill this gap by analyzing a stable community of two closely related ecotypes (“L” and “S”) shortly after they emerged within the *E. coli* Long-Term Evolution Experiment (LTEE). We engineered genome-wide barcoded transposon libraries to measure the fitness effects of all possible gene knockouts in the coexisting strains as well as their ancestor, for many different, ecologically relevant conditions. We found that the fitness effects of many gene knockouts sensitively depends on the genetic background and the ecological conditions, as set by the abiotic environment and relative frequency of both ecotypes. Despite the idiosyncratic behavior of individual knockouts, we still see consistent statistical patterns of fitness effect variation across both genetic background and community composition. Genes that are in the same operon, or that strongly interact with each other, are more likely to be correlated with each other across backgrounds compared to random pairs of genes. Additionally, fitness effects are correlated with evolutionary outcomes for a number of conditions, possibly revealing shifting patterns of adaptation. Together, our results reveal how ecological and epistatic effects combine to drive adaptive potential in a nascent ecological community.

## Introduction

Microbial communities are ubiquitous across all environments, and are key players in disease processes, biogeochemical cycling, and ecosystem functioning (1–6). While most research on natural microbiomes has been fueled by their ecological significance, recent studies have begun to focus on microbial community evolution and uncovered clear signs of adaptation and diversification (7–10). Thus, microbiome assembly, structure, and function may have to be understood against a backdrop of an ever-churning evolutionary dynamics.

That evolutionary and ecological changes often go together has been most clearly shown in controlled experiments on synthetic microbial communities: evolution can change the way microbes consume resources or otherwise interact with each other (11–15). This leads to environmental changes that modify selection pressures, forcing lineages into new evolutionary paths (16–21). Complex adaptive landscapes have been hypothesized to chiefly shape the feedback between ecology and evolution in microbial communities (19, 22), but it is still unclear how diversification and other ecological shifts change those landscapes.

In ecologically simple monoculture populations, population genetic theory has shown that the evolutionary dynamics are largely predictable from knowing local aspects of a static fitness landscape, encoding the fitness effects of all currently available mutations, which is called the “distribution of fitness effects” (DFE) (23–28). Such work has been successful in rationalizing and predicting outcomes of evolution experiments from DFE measurements (29, 30).

High-quality measurements of the DFE in a given system require sampling and measuring the fitness effects of sufficiently many mutations across the genome. This has only become possible recently, due to the rise of sequencing technologies. DNA barcoding systems have become especially influential to better understand microbial adaptive evolution. By taking advantage of amplicon sequencing methods to measure barcode frequency dynamics, these systems have been used with great success to directly observe evolutionary dynamics (30–33), and identify selected mutations and the statistical patterns that characterize them (34–39).

However, the concept of a *single, static* DFE may not be applicable or useful to describe a diversified population. It is possible that different ecotypes experience different adaptive landscapes, even if they are closely related, which moreover may shift in response to compositional or other ecological changes. Despite the importance of microbial communities, very little is known about how much the local landscape depends on biotic interactions with their coexisting strains versus genetic background alone, and how those patterns shift upon diversification.

Here we aim to elucidate the adaptive landscape of a microbial community by measuring how the invasion fitness effects of a large panel of mutations depends on the state of the ecosystem. Invasion fitness refers to the growth rate of a mutant relative to its ancestor when the mutant is rare in the population. To sample from the DFE, we create genome-wide knockout libraries via random-barcoded transposon mutagenesis (40, 41) on the backgrounds of the coexisting ecotypes.

While knockout mutations do not represent all possible mutations in the genome, this approach allows us to sample a wide variety of mutations across the genome and to compare the effect of the same mutation across different genetic backgrounds and community compositions. The resulting ecotype-, and composition-dependent DFE statistically characterizes the abundance and specificity of beneficial mutations and, thus, reveals how the rate and pattern of mutation accumulation depends on the state of the ecosystem.

We reasoned that the ecologically-dependent DFEs accessible by our approach are particularly relevant to the fate of a recently diversified ecosystem, consisting of closely related ecotypes with overlapping niches. Additionally, quantifying the DFEs of such a nascent community would shed light on how the discovery and infiltration of a new niche changes the local adaptive landscape, in both focal and “nearby” environments. The composition-dependence of the DFE would also provide information on the types of mutations available to the community–”pure fitness” mutations would show minimal fitness changes in response to composition shifts, whereas frequency-dependent mutations may point to shifts in niche occupation/strategy. Theory suggests that the relative availability of “pure fitness” versus frequency-dependent mutations may strongly influence the resulting evolutionary dynamics, but there have been few empirical measurements of how many mutations show frequency-dependent effects (19).

We therefore chose to focus on a model ecosystem that spontaneously emerge.d in the course of the *E. coli* Long Term Evolution Experiment (LTEE) – an experiment that has tracked the evolution of several *E. coli* populations over the course of over 70,000 generations (at the time of writing). Early in the LTEE, it was recognized that one of the twelve lineages, the ara-2 population on which we focus in this study, spontaneously diversified into two lineages that coexist via negative frequency dependence, termed S and L (for their small and large colony sizes on certain agar plates) (42). S and L coexist by inhabiting different temporal/metabolic niches in the LTEE environment, set up as serial dilutions in glucose minimal media–L grows more quickly on glucose during exponential phase, while S specializes on stationary phase survival and utilizing acetate, a byproduct of overflow metabolism (43, 44). Following diversification, the lineages have persisted to this day and continued to evolve and adapt, diverging on genetic, transcriptional, and metabolic levels (16, 42–47). While our focal ara-2 line is the best studied case of diversification in the LTEE, it is not the only one. Recent time-resolved metagenomic sequencing of the LTEE has shown that, in fact, 9 out of the 12 populations evolved two separate lineages that coexisted with each other for tens of thousands of generations, while continuing to accumulate mutations and adapt (47), demonstrating that spontaneous diversification followed by coevolution is a major adaptive route for this system.

## Results

### Measuring knockout fitness effects

We sought to measure the knockout fitness effects available to the small LTEE-derived ecosystem of S and L, and how they depend on ecological conditions, specifically, (i) the composition of the community, and (ii) openness of a given metabolic niche. To this end, we created randomly barcoded transposon libraries of three LTEE clones, using previously developed methods (RB-TnSeq) (40, 41)–S and L clones sampled from 6.5k generations, right after diversification (16, 42), and their LTEE ancestor, REL606 (Figure 1A). We used these libraries to measure the knockout fitness effects of nearly all nonessential genes in various environments relevant to the evolution of the population in the LTEE (Table 1), by propagating the libraries in defined conditions (with two biological replicates per experiment) and using Illumina amplicon sequencing to track the frequency trajectories of different barcodes (Figure 1B). By essentially measuring the log-slope of the frequency trajectories, we can estimate the fitness effect, *s*, of a given mutant (Figure 1C), which we report in units of 1*/generation*. Transposon insertion events were highly redundant, with a median of ∼20 insertions per gene, allowing us to combine information from multiple barcode trajectories into one fitness measurement through our statistical fitness inference pipeline and identify significantly non-neutral mutations (FDR correction; *α* = 0.05). We carefully quantified sources of error in barcode frequency measurements and propagated them to our fitness estimates, which was crucial to effective and accurate analysis of the data (see supplement section S3)–for example, we could exclude knockouts with overly noisy fitness measurements, or weight measurements by their error.

**Table 1.**
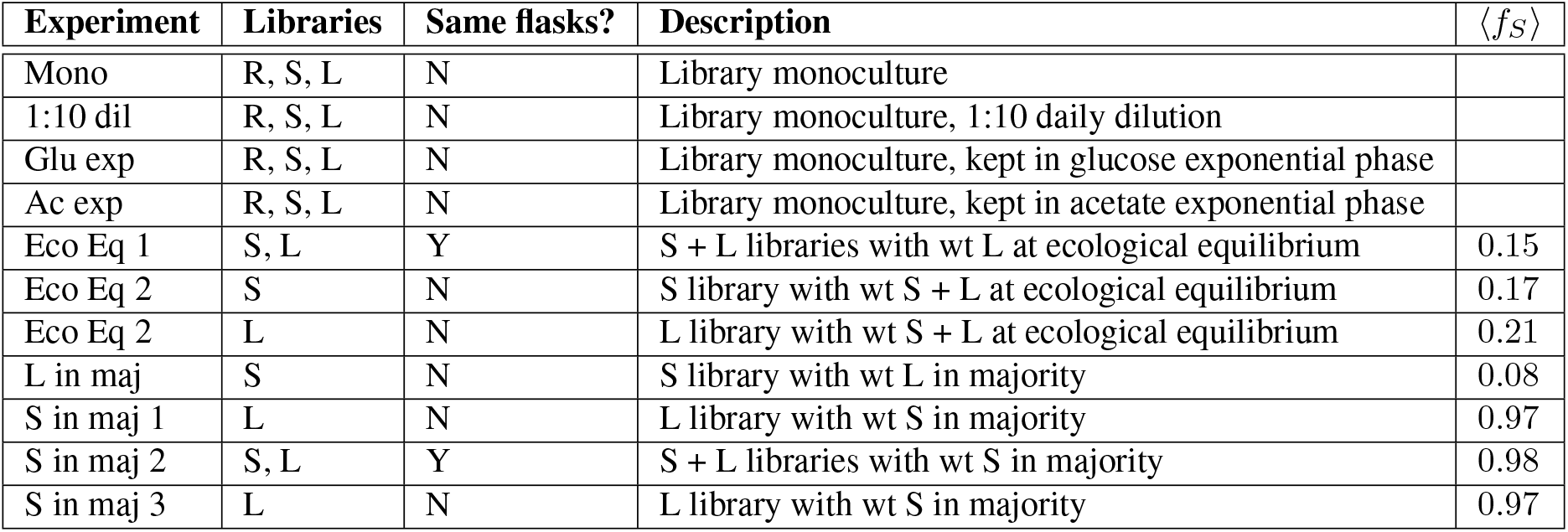
Summary of BarSeq experiments reported in this work. Dilution rate was variable in the glucose/acetate exponential phase experiments, to keep the populations in exponential phase (see supplement section S2), but unless otherwise noted, the daily dilution rate was 1:100, consistent with the LTEE condition. All experiments were performed in the LTEE media, DM25, except for the acetate exponential phase experiment. The abbreviations R, S, and L refer to REL606, and 6.5k S/L, respectively. In coculture experiments, (*f*_*S*_) is the total frequency of S, averaged over all time points and replicates.

**Fig. 1.**
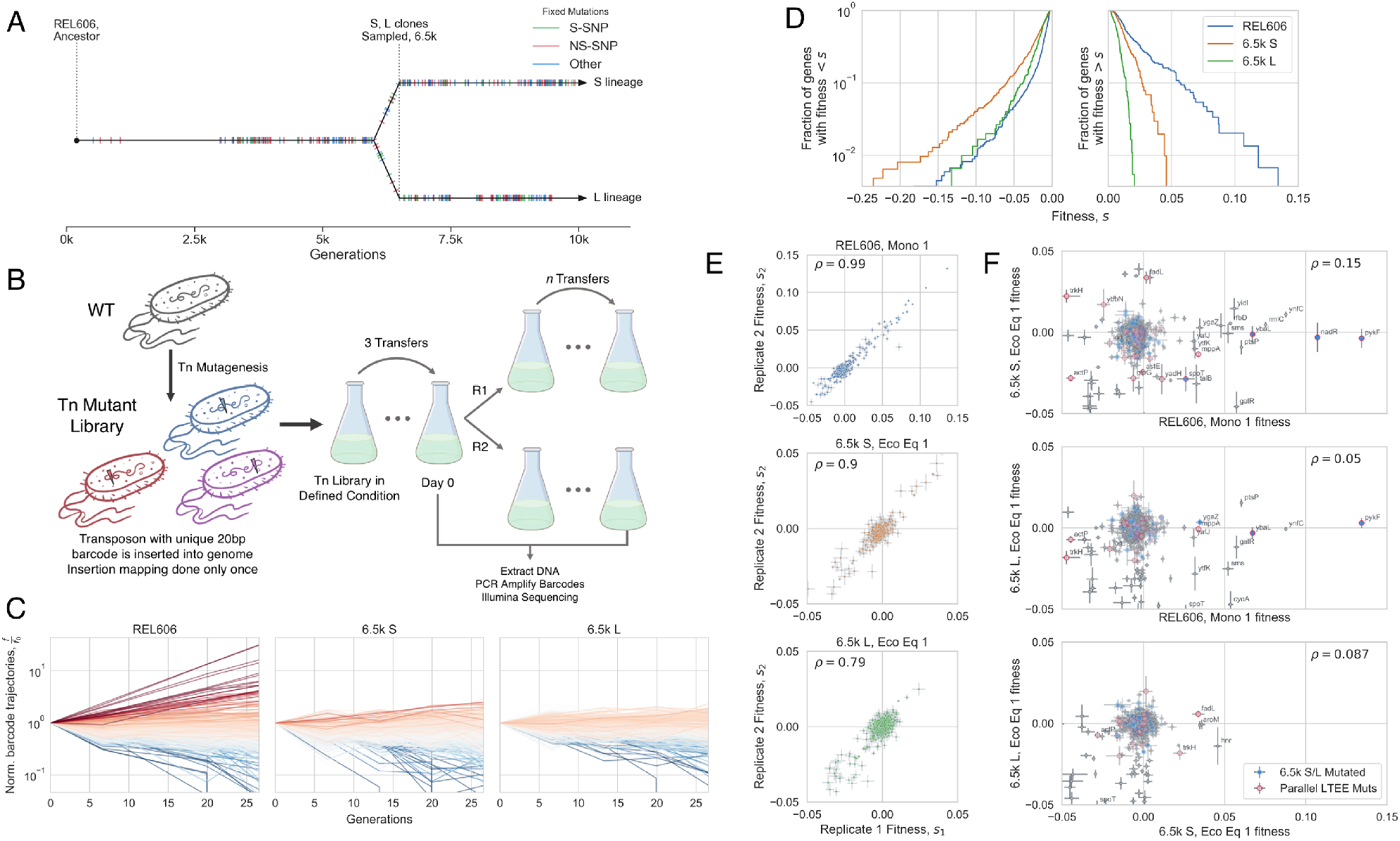
Measuring mutational fitness effects. (**A**) Timeline of evolution in the ara-2 LTEE population, showing mutation accumulation and diversification into S and L around 6k generations, then clone sampling at 6.5k generations; data from (47), jitter added to mutation fixation time for easier visualization. Hypermutator phenotype appeared around 2.5k generations (48). (**B**) Schematic of transposon mutagenesis process to generate barcoded libraries of REL606, 6.5k S and L, as well as experimental procedure to observe barcode dynamics. (**C**) Barcoded knockout mutant frequency trajectories in the evolutionary condition for each genetic background, colored by estimated fitness. All barcodes within a gene were summed together; shown are the trajectories from replicate 1 in the evolutionary condition for each genetic background (monoculture for REL606, together at ecological equilibrium frequency for S and L; representative of both replicates). (**D**) Overall distributions of fitness effects in the evolutionary condition for each genetic background. The majority of knockouts were neutral, so only genes that were called as significantly non-neutral were included (see supplement section S3.2). (**E**) Replicate-replicate correlation of estimated fitness effects. (**F**) Comparison of knockout fitness effects across genetic backgrounds, which are generally uncorrelated. Points with a blue interior correspond to genes that were mutated (excluding S-SNPs) in 6.5k S/L relative to REL606 (sequencing data from (46)). Points with red outlines correspond to genes that were mutated in parallel in nonmutator LTEE populations (data from (47)). The correlation coefficients decrease slightly if we recompute them, excluding likely neutral genes (*ρ* = 0.14, 0.03, 0.03; top to bottom). In panels E-F, knockouts with high measurement noise (*σs >* 0.3%) were excluded (except for labeled genes), and *ρ* is the weighted pearson correlation coefficient. Also in panels E-F, the “cloud” of points around 0 mostly represents likely neutral knockouts.

Barcoded transposon mutagenesis has been successfully and consistently used to measure knockout fitness effects across many contexts (40, 41), but as the knockouts are not bonafide deletions, it is possible that some genes with transposon insertions retain some activity. However, the fact that we have multiple transposon insertions spread across the length of each gene, along with our outlier barcode detection scheme, allows us to be more confident that our fitness measurements are dominated by the typical effects of an insertion.

After inferring the fitness effect of each gene knockout, we can compare fitness effects across genetic backgrounds and environments. We can first look at knockout fitness effects in the evolutionary condition proxies–the closest approximation to the environment where evolution in the LTEE took place: the REL606 library in monoculture, and S and L libraries together, coexisting at the ecological equilibrium frequency. We chose to highlight the condition where S and L were coexisting at their ecological equilibrium to be able to distinguish environmental versus genetic contributions to fitness effects–the libraries were cocultured together, in the same flasks, thus experiencing the same environment. In coculture experiments, the S/L libraries are mixed in the minority together with wild-type S/L clones at the desired frequency (see supplement section S2). The ecotype frequencies do not change considerably over the time period considered (Figure S7).

If we look at the overall DFE in the evolutionary condition proxies, we see that REL606 has access to beneficial knockouts of much larger effect size than either S or L (Figure 1D), suggesting that REL606 would adapt much quicker than S or L. Additionally, S has a larger beneficial DFE compared to L, which may be because S is starting to exploit an under-utilized niche (acetate specialization), where more significant gains can be made by improving the exploitation of the niche. On the other hand, L has inherited the putative old niche (glucose specialization), which was presumably the primary target of adaptation during the first ∼6k generations of evolution. As previously mentioned, the overall shape of the DFE largely controls the instantaneous speed of adaptation (23–28). The evolutionary tendency towards a “shrinking DFE” is known as global diminishing returns epistasis, which has previously been proposed as a mechanism to explain the decelerating fitness trajectories of the LTEE populations (49, 50). While diminishing returns epistasis was previously observed to affect the first couple common LTEE mutations (51), global diminishing returns (affecting the whole DFE) after the accumulation of many mutations had not yet been directly observed.

We can also compare the fitness effects of each knockout mutation both between replicates and across genetic backgrounds (Figure 1E-F), to contrast within-sample to between-sample variance. In contrast to a strong replicate-replicate correlation, we see that fitness effects are largely uncorrelated between genetic backgrounds. It may be unsurprising that mutational effects of S and L are uncorrelated with those of their ancestor, as REL606 may be creating and experiencing a slightly different environment compared to S and L, even though they were all started in the same media. However, as previously mentioned, we measured the fitness effects of S and L while they were coexisting in the same flasks, so the two ecotypes were experiencing the exact same environment. Thus, the lack of correlation between the fitness effects of S and L must be due to epistatic effects. It appears that individual mutations behave idiosyncratically despite statistical patterns of epistasis, in contrast with previous experiments (51, 52) which saw diminishing returns both globally and with individual mutations. Most knockout mutations that were strongly beneficial in REL606 and then acquired a mutation in that gene in the 6.5k S/L background became effectively neutral when knocked out in S/L (*nadR, pykF, ybaL, ygaZ*); it makes sense that mutating a gene that was already mutated (with a fitness effect) wouldn’t have an effect, if the mutation was effectively a loss-of-function. One gene, *spoT*, was beneficial in REL606 but deleterious in both S and L when knocked out, indicating that the natural *spoT* SNP may represent a change-of-function rather than a loss-of-function mutation. However, the majority of selected genes in REL606 were not mutated between 0 and 6.5k generations in S/L, so the fact that their fitness effects significantly changed across genetic background implicates the role of widespread, global idiosyncratic epistasis. Furthermore, there are several genes that were mutated in parallel in multiple lines of the LTEE, but are only beneficial on the S background (*trkH, ybbN*) or both the S and L backgrounds (*fadL*) when knocked out, while being neutral or deleterious on the REL606 background, suggesting that predictable epistasis could have shaped which mutations became beneficial in the LTEE. ‘Coupon collection’ is a null model of mutation accumulation/epistasis, where a beneficial DFE is composed of a finite number of mutations, and only changes due to the depletion of those mutations when they fix in a population. While the coupon collecting model is clearly relevant for some mutations, the lack of fitness effect correlation between genetic backgrounds seems to be largely driven by global epistasis.

As a simple check, we compared the fitness effect of one of the largest effect knockouts in our collection, *pykF*, to previously collected data. We reanalyzed data from Peng et al. (2018) Mol Biol Evol (53) (to recalculate fitness using the metric that we use) and found that their pykF deletion mutant had a selective coefficient *s ≈* 4%, compared to our measurement *s ≈* 12%; the highest fitness effect of a pykF nonsynonymous mutation on the ancestral background was 0 *s* ≈ 9%, which is similar to our measurement. Additionally, our measured fitness effect of pykF is quite consistent—it is approximately the same across all replicates in the Mono 1 and 2 experiments in REL606 (performed on different days). And all of the individual barcodes that landed in pykF appear to have approximately the same slopes. One possibility to describe the discrepancy could be the presence of frequency dependent fitness effects—the strength of selection may be higher when the mutant is rare (as is the case in our data), compared to when it occupies a sizable portion of the population (as in Peng et al. 2018). Another possibility could be that transposon insertions did not completely eliminate pykF activity, as it would in a deletion.

### Knockout fitness effects strongly depend on ecological conditions

The ecological interactions between S and L are mediated through the environment, most likely primarily through cross-feeding (43, 44). Therefore, it’s reasonable to think that the environment will change with the ecosystem composition, which could be modified by both ecological and evolutionary processes–indeed, ecotype composition does change significantly and relatively rapidly over evolutionary time (∼1k-10ks generations) (42, 47). Thus, we sought to explore how mutational fitness effects varied with ecosystem composition (Figure S7). Notably, we see a consistent trend where fitness effects generally have a smaller magnitude when S and L are in monoculture compared to when they are in coculture (Figures 2A, S11A). Additionally, we also see that the overall shape of the DFEs change as a function of frequency, with generally larger fitness effects when the ecotype is in the minority, for both beneficial and deleterious knockouts (Figures 2B, S11B). Analogous to the case of global diminishing returns epistasis, this observation holds on a statistical level, but does not explain all of the fitness effect variation between the different conditions, implying that individual mutations are affected by the ecosystem composition in idiosyncratic ways–statistical properties of the DFE seem to be strongly dependent on the ecosystem composition, but the effects of individual mutations may depend on their underlying physiological consequences and how they affect ecological interactions. Thus, it appears that the impact of both ecotypes on the environment is different enough to make selection pressures strongly dependent on the current mixture of ecotypes.

**Fig. 2.**
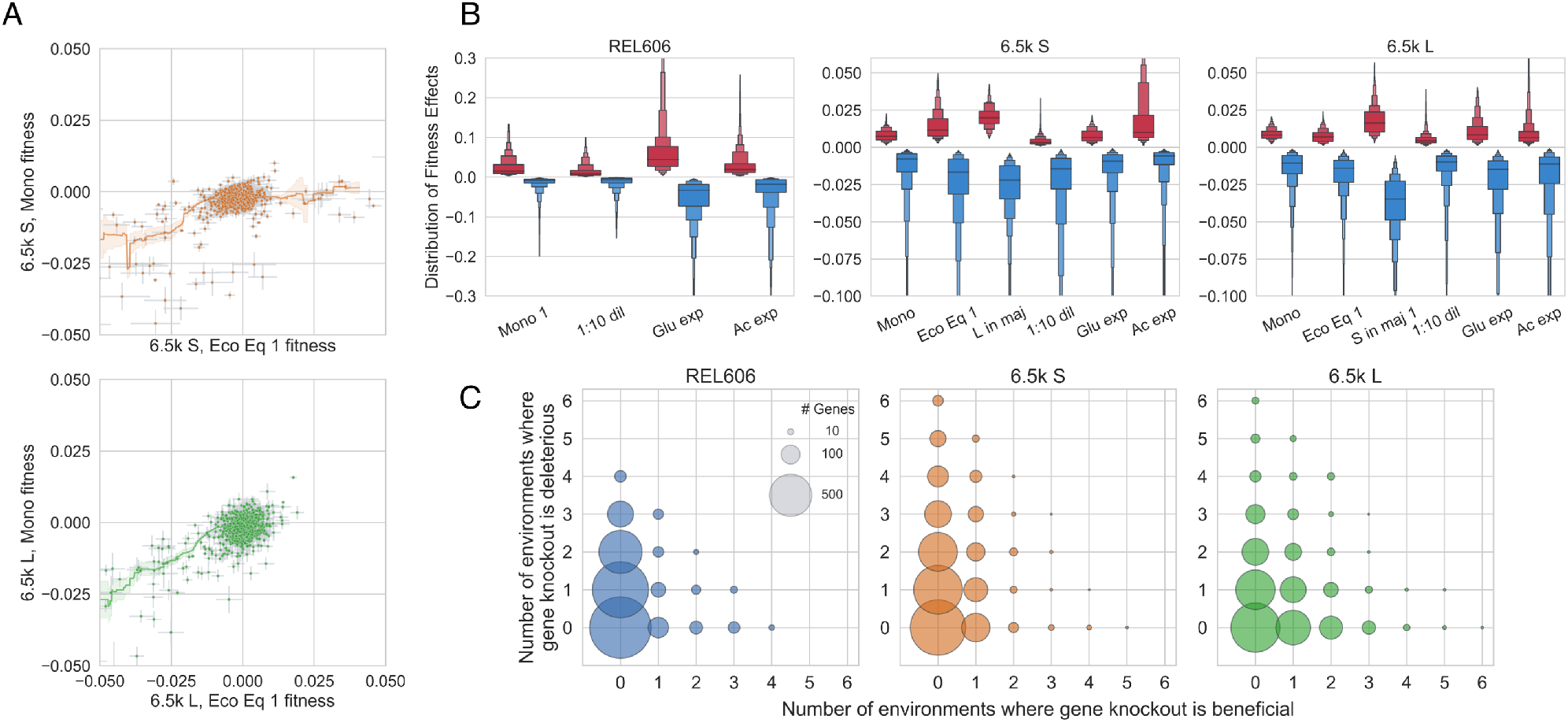
Statistical properties of DFEs as well as effects of individual mutations sensitively depend on environment. (**A**) Knockout fitness effects tend to have a larger magnitude when S and L are at ecological equilibrium versus when they are in monoculture. Line shown is a rolling average of fitness effects ± standard error. (**B**) Distribution of fitness effects across environments, where we only included knockouts that were called as significantly non-neutral. Please note that the DFEs of REL606 are on a different scale than S and L. (**C**) Illustration of how the sign of fitness effects changes across environments. Few mutations are unconditionally beneficial or deleterious, many are non-neutral only in one or few environments, and sign-flipping of fitness effects across environments is pervasive. REL606 only has 4 unique environments, compared to 6 for S and L. Size of circle is proportional to number of genes that fall into each class.

The LTEE environment, while relatively simple, varies quite significantly over the course of a single cycle (43, 54), allowing ecotypes to carve out different temporal ecological niches during cycles of lag, exponential, and stationary phases. To explore how selection pressures vary in different niches in the growth cycle, we measured fitness in exponential growth on glucose and acetate (which appears in the LTEE environment due to overflow metabolism), and at a reduced dilution rate of 1:10 such that portion of the growth cycle in stationary phase is increased (Table 1). We found that the shape of the DFE changed substantially based on the environment (Figures 2B). For example, while S and L have a similar beneficial DFE shape in monoculture, L has access to stronger beneficial knockout mutations in glucose exponential phase compared to S. As another example, the beneficial DFE in acetate is larger than any other DFE in both S and L, potentially pointing to a substantial, as-of-yet unrealized adaptive potential for adaptation on acetate. Interestingly, despite the environmental variation, REL606 always has a more pronounced beneficial DFE compared to S and L.

It is important to note that measurement noise varied non-negligibly across experiments, primarily because of changes in bottleneck size (and thus in the strength of genetic drift) due to differences in library frequency and and other experimental differences (see supplement section S2). Thus, our power to detect selected mutations close to neutrality varied across experiments.

In contrast to previous work (35), it appears that there is no consistent relationship between background fitness and shape of the deleterious DFE, which instead appears to depend more on environment. Possible reasons for the discrepancy include species-dependent differences, and the fact that our set of experiments used backgrounds connected by evolution, while Johnson et al. used evolutionarily unrelated yeast hybrids with varying fitness in the test environment, whose changing DFEs were not controlled by evolution. One possible evolutionary explanation could be second-order selection against mutants with wider deleterious DFEs, because those mutants would be more likely to pick up a deleterious hitch-hiker mutation along with any beneficial driver mutation.

In addition to the strong dependence of the macroscopic DFE on environment, it appears that the fitness effects of individual mutations can also change radically by environment. Strikingly, in the set of considered environments, conditional non-neutrality and sign-flipping appear to occur across all three genetic backgrounds (Figure 2C). The majority of knockouts are non-neutral in at least one measured environment; just about ∼20% of knockouts are called as neutral across all environments. Very few mutations are unconditionally beneficial or deleterious across all environments, and many more mutations flip signs across environments, suggesting the presence of widespread trade-offs between adapting to different components of an environment. The ubiquitous presence of sign-flipping also suggests that subtle changes to environmental conditions–by changes to community composition or niche openness via adaptation– could meaningfully affect evolutionary outcomes by changing which mutations are likely to establish. The presence of sign-flipping still holds if we reduce the p-value cutoff from 0.05 to 10−3 or 10−5 to determine non-neutrality (Figure S12), or only consider genes with |*s* |*>* 1% or |*s*| > 2% as non-neutral (Figure S13), although more genes are called as neutral, as would be expected. However, it is important to note that we only considered genes to be non-neutral if their fitness was significantly different from 0; thus, it is likely that some knockouts were incorrectly called as neutral, especially if their fitness effect is small. Additionally, we have only measured a relatively small set of closely related environments “nearby” the LTEE environment, so we might expect that if we measure fitness in a sufficiently large number of environments, many more genes would be non-neutral in at least one.

By computing the correlation of mutational fitness effects across environments (weighted by measurement error), we can obtain a measure of the functional similarity of environments, which we can also use to cluster said environments (Figure 3A). As a first observation and check, it is reassuring to see the clustering of quasi-replicate experiments, i.e. experiments with relatively minor differences in the experimental set-up and performed on different days–Eco Eq 1/2, S in maj 1/2/3 (L), and Mono 1/2 (REL606) (see supplement section S2). However, the correlations between the quasi-replicates are lower than we see for replicate experiments that we did at the same time–this could indicate either that some fitness measurements are sensitive to the small experimental differences (size of flasks, whether libraries are cocultured or not, etc.), or simply performing the experiments on different days with different environmental fluctuations leads to deviations in measured fitness, as is perhaps the case in other systems (37). The latter hypothesis is further supported by the fact that two experiments were in fact performed at the same time (S in maj 2 and 3), and had among the highest correlation of all quasi-replicates.

**Fig. 3.**
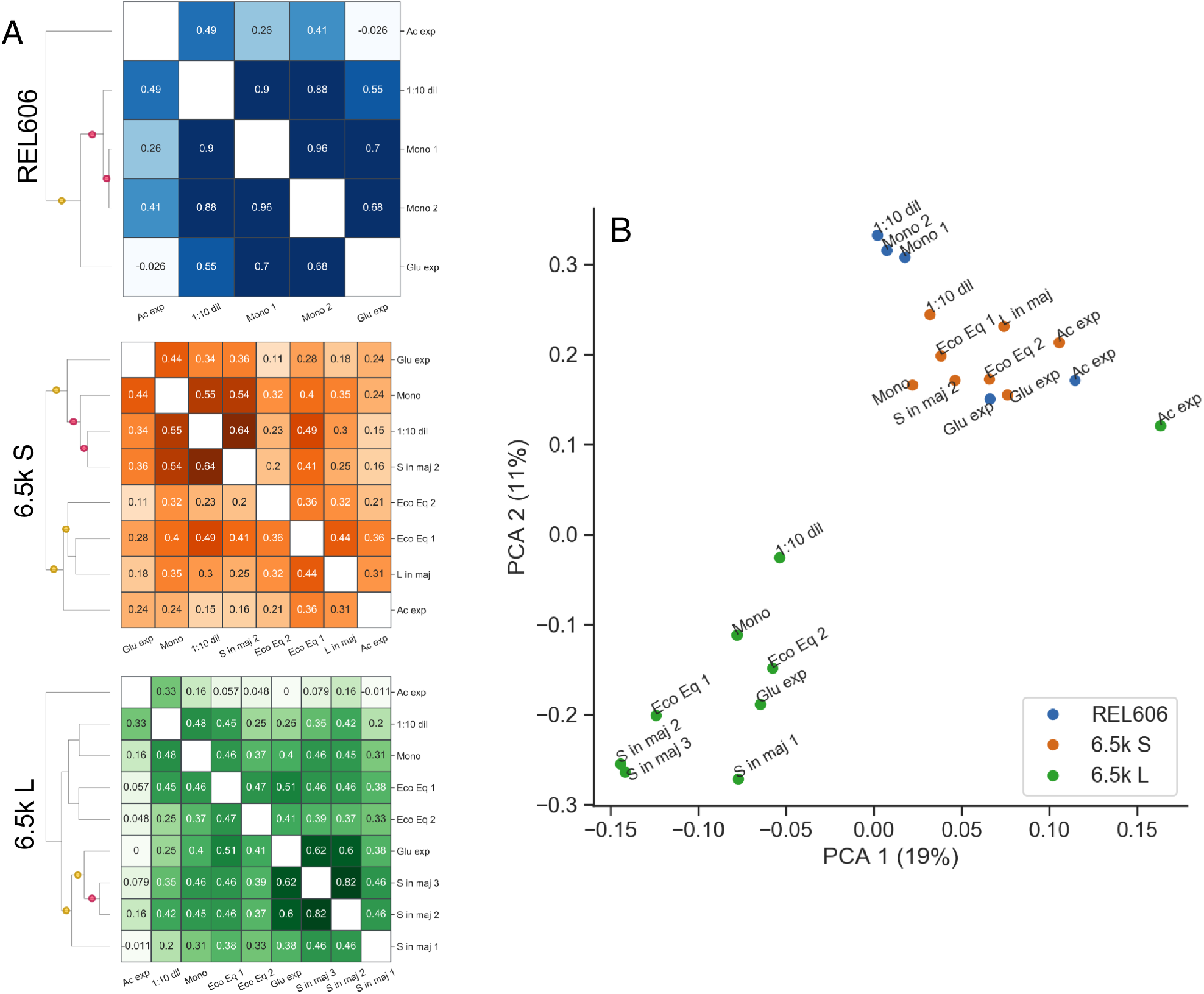
Similarity of fitness effects between environments. (**A**) Clustering environments by using fitness effect correlation as a measure of similarity reveals which environments are the most functionally alike. For example, environments related to the putative ecotype niches–exponential acetate growth and glucose growth, for S and L respectively–cluster with conditions where the ecotype is in the minority. The red and yellow dots indicate that the branch has ≥ 90% or ≥ 70% support respectively, computed via bootstrapping. (**B**) Principal components analysis (PCA) of our data, using (normalized) fitness effects as features (% variance). We see that L experiments cluster separately from the S and REL606 experiments, with the exception of the acetate exponential phase condition.

Otherwise, there are still some interesting patterns that we can pick out by looking at correlations across environments. For example, it looks like the environments related to the putative ecotype niches–glucose and acetate exponential growth in L and S respectively–cluster with conditions where the ecotype is in the minority. On the other hand, the monoculture experiment in S clusters with glucose exponential phase. Also, in REL606 and L, the acetate experiment is the out-group compared to all the other environments, and almost completely uncorrelated with fitness in glucose exponential phase, but most correlated with the 1:10 dilution condition. In S, this is not the case, and acetate fitness is *least* correlated with 1:10 dilution fitness. This may indicate that stationary phase in REL606 and L may have much more acetate with which to grow on compared to S, and adaptation to acetate may involve tradeoffs with adaptation to glucose, at least in REL606 and L. We also performed a principal components analysis on our data, using (normalized) fitness effects as features (Figure 3B). We see that L experiments cluster separately from the S and REL606 experiments, with the exception of the acetate exponential phase condition. Otherwise, the PCA largely reproduces the insights from the previous correlation clustering analysis.

### Correlations between genes across environments

To explore the nature of the strong background dependence that we observed, we sought to understand which genes are correlated with each other across environments, with the intuition that genes that perform the same function should change their fitness effects across environments in similar ways. For example, the *sufABCDSE* operon encodes proteins that help to assemble iron-sulfur clusters (56), and they all have correlated knockout fitness effects across environments in all three genetic backgrounds (Figure S15A)–as they should, if the knockouts all have very similar metabolic/ physiological consequences. However, other gene sets are only correlated in a subset of backgrounds. Most genes in the *fecABCDE* operon are correlated with each other in all backgrounds except for *fecA*, which is well correlated with the others in REL606, less correlated in S, and uncorrelated with the others in L (Figure 4A). Similarly, the genes in the *proVWX* operon are almost perfectly correlated, except one condition where *proV* has a ∼7% higher fitness than the other two knockouts (Figure S15B). We can also look at the fitness effects of a subset of knockouts that are beneficial at least once for every genetic background, across environments (Figure S16). We see that subsets of genes that are sometimes beneficial on a background are positively correlated with each other, e.g. *pykF*/*cyoA* in REL606 and *ptsP*/*mrcA*/*gppA* in 6.5k L, perhaps suggesting that the knockouts have common functional effects. These correlations often break when the mutations appear on different genetic backgrounds, e.g. *pykF*/*cyoA* are no longer correlated on (at least) the 6.5k L background, and *ptsP*/*mrcA*/*gppA* are no longer correlated on the 6.5k S background, while *ptsP*/*mrcA* actually appear *negatively* correlated on the REL606 background. Together, these examples suggest that correlations between knockout fitness effects may change in idiosyncratic ways across genetic backgrounds.

**Fig. 4.**
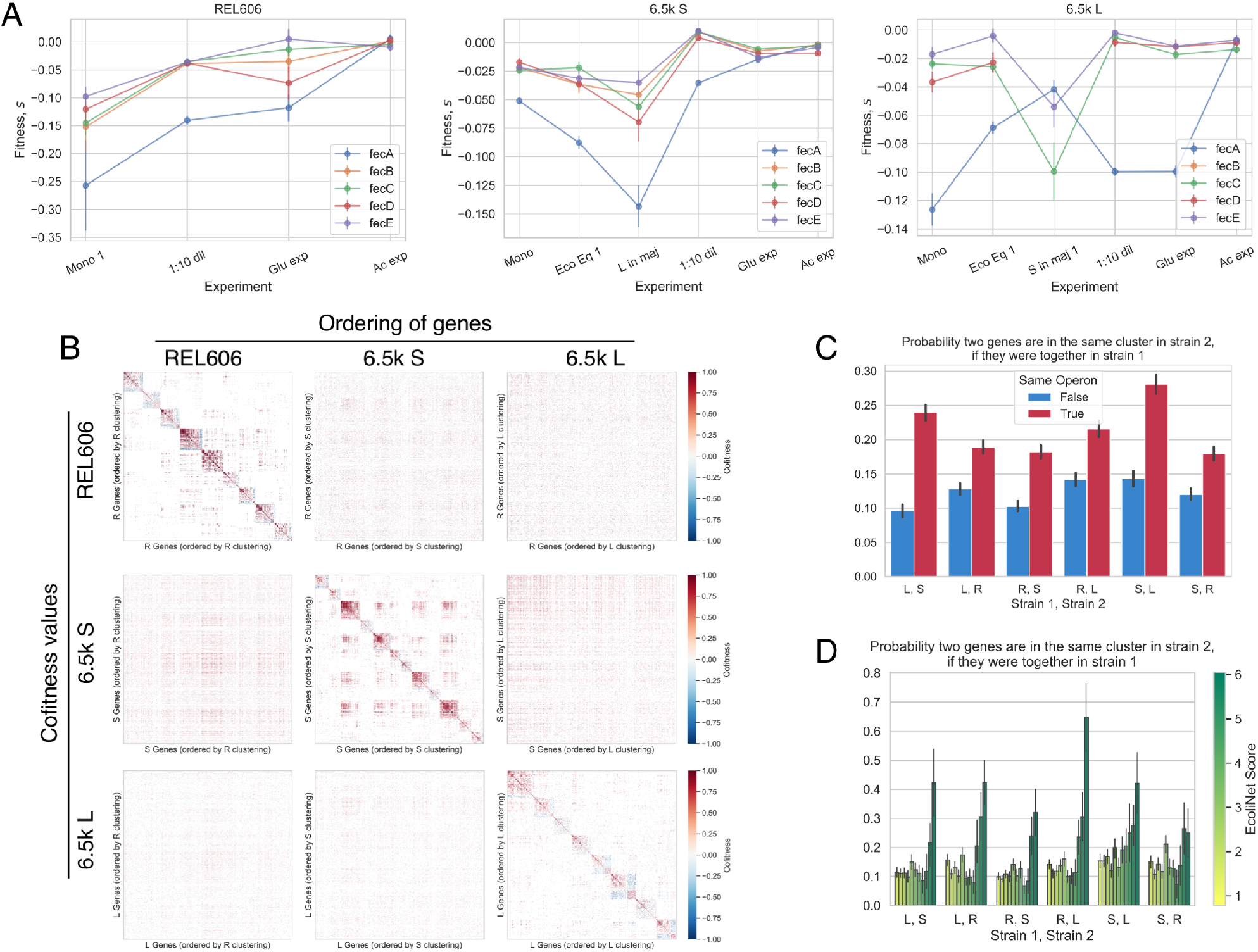
Correlations between genes across environments. We observed that many pairs of genes have correlated fitness effects across environments, for example (**A**) most genes of the *fecABCDE* operon. However, *fecA* is correlated with the other genes to varying degrees, depending on the genetic background. (**B**) We computed the pairwise correlation of fitness effects (cofitness) for all pairs of genes, and then clustered genes with a community detection algorithm (55).We then rearranged the cofitness matrices by reordering genes based on “optimal” clustering of other genetic backgrounds. For each column, we ordered the genes based on the clustering of a given genetic background. For each row, we used the cofitness matrix for a given background. It is apparent that replotting the cofitness matrix using another strain’s clustering does not produce noticeable structure. (**C, D**) Cluster reassortment is not entirely random–pairs of genes (**C**) in the same operon and (**D**) that strongly interact with each other (high EcoliNet score), tend to stay in the same clusters across genetic backgrounds. In contrast, the cofitness of pairs of genes that are not in those categories appear to change in a way that is indistinguishable from random reassortment. In panels C and D, the abbreviations R, S, and L refer to REL606, and 6.5k S/L, respectively.

We systematically quantified the pairwise correlation of knockout fitness across environments–termed “cofitness”, previously defined in (41)–where we used the weighted pearson’s correlation coefficient to account for differences in measurement error across environments. We computed the cofitness of all pairs of genes (excluding those called as neutral across all environments) across the REL606, S and L libraries, as well as a null cofitness distribution for each pair to determine if the two genes are significantly correlated; the set of all significant gene-gene correlations determine the edges in the cofitness networks (see supplement section S4.2). We explored the structure of the resulting cofitness networks via clustering (55) (see supplement section S4.2), where we found sets of communities for all three libraries with *modularity >* 0, indicating that there are more edges within each community than between communities (Figure 4B) (57). We performed a number of controls to ensure that our results weren’t driven by measurement noise or technical effects of clustering; see supplement section S4.2 for more information.

The presence of strong communities suggests that most knockouts are significantly correlated with others, potentially pointing to similar functional effects driving changes in fitness. We then wanted to compare how these clusters differ between the different genetic backgrounds, with the idea that how and if clusters change should reveal information on how the effective functions of genes differ across genetic back-grounds. Surprisingly, we find that gene clusters are not well preserved across genetic backgrounds, and in fact, genes are typically seemingly randomly reassorted between genetic backgrounds (Figures 4B, S21). In further support of correlations breaking between backgrounds, if we recompute the cofitness networks using only one of the biological replicates per experiment, we see that cofitness networks are more similar within genetic backgrounds compared to between backgrounds (Figure S19). There are a couple clusters that show non-random sampling across genetic background, however, the deviation from random sampling is mostly small, with one noticeable exception–clusters 5, 3, and 1 in REL606, S, and L, respectively, all seem to share a larger than random number of genes with each other (*p <* 10−4 for all clusters). From a Gene Ontology enrichment analysis, genes that are associated with biofilm formation (GO:0043708), adhesion (GO:0022610), and pilus organization (GO:0043711) are over-represented in these clusters, along with genes involved in organonitrogen compound biosynthesis (GO:1901566), although to a weaker extent (Figure S22). This suggests that there is at least one (large) functionally related group of genes that stay correlated across genetic backgrounds, implying that their fitness-determining effects are mostly the same, regardless of genetic background.

We wanted to know why other functional groups of genes do not stay correlated with each other, and if there was any structure hiding in the seeming randomness of cluster reassortment. A simple first test could ask if genes in the same operon are more likely to stay correlated with each other across backgrounds, which is the case for several of our aforementioned examples. This indeed appears to be the case across all genetic backgrounds (Figure 4C). However, genes often share functions with other genes outside their operons, so we turned to investigating the relationship between the cofitness and genetic networks. We used EcoliNet as a representation of the *E. coli* genetic network, as it attempts to capture all interactions between genes by integrating various data-types, regardless of the mechanism (transcriptional, protein-protein, etc), and assigns a score to each interaction that effectively represents the strength of the interaction (58). We then computed the probability that two genes are in the same community in one genetic background, given that they’re together in another background, as a function of EcoliNet score (Figures 4D). We see that gene pairs that are predicted to strongly interact (high EcoliNet score) are much more likely to be correlated across genetic backgrounds. We can also see these same patterns without referencing any cluster labels–if we look at the correlation between all cofitness pairs across genetic backgrounds, pairs that are in the same operon (Figure S23A) and those with the highest EcoliNet score (Figure S23B) give the highest correlation. It also appears that the shortest distance between two nodes in the EcoliNet network (Figure S25) also predicts if the two genes will stay correlated across genetic background, albeit the effect is weaker. We should note that it is perhaps the case that there are weaker consistencies across backgrounds for non-operon/non-interacting genes pairs that we don’t have the statistical power to detect. Still, these analyses suggest that evolution significantly changes which functional effects of genes are important for determining fitness, such that the cofitness of genes pairs is much more preserved across genetic background for the most strongly interacting genes, but not as much for other gene pairs.

### Fitness effects are correlated with evolutionary outcomes

We sought to explore if the knockout fitness effects that we measured were correlated with evolutionary outcomes in the LTEE, i.e. establishment of mutations and changes in gene expression. So, we first investigated if genes with non-neutral knockout fitness were more or less likely to be mutated and rise to a sufficiently high frequency in the population. Using the clonal sequencing data from Tenaillon et al. (2016) (59) and Plucain et al. (2014) (46), we identified genes that mutated between selected LTEE time-points, and ran a logistic model with fitness effect as the predictor and mutated status as the response variable (see supplement section S4.3), separately for beneficial (Figures 5A) and deleterious genes (S26A). We used three sequenced clones (one available for each time point) for both S and L, while we used all clones from all non-mutator populations (at a given time point) for REL606. We used the appearance of a mutation (excluding synonymous SNPs) within a gene as a proxy for establishment.

**Fig. 5.**
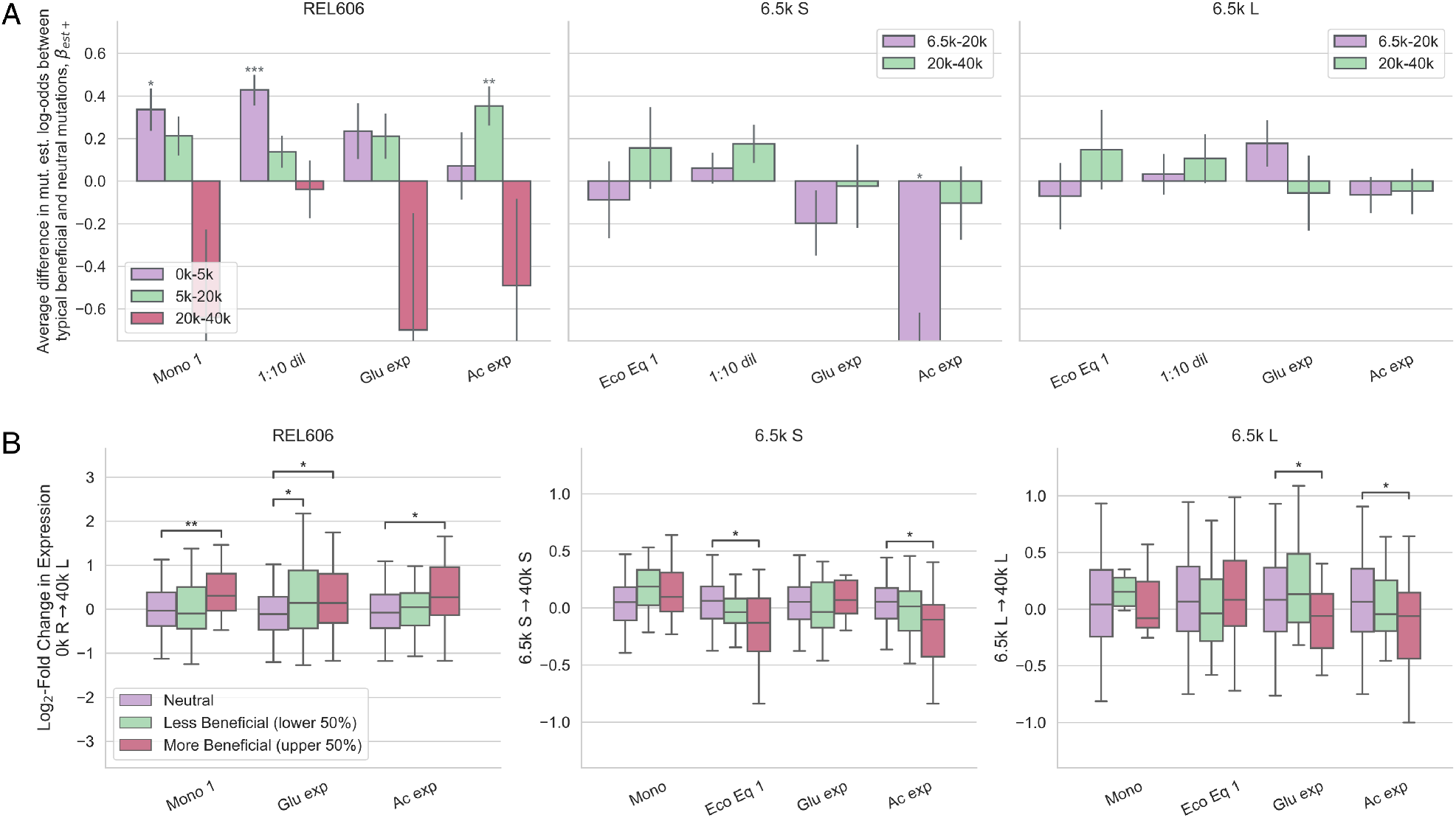
Fitness effects of beneficial genes are correlated with evolutionary outcomes. We explored if genes with beneficial knockout fitness effects are correlated with (**A**) establishment of a mutation in a gene, and (**B**) changes in gene expression over evolutionary time, relative to neutral knockouts. (**A**) Slopes from logistic models, with presence of a mutation in a gene as the response variable. The fitness effects were normalized by the median beneficial fitness effect, so that coefficients can be interpreted as the average difference in log-odds establishment between neutral knockouts and the ‘typical’ beneficial knockout. REL606 beneficial knockout fitness is positively correlated with gene establishment probability for most environments, but in different time intervals, potentially pointing to shifting targets of selection. (**B**) We compared the distributions of log-fold change in expression between genes with neutral knockout fitness effects, less beneficial effects (lower 50%), and more beneficial effects (upper 50%). We used the change in expression from 0k gens (REL606) to 40k gens (L), from 6.5k gens (S) to 40k gens (S), and from 6.5k gens (L) to 40k gens (L) for the REL606, 6.5k S, and 6.5k L panels, respectively. The expression change between ancestor and 40k L (left) is nearly identical to the expression change between ancestor and 40k S as well as other timepoints (Figure S29). Beneficial knockout fitness in REL606 is generally positively correlated with increasing gene expression over time. In S and L, fitness in several environments–including the ecological equilibrium and acetate and glucose growth–is correlated with decreasing gene expression. Asterisks denote coefficients/comparisons that are significantly different from 0 (FDR correction; * *p <* 0.05, ** *p <* 0.01, *** *p <* 0.001).

Fitness of beneficial knockouts in the 1:10 dilution condition and monoculture (LTEE condition) in the REL606 background is strongly correlated with which mutations establish from 0-5k generations, while fitness in acetate exponential phase is only correlated with establishment later in the evolution (difference in slopes between 0-5k and 5-20k is significant at *p <* 0.05 via permutation test for 1:10 dilution and acetate conditions, not for monoculture or glucose conditions). This is potentially a signal that the targets of selection are shifting over time–REL606 may initially adapt via lag phase shortening/stationary phase survival, while only later adapting via increased acetate growth rate. This could happen, for example, by either clonal interference favoring the highest-effect mutations, or due to global epistatic effects (47). The former hypothesis is supported by the observation that three mutations appear in genes with beneficial acetate knockouts at 2k generations, but they then disappeared by 5k generations, potentially indicating that they were out-competed by other beneficial mutations (Figure S28). There is only one S/L condition that shows a significant difference in mutation establishment probability between beneficial and neutral mutations–genes with beneficial fitness in acetate are less likely to mutate compared to neutrals in S. However, changes in gene expression suggest adaptation to acetate may be occuring through indirect routes in S, as detailed below. However, we expect our power to detect correlations between mutational fitness and mutation establishment to be lower for S and L. They have a ∼100x higher mutation rate than REL606 (48), implying that the ratio of neutral hitchhiking to beneficial driver mutations is higher as well.

We also investigated if fitness effects are correlated with changes in gene expression, using microarray data from Le Gac et al. 2012 (16), which measured gene expression in REL606, and S/L at 6.5k, 17k, and 40k generations. These measurements serve as a distinct readout of evolutionary change compared to genomic mutational dynamics, because even if a gene is not directly mutated, gene expression can still change through indirect genetic interactions. Thus, gene expression measurements allow us to probe the effects of the cumulative mutations fixed by evolution. We compared the distribution of log-fold expression changes over approximately 40k generations for genes with neutral and non-neutral knockout fitness effects, separately for beneficial (Figure 5B) and deleterious genes (Figure S26B). We see that the median change in gene expression is significantly different between neutral and beneficial genes across several conditions, but generally only for the upper 50% of beneficial genes. This indicates that the magnitude of the knockout fitness effect is important for determining how much the median gene changes in expression. We can get more power to detect relationships between the magnitude of the knockout fitness effect and log-fold change in gene expression by fitting linear models to the data (Figure S29). The same patterns hold if we restrict our analysis to highly expressed genes (Figure S30).

In REL606, genes with beneficial knockout fitness effects tend to increase in expression (relative to neutral genes) over evolutionary time; this is perhaps surprising, because we would expect selection to decrease gene expression if knocking out that gene is beneficial. We saw the same pattern with deleterious genes (Figure S26B). One possibility to explain tvery, the expression of growth-relevant genes is increased by some mutation with a highly pleiotropic effect (e.g. in a master regulator), whose overall benefits outweigh the costs of raising the expression of beneficial knockout genes.

In contrast, in S and L, there are a couple of environments where gene expression significantly decreases over evolutionary time for genes with beneficial knockout fitness effects (compared to neutrals). These conditions include environments related to the putative ecotype niches–acetate and glucose exponential growth in S and L respectively. On the other hand, while fitness in the ecological equilibrium is associated with decreased gene expression, this is not the case for fitness in monoculture and the 1:10 dilution environments, indicating again that the latter environments are less relevant for evolution in the LTEE environment. Despite the fact that acetate-adapting mutations are not establishing on the S background (at least initially), gene expression still decreases by 40k generations, perhaps indicating that adaptation to acetate is occurring through routes other than directly mutating genes with beneficial knockout effects.

We also saw that S and L beneficial knockout fitness in glucose exponential phase is *positively* correlated with an increase in gene expression from 0-6.5k (Figure S29). On average, those same genes decrease in relative gene expression when evolving on the L background, whereas they do not change on the S background. This set of data could indicate that from 0-6.5k many genes increased in gene expression via adaptive evolution that were actively unhelpful for glucose growth, either because of transcriptomic misallocation or other types of antagonistic pleiotropy, such that knocking them out conferred a benefit. Upon diversification of S and L, the direction of gene expression change appears to switch for L, perhaps suggesting that L is evolving towards a more glucose growth-optimized transcriptome, while S is not. This set of observations provides a possible example of how diversification changes the selection pressures acting on organisms.

Interestingly, deleterious knockout fitness effects across all environments in S/L tend to be associated with an increase in gene expression between 0 and 6.5k generations (Figure S26B). This observation may provide a partial explanation for why some knockouts become deleterious in S/L when they were neutral in REL606–6.5k generations of evolution caused the genes to suddenly become important, so they became more costly to knock out. Another, unrelated observation could help us to understand why some genes have deleterious knockout fitness effects–it appears that deleterious genes are more highly connected in the *E. coli* gene interaction network (EcoliNet) compared to neutrals (on average), indicating that some genes may be deleterious because when they’re knockout out, they also affect the functioning of many other genes (Figure S31).

## Discussion

In order to be able to predict how evolution will proceed in community contexts, we need to know the distribution of mutational fitness effects, along with how it depends on genetic background and ecological conditions. To that end, we measured the genome-wide knockout fitness effects of a recently diversified ecosystem, S and L, and their ancestor, REL606. Despite the fact that the fitness effects of individual mutations appear to be highly dependent on both genetic background and environment (strong (G ×) G × E effects), we saw consistent statistical patterns of variation across both axes, namely global diminishing returns epistasis and a negative frequency-fitness correlation (in S and L). In contrast, previous studies that observed diminishing returns epistasis saw both the mean of the DFE as well as the fitness effects of individual mutations decrease as a function of background fitness (51, 52); this discrepancy may indicate that uniform negative epistasis of individual mutations may only be relevant for the first handful of mutational steps, before yielding to more complex and idiosyncratic forms of epistasis. While the underlying mechanism that generates this form of global epistasis is still unclear, our observations are consistent with recent theoretical (60) and experimental work (61) that suggest that global diminishing returns epistasis may arise as a general consequence of idiosyncratic epistasis.

Even though S and L only diverged ∼500 generations ago, the mixing ratio of the two ecotypes strongly affects the DFEs, suggesting that strong eco-evolutionary coupling is possible even in closely related strains. This would imply that selective pressures depend strongly on the community mixture, which changes significantly and relatively rapidly due to evolution (42, 47). The sensitivity of knockout fitness effects to relatively minor variations on the LTEE environment, such as changing niche availability or ecosystem composition, may be evolutionarily significant–we know that the growth traits of S and L also change quite drastically during their coevolution (16, 44), which along with changes to ecosystem composition, will change the environment, and thus change which mutations are favored by selection. One specific hypothesis that emerges from our data is that selective forces may be more similar to environments related to the putative ecotype niches when the ecotype is rare, for both S and L. This is supported by both clustering environments by fitness effect correlations, and which environments were correlated with changes in gene expression. It would follow that selection could favor different degrees of specialization within the current niche as the ecotype frequencies and growth traits change due to evolution. Regardless of the specific implementation, the process where (i) mutations change growth traits and ecosystem composition, which (ii) change ecological conditions, which in turn (iii) change the mutational fitness effects of both ecotypes, could represent an important and pervasive type of eco-evolutionary feedback.

We aimed to better understand the background and environment dependence of mutational fitness effects by systematically studying fitness correlations across environments. Our intuition was that knocking out genes with similar functions should have similar effects across environments. We saw that, by and large, different sets of genes were correlated with each other across genetic backgrounds; only strongly interacting pairs of genes were likely to be correlated across all backgrounds. These widespread changes could be caused by a number of different evolutionary phenomena–for example, evolution could have induced widespread changes in the functional effects of genes or which functional effects matter for fitness. Additionally, inasmuch as fitness in an environment is a reflection of phenotype–e.g. fitness in exponential phase is likely a simple function of exponential growth rate– the extensive changes in fitness across environments could be interpreted as support for ubiquitous pleiotropic effects of knockout mutations.

We investigated if our measured knockout fitness effects were correlated with evolutionary outcomes, i.e. mutation establishment and gene expression changes. We found significant correlations across several, but not all environments, leading to hypotheses on how selection has acted on LTEE populations. From correlations of knockout fitness effects with mutation establishment, we found potential signals of shifting selection over time in REL606. Changes in gene expression provide a distinct window into evolutionary change, as expression can change through genetic interactions, even if a gene is not directly mutated. Among other patterns, the fitness correlations with gene expression changes potentially reveal how the traits under selection changed from preto post-diversification, and how they are different between S and L. Pinpointing the precise causes of these patterns could be a fruitful avenue for future work. Overall, the connections between evolutionary changes and knockout fitness effects demonstrates the utility of our approach to understand how adaptation happens in the “natural” evolutionary context.

Ultimately, we would like to predict the outcomes of evolution in community contexts. By showing how the distribution of invasion fitness effects changes as a result of genetic background and ecological conditions, our dataset represents a major step forward in that direction. The invasion fitness effects directly impact the establishment probability of a beneficial mutation, as well as the mutant dynamics until it reaches a substantial proportion of the population. The distribution of deleterious invasion fitness effects also controls other relevant evolutionary phenomena, including the equilibrium reached by mutation-selection balance, and the probability that a deleterious mutation will hitchhike on a beneficial mutant (“genetic draft”). However, in principal, the fitness effect of a mutation could change as it approaches fixation (within the ecotype) due to frequency-dependent effects. We are not able to measure these effects with our experimental set-up, as our ability to measure fitness effects in high-throughput requires that mutants remain rare. However, frequency-dependent mutations could significantly alter expected evolutionary dynamics, so as such, measuring such effects are a major direction for future work.

As previously mentioned, we only surveyed the fitness effects of knockout mutations, which represent a subset of all mutations available to an organism. While it is possible that other types of mutations could display different patterns, knockout mutations appear to be prevalent and important for adaptation in the LTEE (47, 62), and our measured knockout fitness effects are correlated with evolutionary outcomes. Additionally, we studied a relatively simple ecosystem, consisting of just two recently diverged ecotypes; measuring the mutational effects in more complicated ecosystems and how they change as a result of longer periods of evolution is likely a fruitful future avenue of investigation. Overall, the methods and results presented here pave the way for future studies investigating how mutational fitness effects depend on ecoevolutionary processes, and how eco-evolutionary feedback arises from changing fitness effects.

## Methods

See supplementary information.

### Data, code, and strain availability

Glycerol stock copies of the REL606, 6.5k S, and 6.5k L Tn5 barcoded libraries are available upon request. All code used to process the data and perform the analyses as well as processed data are available on GitHub, https://github.com/joaoascensao/S-L-REL606-BarSeq

## ACKNOWLEDGEMENTS

We thank Matti Gralka and QinQin Yu for early-phase experimental assistance. We thank current and former members of the Hallatschek and Arkin labs for productive discussions and feedback regarding this project, especially Morgan Price, Jonas Denk, and QinQin Yu. We thank Richard Lenski for sending us the REL606, 6.5k S and L clones, along with experimental advice and feedback. Research reported in this publication was supported by a National Science Foundation CAREER Award (1555330) and by the Miller Institute for Basic Research in Science, University of California, Berkeley. Some elements of this project were funded by ENIGMA-Ecosystems and Networks Integrated with Genes and Molecular Assemblies (http://enigma.lbl.gov), a Science Focus Area Program at Lawrence Berkeley National Laboratory, and is based upon work supported by the U.S. Department of Energy, Office of Science, Office of Biological & Environmental Research under contract number DE-AC02-05CH11231. JAA acknowledges support from an NSF graduate research fellowship, a Berkeley fellowship (from UC Berkeley), and Lloyd and Brodie scholarships (from UC Berkeley Dept of Bioengineering). BHG acknowledges support from an award from the Alfred P. Sloan Foundation (FG-2021-15708). This research used resources of the National Energy Research Scientific Computing Center (NERSC), a U.S. Department of Energy Office of Science User Facility located at Lawrence Berkeley National Laboratory, operated under Contract No. DE-AC02-05CH11231 using NERSC award ERCAP0021398. This work used the Vincent J. Coates Genomics Sequencing Laboratory at UC Berkeley, supported by NIH S10 OD018174 Instrumentation Grant. The bacteria (IGI prokayote icon; https://innovativegenomics.org/glossary/) and erlenmeyer flask (flask-2 icon by DBCLS; https://bioicons.com/) clipart from Figure 1B are used and modified under creative commons licenses, CC BY-NC-SA 4.0 and CC-BY 4.0 respectively. BioRxiv preprint was prepared in Overleaf with the (modified) Henriques Lab LaTeX template.

## Supplement

### S1 Barcoded transposon library construction

To construct the barcoded transposon libraries, we isolated subclones of REL606, REL11555 (6.5k S), and REL11556 (6.5k L), all gifts of Richard Lenski (Michigan State University). Transposon mutagenesis was performed as previously described (40, 41) by mating each LTEE clone with an *E. coli* WM3064 donor (Diaminopimelic acid [DAP] auxotroph and pir^+^) containing previously described (40) randomly barcoded Tn5 plasmids with a kanamycin cassette and an R6K origin of replication. The LTEE clones were grown in DM2000 (Davis Minimal Media with 2000mg/L Dglucose), and the donor was grown in LB/Kan, all to mid-log phase. After washing the cultures, each LTEE culture was then mixed with the donor in a 1:1 ratio, then placed on 0.45 µM nitrocellulose filters (Millipore cat. no. HAWP04700) on top of a 1% agar plate with EZ-MOPS rich, defined media (Teknova cat. no. M2105) + 20mM sodium pyruvate (’EZ-py’) + 0.3mM DAP. The rich media was chosen because it had a number of different carbon sources (glucose, amino acids, pyruvate) and sufficient amounts of all other required macro/micronutrients, lessening the chances of substantial negative selection in the growth media. After conjugation, the filters were picked up and placed in rich media; subsequently, the resuspended cells were plated on EZ-py agar plates supplemented with 50 µg/mL kanamycin. After approximately 24hrs of growth at 37C, colonies were scraped up and grown in EZ-py liquid media with 50 µg/mL kanamycin until OD∼1; we then saved the cultures in several 10% glycerol stocks. Transposon insertion mapping (TnSeq) libaries were prepared as previously described (40); libraries were then sequenced on the Illumina HiSeq 4000 (150PE) at the Vincent J. Coates Genomics Sequencing Laboratory at UC Berkeley. The resulting sequencing data was used to create a table relating each barcode to a genomic insertion location, using a previously developed script (MapTnSeq.pl) (40).

### S2 BarSeq experiments

#### S2.1 Set-up of experiments

To start a BarSeq experiment, we first unfroze 1mL glycerol stock of the REL606, 6.5k S and/or 6.5k L transposon libraries and transferred the entirety to 10mL EZ-py media (media used for library construction) in 50mL glass erlenmeyer flasks, which were grown for 16-24hrs at 37C, shaken at 120rpm. All cultures for all experiments were grown with the same shaker, in the same 37C warm room. In several experiments where we measured fitness effects of 6.5k S/L barcoded libraries at various ecotype frequencies, we also grew the wild type S/L with the same media, under the same conditions. The next day, we washed the cultures by pelleting via centrifugation for 3 minutes at 5000rpm, aspirating the supernatant, and resuspending in DM0 (Davis Minimal Media without a carbon source) three times. After thoroughly vortexing the cultures, we transferred them 1:1000 to the appropriate media in *n* flasks (see below)–depending on the experiment, we used different numbers of flasks and different sizes, either 10mL media in 50mL glass flasks or 200mL media in 1L glass flasks (same ratios, scaled up). We used multiple flasks and larger flasks to increase the total population size, decreasing fluctuations due to genetic drift. We then performed two more transfers in the appropriate conditions for the experiment to help physiologically adapt the cultures to the conditions. If we were doing a coculture experiment, we would mix the cultures at the appropriate frequencies during the second transfer. If we used multiple flasks in an experiment, we would sample an equal amount of culture from each flask into a microcentrifuge or Falcon tube, thoroughly mix the cultures, and redistribute among the same number of flasks with new media–thus, the cultures distributed in multiple flasks were effectively all part of the same population. After the third transfer, we would collect cells for day 0 of the experiment, and use that culture to start two biological replicates that are independently propagated for the remainder of the experiment. All cultures were grown at 37C, shaken at 120rpm. Cells were harvested at defined time points by centrifugation at 15000rpm for 10min of ∼60mL culture for all experiments except Ac Exp (10mL) and Mono 2 (30mL), pooling culture from all flasks in an experiment/replicate at equal ratios. Subsequently, the pellets were stored at -80C until the experiment was finished.

#### S2.2 Conditions for each experiment

##### S2.2.1 Monoculture

For the Mono (1) experiments, we propagated the libraries alone in DM25 (Davis Minimal Media with 25mg/L D-glucose) in 5x 50mL flasks over the course of 4 days. For the REL606 Mono 2 experiment, we used 3x 50mL flasks over the course of 8 days, with four biological replicates in DM25. We transferred cultures 1:100 every 24hrs, and took the number of generations per transfer as log_2_ 100.

##### S2.2.2 Coculture experiments

As mentioned above, we started wildtype cultures of 6.5k S and/or L clones (same clones used to make the RB-Tn libraries) at the same time and with the same procedure as the library cultures (Table 1, main text), and mixing the cultures at the appropriate frequencies at the second “adaptation” serial transfer. We measured the ecological equilibrium frequency to be approximately 15 − 20% S (Figure S8), so we ensured that the S frequency was started in that range for the “ecological equilibrium” experiments. We started the “S/L in majority” experiments such that the minority ecotype was > 10% of the total population (Figure S7).

We used DM25 media and propagated the cultures for 4 days, except for S in maj 2/3 where we used 6 days, transferring 1:100 every 24hrs (log_2_ 100 generations) for all coculture experiments. For the Eco Eq 1 experiment, we mixed both S and L libraries in the same cultures along with wildtype L, using 4x 1L flasks. For the Eco Eq 2 experiments, S and L libraries were in separate cultures, both with wildtype S and L set at the appropriate frequency, with RB-Tn library frequency around 5 − 10% (Figure S7); cultures were propagated in 10x 50mL flasks. For the L in Maj and S in Maj 1 experiments, we mixed wt L + S library and wt S + L library, respectively; cultures were propagated in 10x 50mL flasks. For the S in maj 2/3 experiments, we mixed wt S with S+L and L libraries respectively; cultures were propagated in 4x 1L flasks.

We measured the frequency of S/L in the population by plating and counting colonies at the end of a transfer on TM plates (tetrazolium maltose; 10g/L tryptone [Sigma T7293], 1g/L yeast extract [Sigma Y1625], 5g/L NaCl, 16g/L agar, 10g/L maltose, 1mL/L 5% TTC [Sigma T8877]), where S appears as red colonies and L appears as white colonies, previously used in (46). We could also measure the frequency of cells from RB-Tn libraries by plating the cultures on LB/Kanamycin plates, as the transposon has a kanamycin resistance cassette (Figure S7). We diluted all cultures (at the end of a cycle) in DM0. Dilution rates varied over experiments: in Eco Eq 1, we diluted cultures by a factor of 2 ∗ 10−5 *mL*^−1^ to plate on both TM and LB/Kan plates, in Eco Eq 2 we used dilution rates of 10−5 *mL*^−1^ and 10−4 *mL*^−1^ to plate on TM and LB/Kan plates respectively, in the L in Maj and S in Maj 1 experiments we used a 2 ∗ 10−5 *mL*^−1^ dilution rate to plate on just TM plates, and in the S in maj 2/3 experiments we used dilution rates of 2 ∗ 10−5 *mL*^−1^ and 2 ∗ 10−4 *mL*^−1^ to plate on TM and LB/Kan plates respectively.

##### S2.2.3 1:10 dilution

We propagated cultures with a 1:10 dilution, instead of the standard LTEE dilution rate of 1:100, to investigate the effect of a lengthened stationary phase relative to exponential phase. We used DM27.8 media (Davis Minimal Media with 27.8mg/L D-glucose), because the concentration of glucose would fall to 25mg/L after dilution. We used 1x 1L flask for each library culture (180mL media + 20mL culture), propagating the cultures for 8 days every 24hrs with log_2_ 10 generations per day. We pelleted and saved cultures every other day (0,2,4,6,8).

##### S2.2.4 Acetate exponential phase

We sought to measure knockout fitness effects when the RB-Tn libraries were kept in acetate exponential phase, where we used DM2000-acetate (Davis Minimal Media with 2000mg/L Sodium Acetate) and grew the cultures in 1x 50mL flask. We first measured exponential growth rates for wt REL606, L, and S clones in DM2000-acetate, which were approximately 0.08/hr, 0.12/hr, and 0.18/hr respectively. We also observed that all cultures were still in mid-exponential phase at OD∼0.6. So, if we started at initial OD_0_ of 0.09, 0.03, 0.008 for REL606, L, and S respectively, the cultures would end up at OD∼0.6 after 24 hours. Thus, for each transfer, we would measure the actual OD for each culture (after 24hrs of growth) and transfer the appropriate volume of old culture to new 10mL DM2000acetate such that the final concentration was the appropriate OD_0_. We recorded the number of generations for each cycle as log_2_ *OD*_*f*_ /*OD*_0_. Due to the variable number of generations per transfer for each genetic background (owing to different growth rates), we collected samples at days 0,2,4,6,8 for REL606; 0,1,2,4,5,6 for L; 0,1,2,3,4,5 for S.

##### S2.2.5 Glucose exponential phase

We measured knockout fitness effects in glucose exponential phase with DM25 media in 1x 1L flask. We measured the length of DM25 exponential phase to be about 8.25 hrs for REL606, and 5.25 hrs for both S and L after a 1:100 dilution into new media. For the adaptation phase, we did two full 24hr cycles of growth in DM25, followed by one cycle of growth for ∼8hrs and ∼5hrs for REL606 and S/L, respectively. After the adaptation phase, we transferred cultures 1:100 into new DM25 media (warmed to 37C) four times, after 7.5-8hrs for REL606 and 4.5-5hrs for S and L. As DM25 media is quite dilute and thus OD measurements are relatively inaccurate, we estimated the number of cells that were transferred by plating the cultures on LB plates at a 2 ∗ 10−5 *mL*^−1^ dilution rate and counting colonies, calculating the number of generations for that transfer as log_2_ 100 *CFU*_*f*_ */CFU*_0_. We only ended up including the first two transfers of the REL606 library experiment (time points 0,1,2), as it was apparent from CFUs that the third transfer resulted in a large bottleneck owing to a smaller than expected population size before the transfer, likely because of slower than expected growth.

### S2.3 DNA extraction, PCR, Sequencing3

After the experiment was finished, pellets were pulled from the -80C freezer and genomic DNA was extracted with the Qiagen DNeasy tissue and blood extraction kit (cat no. 69504), eluted in double distilled water with typical yields around 50ng/µL. DNA barcodes were amplified from gDNA samples via PCR with Q5 Hot Start Polymerase (NEB, cat. no. M0493S); 50ul reactions were composed of 5µL PCR primers, 5µL gDNA, 10µL 5x buffer, 10µL GC enhancer, 1µL dNTPs, 0.5µL Q5 polymerase, 18.5µL water. We used custom dual-indexed primers that contained binding sites up- and down-stream of the barcode region, along with the necessary Illumina read/index binding sites; fwd primer (AATGAT ACGGCG ACCACC GAGATC TACACT CTTTCC CTACAC GACGCT CTTCCG ATCT N_*n*_XXXXXX GTCGAC CTGCAG CGTACG) where X stands for the custom forward 6bp index, and N_*n*_ is 1-4 random nucleotides, varying with the primer pair; rev primer (CAAGCA GAAGAC GGCATA CGAGAT XXXXXX GTGACT GGAGTT CAGACG TGTGCT CTTCCG ATCTGA TGTCCA CGAGGT CTCT) where X stands for standard Illumina 6bp IT index. We used a different primer pair for each gDNA sample from a different experiment/replicate/time point, so that we could demultiplex the samples after sequencing. The PCR program was 4min at 95C, [30sec at 95C, 30sec at 55C, 30sec at 72C] x25 cycles, 5min at 72C. We verified that we had the correct PCR products via agarose gel electophoresis. All PCR reactions were then pooled and cleaned with the Zymo DNA Clean and Concentrator kit (cat. no. D4013), and eluted in double distilled water. The final pooled sample was then sequenced on an Illumina HiSeq 4000 (50SR) at the Vincent J. Coates Genomics Sequencing Laboratory at UC Berkeley.

## S3 Fitness inference pipeline

### S3.1 Read counting and error correction

We first processed the raw (demultiplexed) sequencing reads using a previously developed Perl script (40, 41) that pulls out the barcode sequence by trimming regions corresponding to the sequencing primers and regions up/downstream of the barcode, as well as discarding reads that do not match the secondary sequencing index or have insufficiently high quality scores (MultiCodes.pl). Then, counts of unique barcodes are tabulated to get a table corresponding barcode sequence to counts. However, due to errors that arise during PCR and sequencing, some of the barcode reads acquire mutations that would prevent them from directly mapping to a transposon insertion location. Thus, we must correct for these sequencing errors by matching mutated barcodes to their parent, and merging the read counts together. The aforementioned Perl script identifies off-by-one barcode pairs; if the minority barcode (the one with fewer counts) unambiguously maps to a single majority barcode, the barcode counts are merged. To detect larger mutational distances between the derived and parent barcodes, we computed the Levenshtein (edit) distance between pairs of barcodes (as implemented in the Python C package `Levenshtein` (63)). Barcode read counts were merged if the edit distance was 4 or less, and if the minority barcode only mapped to one majority barcode at the minimum edit distance.

We then used previously acquired TnSeq data that maps the barcode identity to its transposon insertion location in order to identify which gene (if any) the barcoded transposon disrupted. Transposons that hit the first or last 5% of the gene sequence were excluded, as it is possible that these insertions do not result in disruption of production of the gene product. To ensure that barcodes at least begin their trajectories at a sufficiently high read count, if there were barcodes within a gene with low initial counts, *r*_0,*i*_ *<* 80, we summed the lowest (initial) count barcode into the next-lowest count barcode until min_*i*_ *r*_0,*i*_ *≥*80. We restricted our analysis to genes that had ≥ 4 barcodes, allowing us to gain confidence that the measured knockout fitness is not dependent on rare fluctuations or secondary mutations. Additionally, some barcodes went extinct during the course of the experiment, either due to genetic drift or selection; if a barcode went extinct, i.e. has 0 counts from *t*_*ext*_ to *T*, we would trim all time points after, but not including, *t*_*ext*_. We eliminated barcodes that go extinct after just one time point.

**Table S1.**
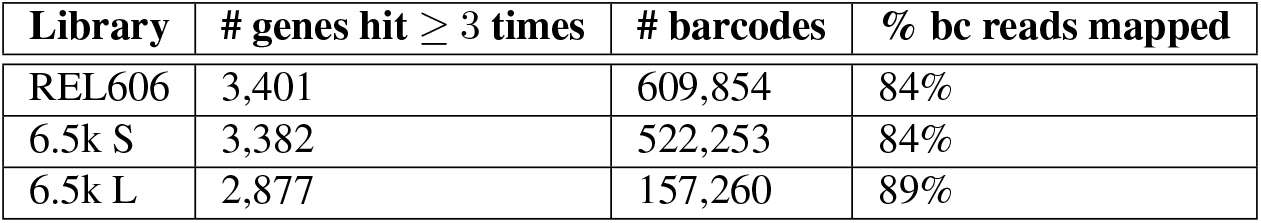
Summary of statistics of constructed RB-TnSeq libraries.

| Library | # genes hit $\geq 3$ times | # barcodes | % bc reads mapped |
| --- | --- | --- | --- |
| REL606 | 3,401 | 609,854 | 84% |
| 6.5k S | 3,382 | 522,253 | 84% |
| 6.5k L | 2,877 | 157,260 | 89% |

### S3.2 Probabilistic model of read count trajectories and fitness inference

To infer the fitness of individual genotypes from BarSeq count data, we must first understand what frequency trajectories we would expect for a given fitness, and how technical noise (e.g. from sample preparation and sequencing) and genetic drift affect those trajectories. Consistent with previous work (30, 31, 35), we construct a maximum-likelihood estimator to infer fitness from trajectories of barcode read counts, using a deterministic approximation of frequency dynamics.

On average, when the frequency of a lineage is sufficiently small *f*_*t,i*_ « 1, the frequency dynamics will exponentially grow/decay according to the genotype fitness, *s*, as well as the mean fitness of the population, 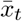 (see section S3.4),

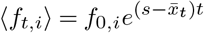

We measured the time in *generations*, which we measured for each time point in each experiment (see section S2.2). The reason we used a timescale of *1/generation* instead of e.g. *1/cycle* was to be able to better compare the magnitude of effects across experiments–e.g. the two exponential phase experiments had varying numbers of generations from cycleto-cycle and between strains (due to differences in exponential growth rates). However, the fitness effects can be scaled by a factor of approximately 6.64 to get per-cycle fitness effects, at least in the 1:100 serial dilution experiments. The two sources of noise–genetic drift and measurement noise– both arise from counting processes, so the combined noise will follow var(*f*_*t,i*_) ∝ ⟨ *f*_*t,i*_ *⟩* (see section S3.3). To account for the inherent discreteness of counting sequencing reads– especially important to accurately model deleterious genotypes that quickly drop to low frequencies–we modeled the observed counts at time *t* (always measured in generations) of barcode *i* inserted in a given gene, *r*_*t,i*_, as a negative binomial random variable,

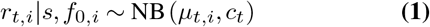

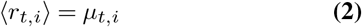

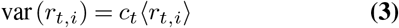

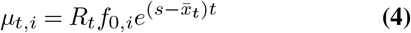

Where *R*_*t*_ is the total number of counts, and *c*_*t*_ is the measured variance parameter. The final likelihood for the fitness, *s*, of a given gene knockout is obtained by numerically integrating over *f*_0,*i*_ (’integrated likelihood’ with a flat prior)– incorporating the uncertainty in the intercept nuisance parameters into the fitness estimate and turning the problem into a one-dimensional maximum likelihood–and then combining the likelihoods of all barcodes inserted into the gene,

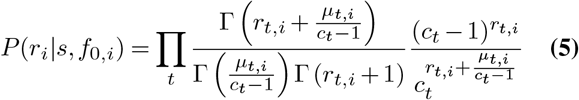

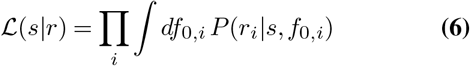

The point estimate of the knockout fitness, *ŝ*, is then numerically computed as the maximum likelihood, and the standard error is approximated as the inverse, square-root observed information,

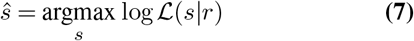

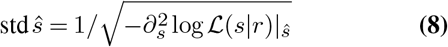

We ran biological replicates for all experiments reported here; to obtain combined genotype fitness estimates across replicates we simply multiplied the likelihoods together, repeating the maximum likelihood procedure.

As the majority of barcoded knockouts are neutral or nearly so (*s* ≈ 0), we must have a method to distinguish between likely neutral and selected knockout mutations; this can be accomplished by computing a p-value under the null hypothesis *s* = 0. For ease of computation and generality we compute the p-value as the posterior probability that the likelihood ratio between null and alternative hypotheses is greater than 1, i.e. the probability that the data more strongly support the null hypothesis over the alternative,

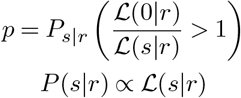

This convenient definition has been shown to be equivalent to the frequentist definition of the p-value using a likelihood ratio test statistic (if the distribution is invariant under transformation) (64, 65), and does not require asymptotic approximations.

In practice, this p-value can be calculated by first, finely discretizing the likelihood curve along *s* and normalizing it to get the posterior,

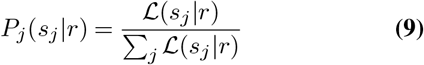

Then, calculating the log-likelihood ratio along all discretized *s* values,

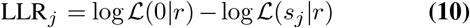

And finally, summing to get the posterior probability that the data supports the null hypothesis more than the alternative, where *I*[·] is the indicator function,

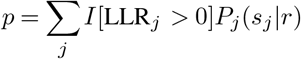

We used the standard method of Benjamini & Hochberg to control for the false discovery rate at *α* = 0.05.

### S3.3 Estimation of error parameters

In order to estimate fitness of individual genotypes from BarSeq data, we must first obtain an estimate of the error parameters for each time point in the experiments. There are two distinct sources of noise in our BarSeq measurements–measurement (technical) noise, arising from library preparation and sequencing error, which is uncorrelated in time, and variance due to genetic drift, which accumulates over time. Both sources of noise are count processes, where the variance of barcode population frequencies will be proportional to the mean,

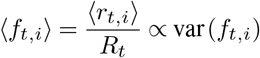

In order to eliminate the dependence of the variance on the mean, we apply a variance-stabilizing transformation,

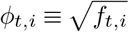

The variance of barcode frequencies of neutral lineages over two time points will then depend on the variance that has accumulated due to genetic drift, as well as the technical noise at the sampled time points. If there are sufficiently many read counts/individuals such that the central limit theorem applies, the variances will simply be additive,

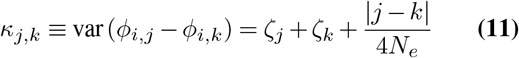

Where *ζ*_*t*_ is the technical noise at time point *t, N*_*e*_ is the effective population size, and | *j* − *k* | is the number of transfers performed between times *j* and *k*. The above equation defines a set of linear equations, with *ζ*_*t*_ and *N*_*e*_ as unknown parameters.

We can measure *κ*_*j,k*_ for all possible combinations of *t*_*j*_ and *t*_*k*_ given large enough set of neutral barcodes. Our RB-TnSeq libraries have a large number of transposons that were inserted into intergenic regions, the vast majority of which presumably have no fitness effect; thus, we use these intergenic barcodes as our set of putatively neutral barcodes. We confirmed that our measured *κ*_*j,k*_ did not systematically vary as a function of *r*_*j*_ (Figure S1), indicating that the expected mean-variance relationship, var(*f*_*t,i*_) ∝ ⟨ *f*_*t,i*_ *⟩*, is consistent with our data.

We only included intergenic barcodes that satisfy 50 *< r*_*t,i*_ *<* 500, as our computation depends on having sufficiently many counts such that the central limit theorem applies, and barcodes at a higher frequency are more likely to have acquired secondary mutations and be impacted by selection. In order to further guard against the effects of potential ‘outlier’ barcodes (those with non-neutral fitnesses), we compute variance estimates, 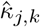, with a more robust measurement of variability, the median absolute deviation (MAD),

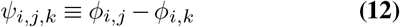

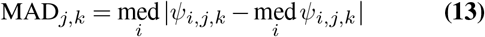

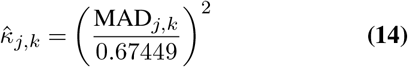

We resampled barcodes with replacement (standard bootstrapping) 500 times to compute the relative errors on the 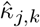 measurements. To decompose variability into the correlated (1*/N*_*e*_) and uncorrelated (*ζ*_*t*_) components, we numerically minimized squared error of the expected relationship (eq. 11) between the noise parameters and the measured 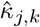, with inverse variance weighting,

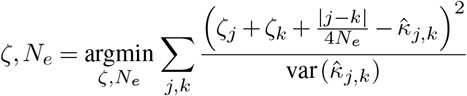

We subjected the minimization to the constraint that 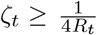, i.e. technical noise must be at least as large as variance due to sampling. After converting the variance parameters from frequencies back to read counts, the total marginal variance parameter at a single time point is,

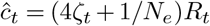

The number of intergenic barcodes included varies across RB-TnSeq libraries, experiments, and time points, but approximately on the order of ∼104 intergenic barcodes are used to estimate the variance parameters. The errors on the estimated *ĉ*_*t*_ are generally small (≲ 1%), so the point estimate *ĉ*_*t*_ was directly used for all downstream inferences.

### S3.4 Estimation of mean fitness dynamics

As beneficial mutations increase in frequency, and deleterious mutations decrease, the mean fitness of the population changes over time, impacting the rate of frequency change of all genotypes in the population. To estimate the mean fitness dynamics for each experiment, we can track the dynamics of neutral genotypes, again using the large set of intergenic barcodes. We obtain an estimate of the mean fitness between times 0 and *t* by simply taking the negative log slope over many barcodes,

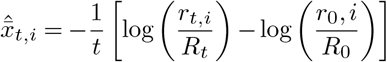

**Fig. S1.**
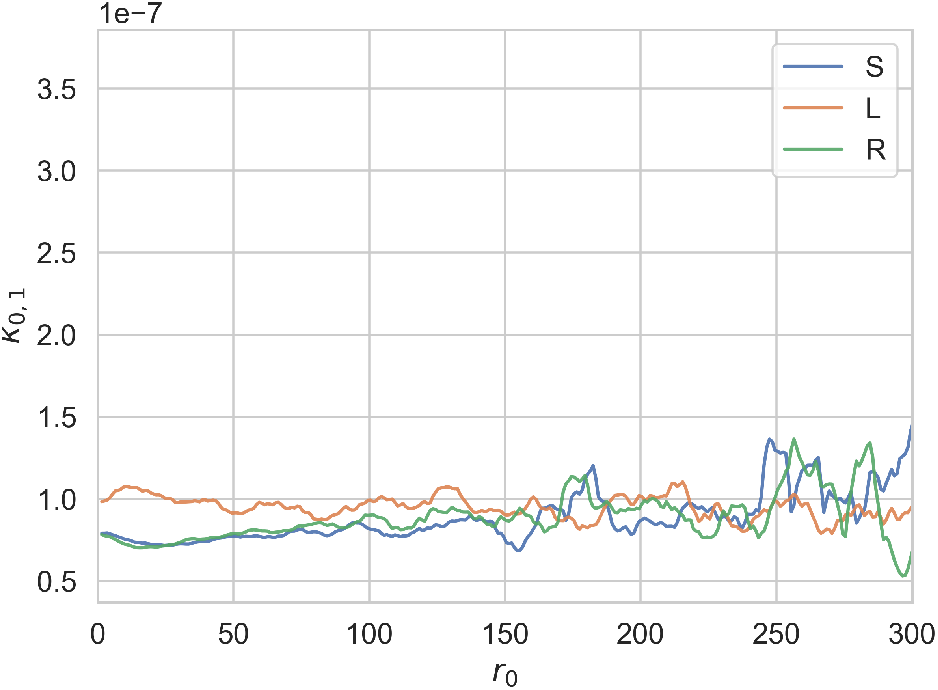
The measured noise parameter *κ*_*j,k*_ is consistently approximately constant as a function of initial number of barcode reads. Data is from S/L/REL606 monoculture experiments, replicate 1. Curves are smoothed with a moving average, ±2 reads.

As detailed in the previous section, it is advantageous to use robust forms of estimation to guard against the presence of outliers. Groups of ∼100 randomly selected intergenic barcodes with *r*_*t,i*_ *<* 500 were summed together to create “superbarcodes”, in order to improve individual estimates. The mean fitness 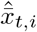 was estimated for each super-barcode separately, and then the final estimate 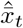 was obtained by taking the median over all super-barcodes. The standard error was estimated via the median absolute deviation between all super-barcodes, analogous to equations 13-14. Again, the point estimate 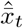 is used for all downstream analyses, as mean fitness error was consistently small.

### S3.5 Identification of putative outlier barcodes

We observed that some barcodes had trajectories that noticeably differed from the rest of the barcodes within the genotype, likely caused by the presence of secondary (selected) mutations that arose elsewhere in the genome or rare frequency fluctuations. We observed outlier barcodes with both beneficial and deleterious trajectories relative to the rest of the barcodes within the genotype. Problematically, some of these outlier barcodes were at high abundance relative to the other barcodes in the genotype, thus dominating the genotype fitness estimate. This necessitated a need to either accommodate outliers in our fitness estimation procedure or detect and reject outliers. We found that a number of robust estimators that we explored (e.g. maximum median/trimmed likelihood) had unreasonably high variance in fitness given our data (std *ŝ ŝ*). Thus, we opted to use a method to detect and reject outlier barcodes within genotypes. We based our outlier detection method on the resistant diagnostic *RD*_*i*_ introduced by Rousseeuw and Leroy (1987) (66), a high-breakdown measure of statistical deviation.

For every genotype with at *n*_*bc*_ ≥ 4 unique barcodes, we computed a fitness estimate for each barcode, *ŝ*_*i*_, via maximum likelihood (eqs. 5-8). We then used a resampling approach to randomly sample 200 different combinations of *n*_*r*_ = I*n*_*bc*_*/*21 barcodes, where samples are labeled *J*. To get an estimate of the ‘typical’ fitness, *ŝ*_*J,typ*_, of the barcodes within a gene, we either take the weighted median (*n*_*r*_ *<* 10) or weighted trimmed mean (*n*_*r*_ *≥* 10, trim 30% off each tail) of the resampled barcode fitnesses, where in both cases, samples are weighted by their inverse variance, *w*_*i*_ = 1*/*(var *ŝ*_*i*_). The weighted median is used for low number of samples, while the trimmed weighted mean is used for high number of samples, because the trimmed weighted mean generally has lower sampling variance when the number of samples remaining after trimming is sufficiently large. To compare the strength of evidence for a fitness of *ŝ*_*i*_ or *ŝ*_*J,typ*_ for barcode *i*, we compute the likelihood ratio,

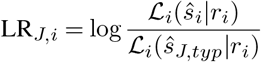

The deviation of barcode *i* from the rest of the barcodes in the genotype is then,

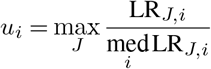

The final resistant diagnostic is finally calculated as a standardized version of *u*_*i*_,

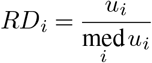

If *RD*_*i*_ > cutoff, then barcode *i* is considered an outlier and thrown away.

#### S3.5.1 Simulations

To determine an appropriate cutoff value, we performed simulations of the data generating process, and calculated the *RD* for each barcode within a simulated gene using the above method. Specifically, we simulated trajectories of lineage frequencies with *s ∈* {−0.02, 0, 0.02} gen^−1^ with the standard diffusion approximation, assuming *f* « 1,

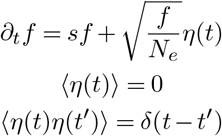

We ‘observed’ trajectories at the end of each ‘day’ (≈ 6.64 gen) for 4 days, and added measurement noise,

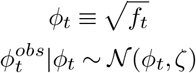

We used *N*_*e*_ = 108 *day* and *ζ* = 2 ∗ 10−8. We then grouped 20 simulated lineages together into a ‘gene’ (approximate median number of barcodes per gene in our libraries), with *n* ∈ {1, 2, 3} selected lineages (of the same sign), and the rest as neutral lineages. After calculating the *RD* for each simulated gene, we calculated the true positive/negative rate for calling a lineage as an outlier for a given threshold (Figure S2).

**Fig. S2.**
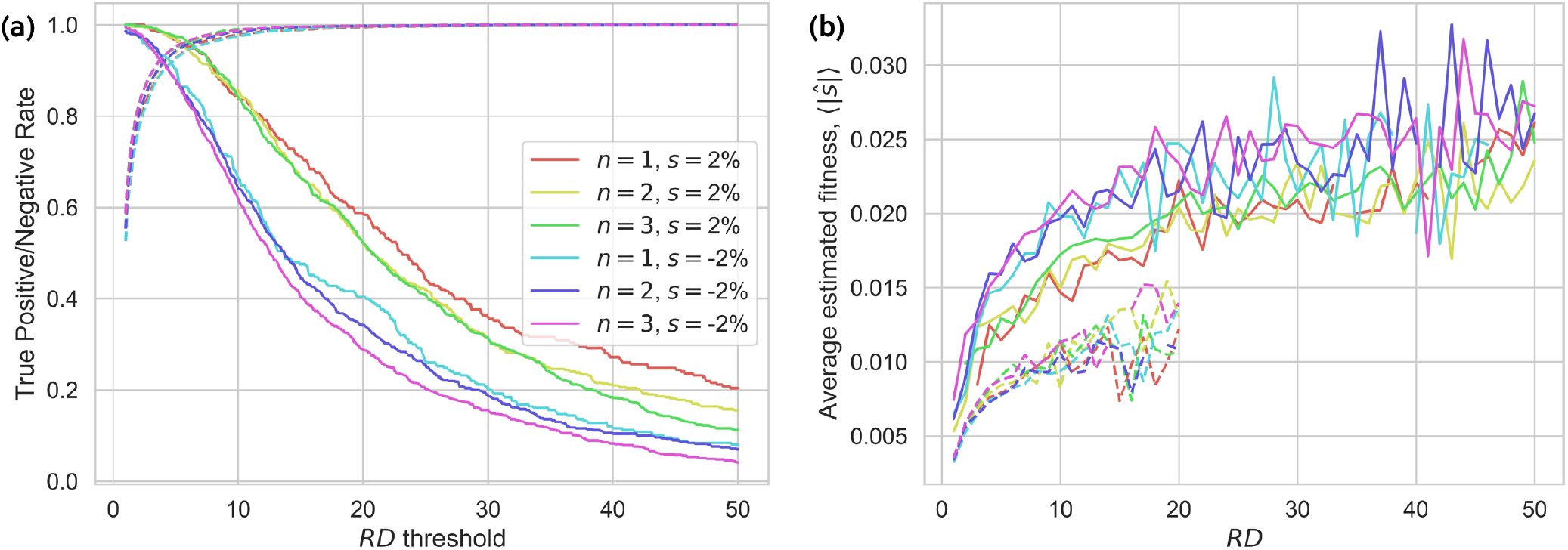
(a) Detection of selected, outlier barcodes in otherwise neutral genes. Dotted lines are the true negative rate, solid lines are the true positive rate. (b) Average inferred fitness (ie apparent fitness, differing from the true fitness by fluctuations) of barcodes with different *RD*s. Dotted lines are from neutral barcodes, solid lines are outlier barcodes. ‘Neutral’ barcodes with *RD* ≈ 6 have sufficiently large fluctuations to have trajectories that appear to have a 1% deviation from neutrality.

**Fig. S3.**
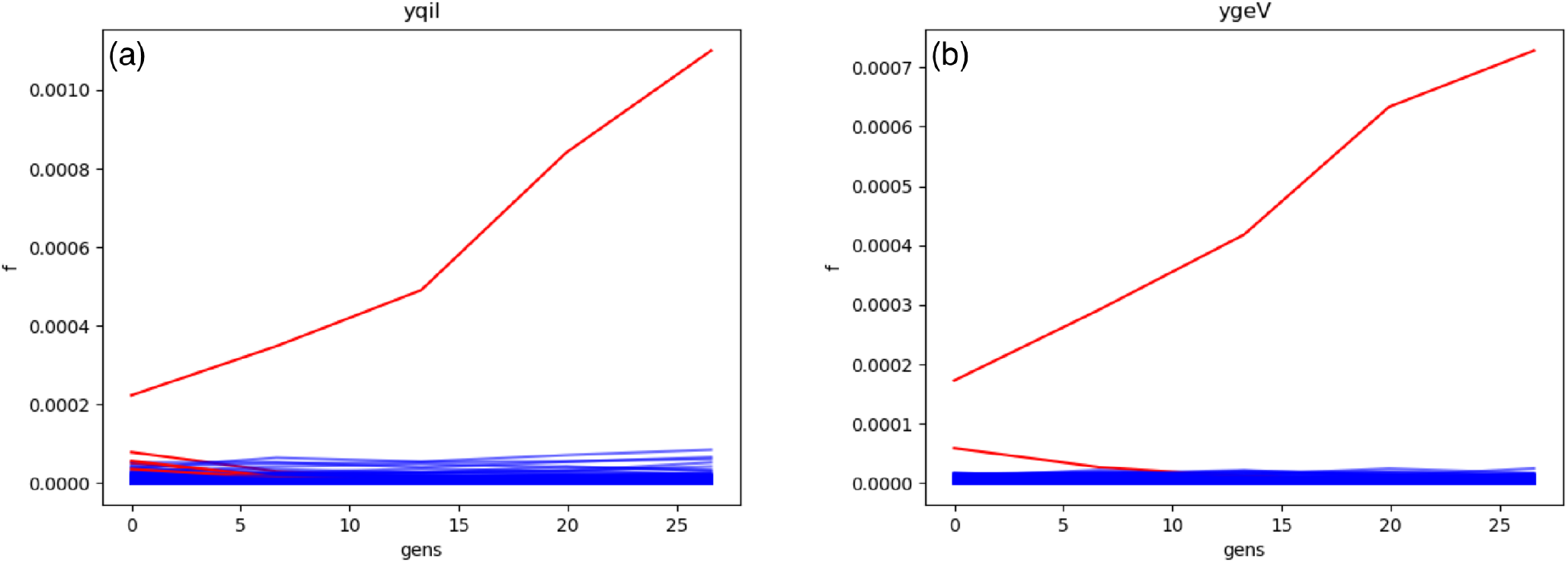
Examples of high-abundance outlier barcodes detected in otherwise neutral genotypes. Red barcodes were called as outliers. Examples taken from an experiment with the 6.5k S library in co-culture with L at the equilibrium frequency.

We can see that the method can sensitively detect relatively small, ∼2%, differences in fitness, while minimizing the number of neutral barcodes that are incorrectly thrown away. True positive rate decreases somewhat if there are multiple outlier barcodes within a gene, but the difference appears to be minimal, as expected from the construction of the *RD* as a high-breakdown deviance statistic. From the simulations, we chose a cutoff of 6, which only falsely throws out ∼5% of neutral lineages, while detecting ∼85 − 95% of outliers. This threshold also seems to empirically work with our data, detecting at least the most obvious outliers (see e.g. Figure S3).

### S3.6 Consequences of potential barcode frequency biases

One major assumption of the above analyses is that the frequency of barcodes from BarSeq data represents an *unbiased* estimate of the actual frequency of barcoded cells in the population. While we expect this assumption to generally hold, there are two major ways that this assumption could be violated: (1) if barcodes are differentially amplified due to e.g. differences in GC content, and (2) if genomic regions near the chromosomal origin of replication are present at a higher copy number due to fast growth. Both types of biases have been observed in some previous RB-TnSeq experiments (40, 41). We can check for the presence of frequency biases by comparing the inferred value of the error parameter *κ*_*t*_ (see section S3.3) for barcodes with different GC contents and across genomic positions, as biases in frequency measurements will change the apparent strength of genetic drift. We see that *κ*_*t*_ generally does not change across these conditions (Figure S4), and thus the aforementioned sources of frequency biases do not seem to be particularly prevalent or strong in our system.

Of course, other unknown sources of frequency bias could be present, or too weak to detect; but, under our inference pipeline, biases in frequency would only affect the variance of inferred *s*, not its expected value, as long as the bias across time points remains constant. We can see this by considering the deterministic (mean) dynamics of mutant frequencies *f* in a population with *m* genotypes,

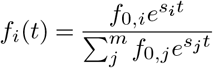

We could then include a strain-specific, constant multiplicative bias parameter, *γ*_*i*_. The observed frequencies would then follow,

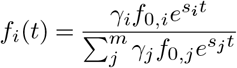

By observing these biased frequencies instead of the actual frequencies, we would infer *s*_*i*_ and *γ*_*i*_*f*_0,*i*_, therefore only biasing the nuisance intercept parameter.

As expected from the above analysis, there was no consistent, detectable correlation between genomic position and inferred fitness (Figure S6). However, there is one exception: in a couple of the L experiments, it looks like there is a dip in median fitness around ∼2.7 Mb, seemingly caused by a lack of neutral/beneficial variants. This position is about ∼1 Mb downstream from the origin of replication (3.8Mb), and ∼1 Mb upstream of the termination of replication and Dif site (∼1.5Mb). So it appears to be unlikely an artifact of uneven copy numbers or a DNA extraction bias. The origin of this signal is unclear, but seems to indicate that there is a region of the L genome that is more likely to have deleterious effects from knockout mutations. However, in any case, the dip seems to be isolated to a seemingly unremarkable portion of the genome, and thus does not call into question the general validity and assumptions of our model.

## S4 Analysis

### S4.1 Similarity of fitness effects across environments

To compute the correlation of knockout fitness effects across environments for a given genetic background (main text Figure 3), we first removed genes with noisy fitness effects (*σ*_*s*_ > 1%), then calculated the weighted pearson correlation coefficient, where genes are labeled *k* and environments are labeled *i, j*,

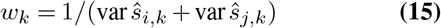

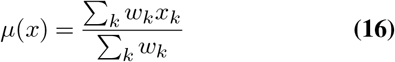

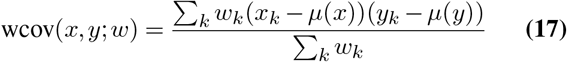

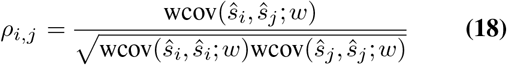

We then performed hierarchical clustering using Ward’s method across environments for each genetic background, with 1 − *ρ*_*i,j*_ as the distance metric. Environment pairs with *ρ*_*i,j*_ *<* 0 are set to 0 for the purposes of clustering, as there were few negative correlations, and all were small.

We used a bootstrapping procedure to estimate the statistical support for each cluster of environments. Using only the intersection of genes that passed across all environments, we performed standard resampling of genes with replacement, and then repeated the correlation measurement of knockout fitness values for each pair of environments. Then we repeated the hierarchical clustering and compared each branching of the original tree to the bootstrapped tree using the method of (67). We repeated the resampling procedure 5000 times for each genetic background and reported the average support for each clade.

We performed a principal components analysis on our data, using normalized fitness effects as the features. We only included genes that had measured fitness effects across all experiments. We normalized the fitness data separately for each experiment so that the scale of fitness effects was comparable across conditions. We first performed a quantile transform (to a gaussian distribution) on the fitness effects using `sklearn.preprocessing.quantile_transform`, and then subsequently centered and scaled the data to turn it into a standard normal. We performed the PCA with `sklearn.decomposition.PCA`.

### S4.2 Network of gene-by-gene correlations

To investigate potential relationships between genes in the different strain investigated in our work, we sought to quantify the degree of correlation of fitness measurements across all environments between every pair of genes, a quantity that has previously been referred to as cofitness (41). Highly correlated fitness measurements may indicate that genes are connected via gene regulation. In order to account for the fact that the measurement error in fitness measurements varies between genes and environments, we computed the cofitness of every pair of genes *i, j* as the weighted pearson correlation coefficient, where environments are labeled *k*, analogous to equations 15-18. We excluded genes that were not called as significantly non-neutral in at least one experiment, and genes with successful fitness measurements in < 4 experiments.

The vast majority of non-zero correlations are likely generated by chance, due to the relatively small number of environments where fitness is measured. Therefore, for each pair of genes, we generated a null cofitness distribution through a resampling procedure performed 300 times, by (1) randomly permuting the fitness assignments for both genes, (2) resampling each fitness value such that *ŝ*_*boot*_ ∼ 𝒩 (*ŝ*, std *ŝ*) (“parametric bootstrapping”), and (3) recalculating cofitness via equations 15-18. We then compared the measured cofitness to the null distribution to generate a 1-sided p-value. After correcting the set of p-values with a Benjamini-Hochberg FDR correction, we considered gene pairs to be signficantly correlated at *α* = 0.05, effectively drawing an edge between the two genes in the cofitness network.

After identifying statistically significant correlations between genes across environments, we sought to cluster genes into communities, without considering the magnitude or sign of the cofitness values. We used the ‘Fluid Communities’ algorithm (55), as implemented in the `networkx` python package (68), because of the flexibility of the algorithm, and the resulting communities had the highest modularity of all community-finding algorithms we explored. As the fluid communities algorithm is initialized stochastically, and requires pre-specifying *k* communities, we ran the algorithm on our data across varying community sizes, *k ∈* [4, 20], with 200 replicates for each *k* (Figure S17). We then picked the communities with the highest modularity for each genetic background. For the purposes of community finding, we treated all significant edges as the same, without considering the actual cofitness value of the edge. All community sets found had *modularity* > 0, indicating that genes were more tightly connected within their community compared to between communities.

**Fig. S4.**
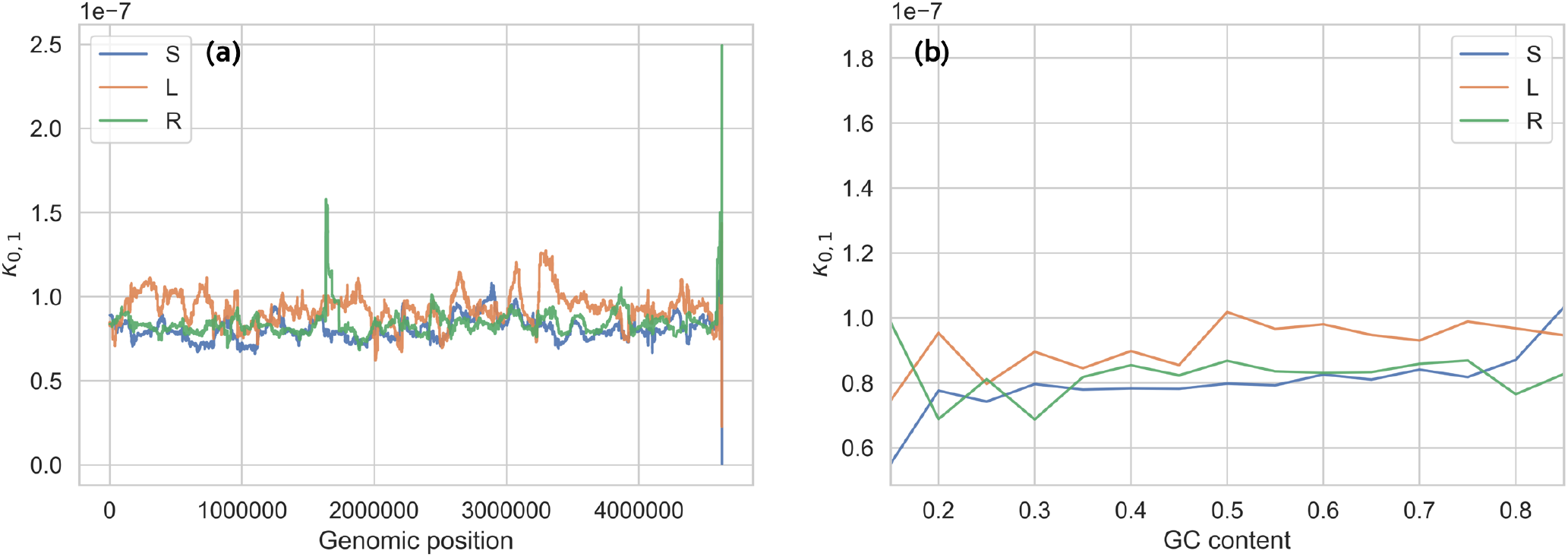
The measured noise parameter *κ*_*j,k*_ does not vary systematically over (a) genomic position, or (b) barcode GC content, indicating that these factors do not measurably bias barcode frequency measurements. Data is from S/L/REL606 monoculture (1) experiments, replicate 1.

Standard gene ontology enrichment analysis was performed on each community in each genetic background with the `goatools` python package (69), using Fisher’s exact test to find significantly over-represented annotations in a gene set, with an FDR correction and *α* = 0.05.

We sought to check if variance in fitness across environments for any given knockout could predict if two genes would stay in the same cluster across genetic backgrounds, as a control for the observed correlation with EcoliNet score. We average fitness variance across environments over the two knockouts of interest, referring to the quantity as ⟨ var(*s*) ⟩. We fit a logistic model with normalized EcoliNet score of the gene pair, *nscore* ≡ score/stdscore and *nvar* ≡ ⟨ var(*s*) ⟩/std ⟨var(*s*) ⟩ as the predictors (standard deviation is taken over all knockout pairs), and the probability that the two genes are together in strain 2, if they were together in strain 1 as the response variable, log *p*_*i*_/(1 − *p*_*i*_) = *nscoreβ*_*score*_ + *nvarβ*_*var*_ + *β*_0_ + ∈_*i*_. The results are shown in Figure S24.

It is known that community detection algorithms can have potential surfaces with large plateaus without a clear maximum, i.e. can give many solutions with similar modularity but different groupings (70). We wanted to see if the observed (mostly) “random reassortment” of genes among clusters between genetic backgrounds could be explained by this effect. Thus, we compared the optimal partition of each background to the 100 next-best partitions across all backgrounds (Figure S18). For each suboptimal partition, we asked if two genes were in the same cluster in the optimal partition, what is the probability that they are also in the same cluster in the suboptimal partition. We see that if we compare partitions in the same genetic background, this probability is around 40%, while it is around 10% when comparing partitions across background. This suggests that different reasonable partitions of the cofitness networks are much more similar within genetic backgrounds than between backgrounds. We also re-ordered the genes of the cofitness network such that they followed the ordering of another genetic background’s optimal partition (Figure 4B). It is apparent that replotting the cofitness matrix using another genetic background’s clustering does not produce noticeable structure. Together, these results suggest that while different reasonable partitions can give slightly different clusters, the observed reassortment of knockout fitness correlations among backgrounds cannot just be explained by failures of the community detection algorithm. We also investigated the extent to which the structure of our cofitness networks was driven by measurement noise (Figure S19, S20). We leveraged the fact that we had at least two biological replicates per experiment, and computed new cofitness networks (in the same manner as described above), only using either biological replicate “1” or “2”. We can see that even when the data is independently split, the cofitness networks within a genetic background are more similar than between backgrounds.

### S4.3 Genome evolution

We sought to understand if knockout fitness measurements could predict the probability that a gene would mutate in the LTEE. To that end, we downloaded clonal sequencing data from Tenaillon et al. (2016) (59), where the authors isolated and sequenced clones from a number of time points across all 12 lines of the LTEE, and identified mutations relative to the REL606 ancestor. We excluded synonymous SNPs from our analysis. A representation of the raw data can be found in Figure S28.

We then sought to understand if knockout fitness effects can predict if a mutation will appear in a gene in the Tenaillon et al. dataset, as a proxy for establishment. For REL606, classified a gene as mutated if a mutation appeared in one of the 12 LTEE lines (excluding mutator populations). For S and L, we classified genes as mutated only if they were present in the appropriate sublineage, i.e. in REL11830, REL11036 or REL11831, REL11035 for S and L respectively. We also excluded mutations that were already present in our S and L clones, which we determined from clonal sequencing data from Plucain et al. (2014) (46). We then fit a logistic model with knockout fitness effect as the predictor variable and gene mutated status (between time points) as the response variable,

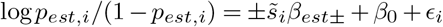

We fit two different coefficients for beneficial and deleterious mutations in each environment, *β*_*est*+_ and *β*_*est*−_ respectively. We only include genes that are putatively neutral, i.e. | *s* | *<* 0.005 and not called as significantly non-neutral, along with genes that are either significantly beneficial or deleterious, all at significance level *α* = 0.05. We normalized the fitness values by the median value of the non-neutral genes, i.e.

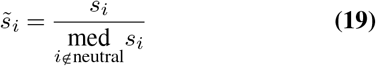

We use the logistic model implementation in the `statsmodels` python package (71). We used the standard method of Benjamini & Hochberg to control for the false discovery rate, pooling all tests across beneficial and deleterious coefficients. To test if there is a significant difference between REL606 logit slopes at 0-5k and 5-20k, we employed a permutation test. To construct a null distribution of the difference in slopes, for each gene we shuffled whether it ‘established’ (0 or 1) between 0-5k and 5-20k and recomputed the regression coefficients 1000 times, recording the difference. We then compared the actual difference in coefficients to the null distribution to get p-values.

### S4.4 Changes in gene expression

We used a microarray gene expression dataset previously reported by Le Gac et al. (2012) (16) to compare to our knockout fitness measurements, downloaded from the NCBI Gene Expression Omnibus (72), importing data with GEOquery (73). We primarily used the GEO2R tool to process the raw microarray data along with the R package `limma` (74, 75). After applying a log_2_ transform to the data, we ensured that all collected samples had approximately the same intensity distributions by performing a quantile normalization. Then, pooling all replicates within a strain, we fit a linear model to our data to determine the relative log-fold change in expression between different strains, taking into account the measured mean-variance relationship. A representation of the raw data can be found in Figure S27. We also compared the distribution of log-fold fitness effects between neutral and nonneutral genes (Figure 5B). We computed p-values to compare the distributions with standard Mann-Whitney U tests. We then fit a linear model to investigate if there was a correlation between fitness measured in a given environment, *s*_*i*_, and log-change in gene expression between evolutionary time points Δ*E*_*i*_, such that

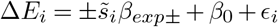

Similar to the gene establishment model, we fit two different coefficients for beneficial and deleterious mutations in each environment, *β*_*exp*+_ and *β*_*exp*_ − respectively (Figure S29). We only include genes that are putatively neutral, i.e. | *s* | *<* 0.005 and not called as significantly non-neutral, along with genes that are either significantly beneficial or deleterious, all at significance level *α* = 0.05. We normalized the fitness values by the median value of the non-neutral genes, in the same manner as equation 19. We fit the model with weighted least squares, as implemented in the `statsmodels` python package (71), with weights *w*_*i*_ ∝ 1/varΔ*E*_*i*_, to incorporate the fact that there are different levels of measurement error in the log-fold change expression for each gene. We used the standard method of Benjamini & Hochberg to control for the false discovery rate, pooling all tests across beneficial and deleterious coefficients.

As a control, we also investigated if our results would change if we excluded poorly expressed genes. It is perhaps the case that neutral knockouts are potentially a bad comparison class, because many of them may be poorly expressed at all times, and thus ineligible to undergo large changes in expression. We can test for this alternative hypothesis by focusing our analysis on solely initially highly expressed (50th percentile) genes, excluding poorly expressed genes. The results are shown in figure S30. The regression coefficients change somewhat, but not qualitatively, showing that the aforementioned hypothesis is not likely the driver of the signals we observed.

**Fig. S5.**
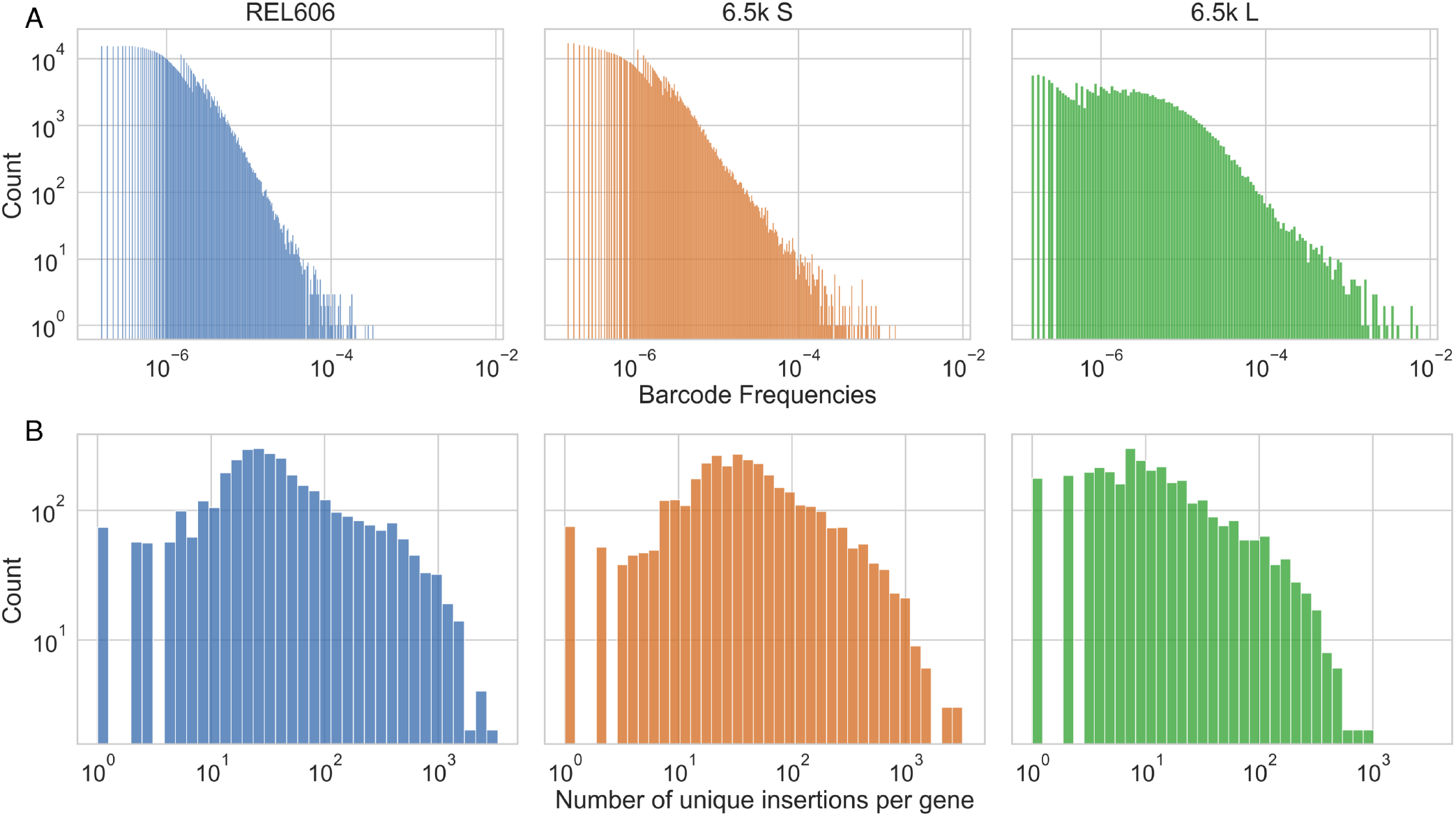
Statistics of RB-TnSeq libraries, (**A**) initial distribution of barcode frequencies in library populations, and (**B**) distribution of number of unique barcoded transposon insertions into each gene (cds).

**Fig. S6.**
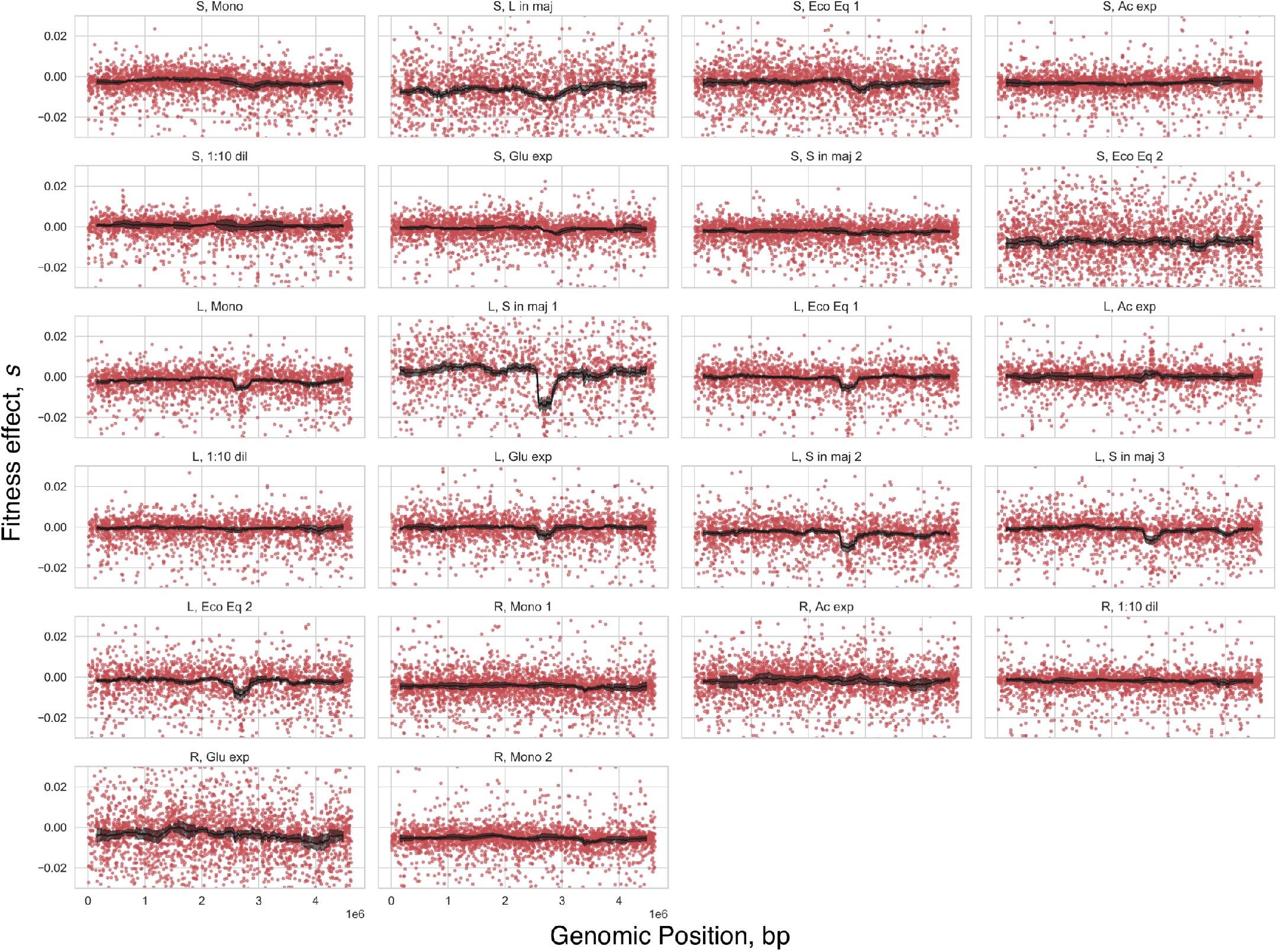
Relationship between genomic position and fitness effect. Red dots are the fitness effects of individual knockouts, black line is the rolling median fitness effect (error bars are standard errors).

**Fig. S7.**
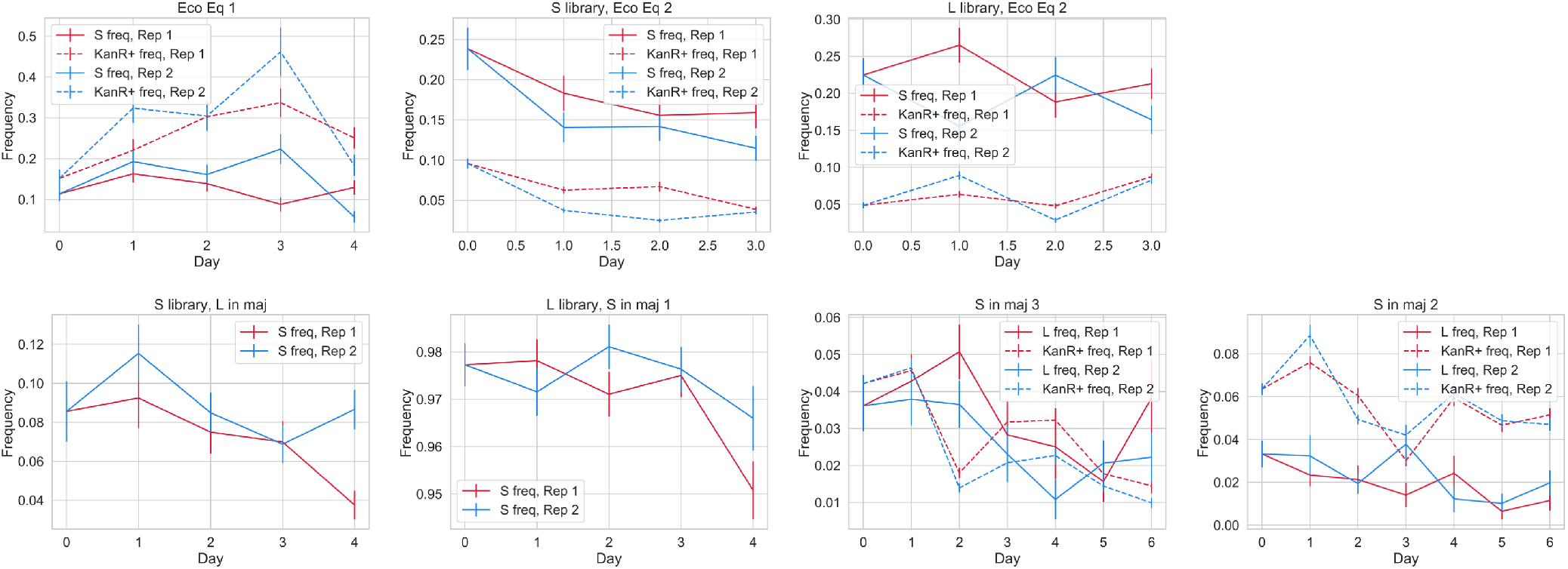
Frequency trajectories of mixed culture experiments from CFUs. For each coculture experiment, we diluted and plated cultures on both TM plates (S/L indicator plates) and LB/Kan plates (pulls out cells from the RB-TnSeq libraries). We didn’t plate experiments “S/L in maj (1)” on LB/Kan plates because we only cocultured wt S/L with L/S RB-TnSeq libraries respectively. Please note that each subplot is on a different scale.

**Fig. S8.**
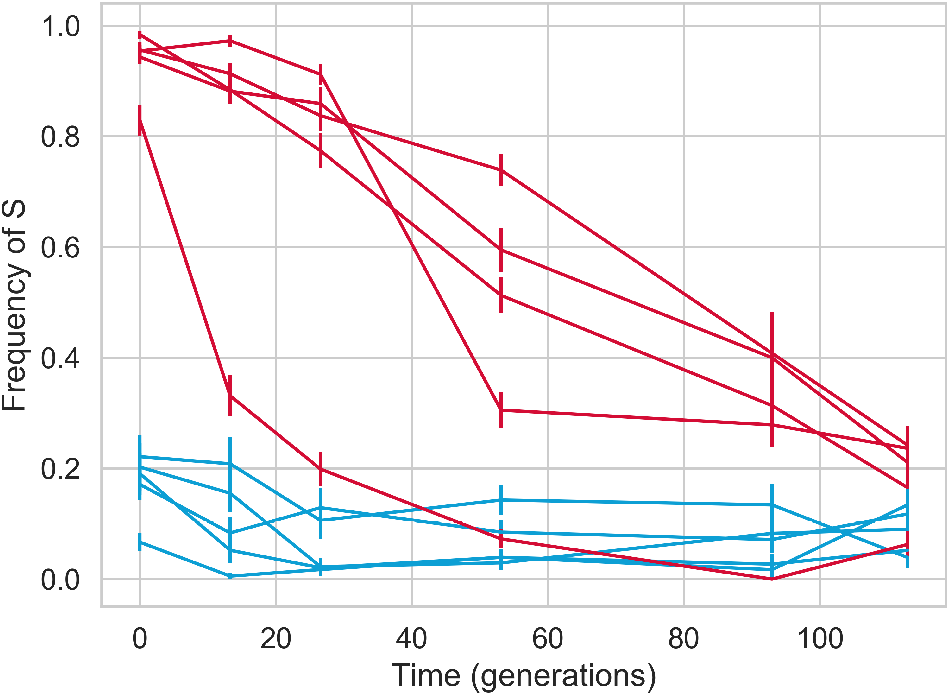
Measured S/L frequency dependent fitness and ecological equilibrium via CFUs on TM plates (S/L indicator plates). Independent cocultures of S and L wt clones were propagated in standard LTEE conditions. Red lines indicate cultures that were started at high frequencies of S, blue lines indicate cultures started at low frequences. Error bars represent standard errors.

**Fig. S9.**
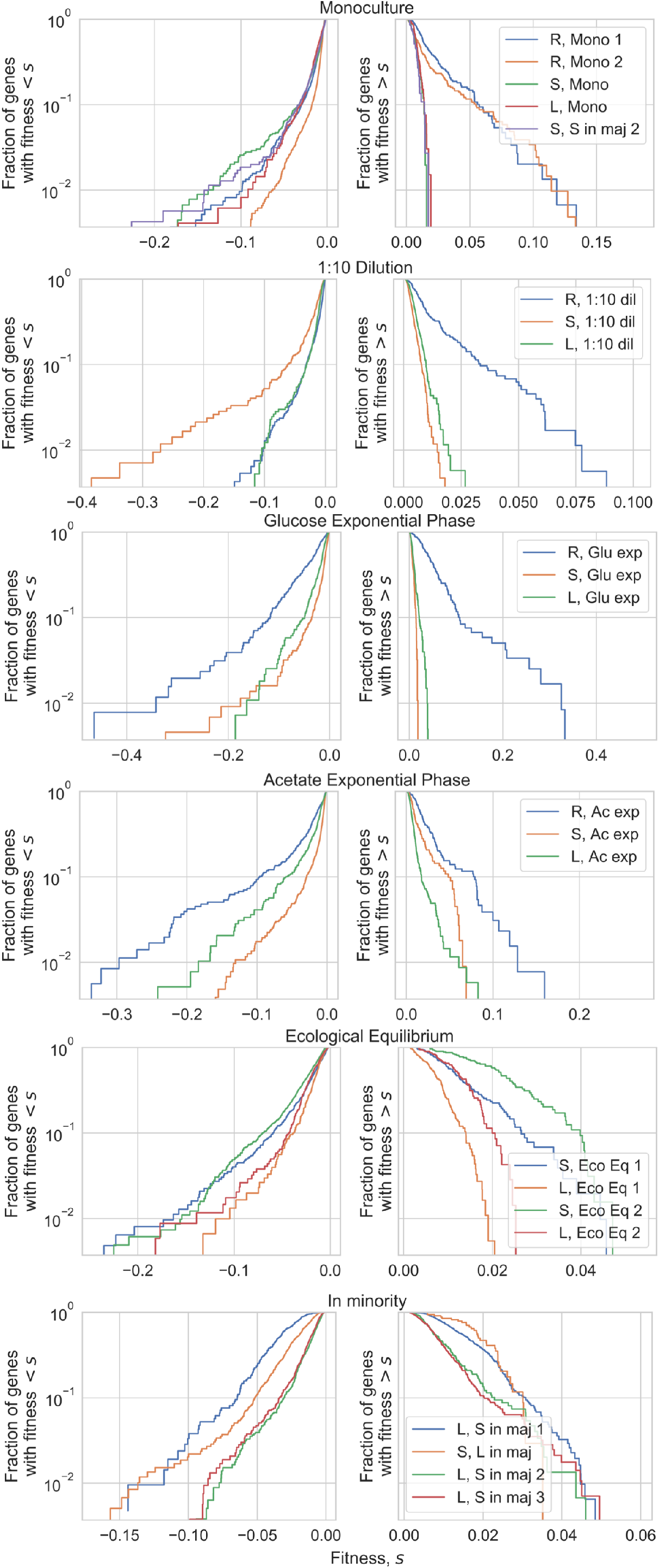
All measured DFEs across experiments, arranged by environment.

**Fig. S10.**
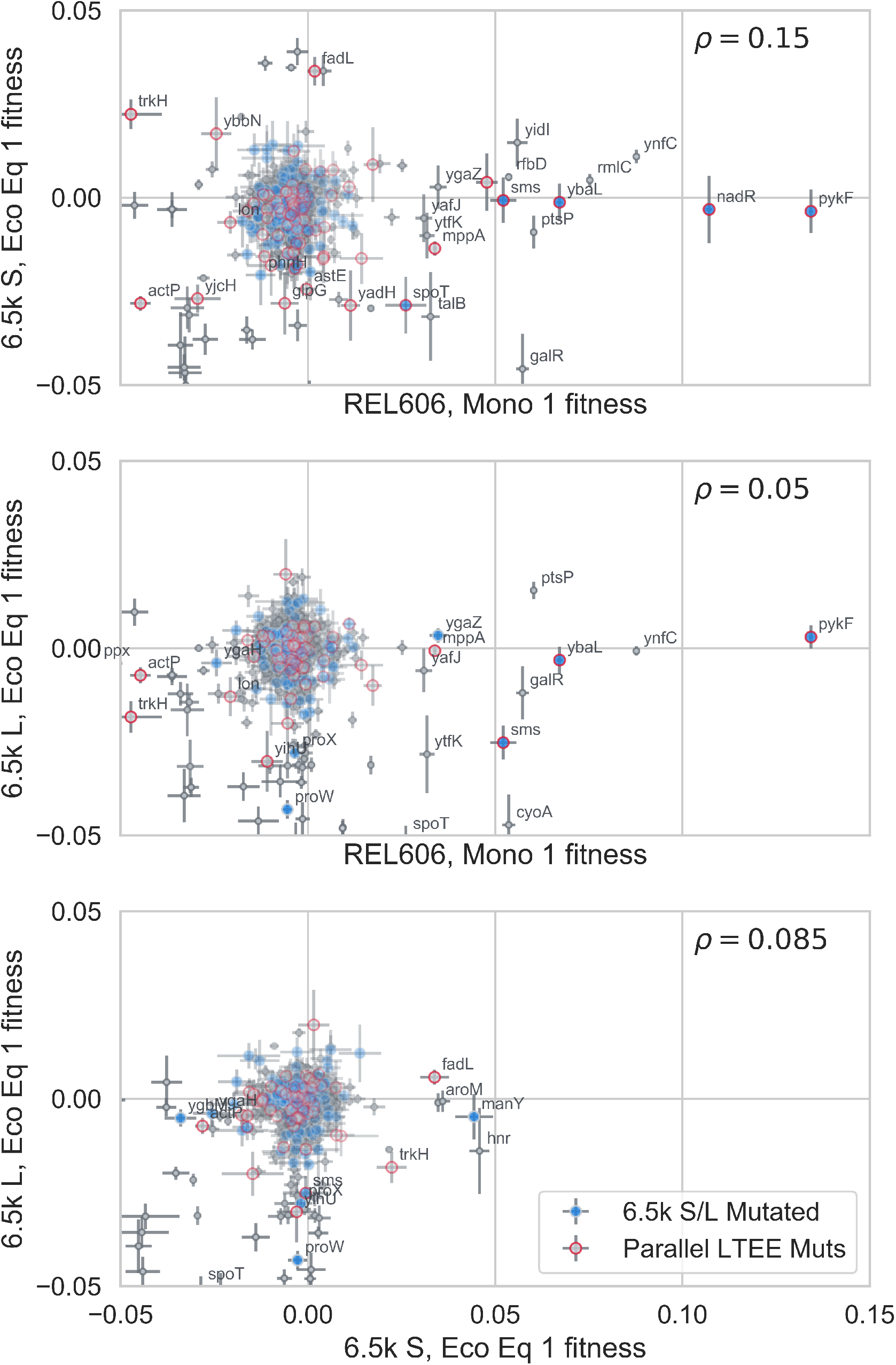
Comparison of fitness effects; identical to Figure 1F in the main text, except we highlighted all genes in mutated operons. It is still the case that there are many genes that did not get a mutation in their operon, but still changed from a beneficial to non-beneficial fitness effect across genetic backgrounds.

**Fig. S11.**
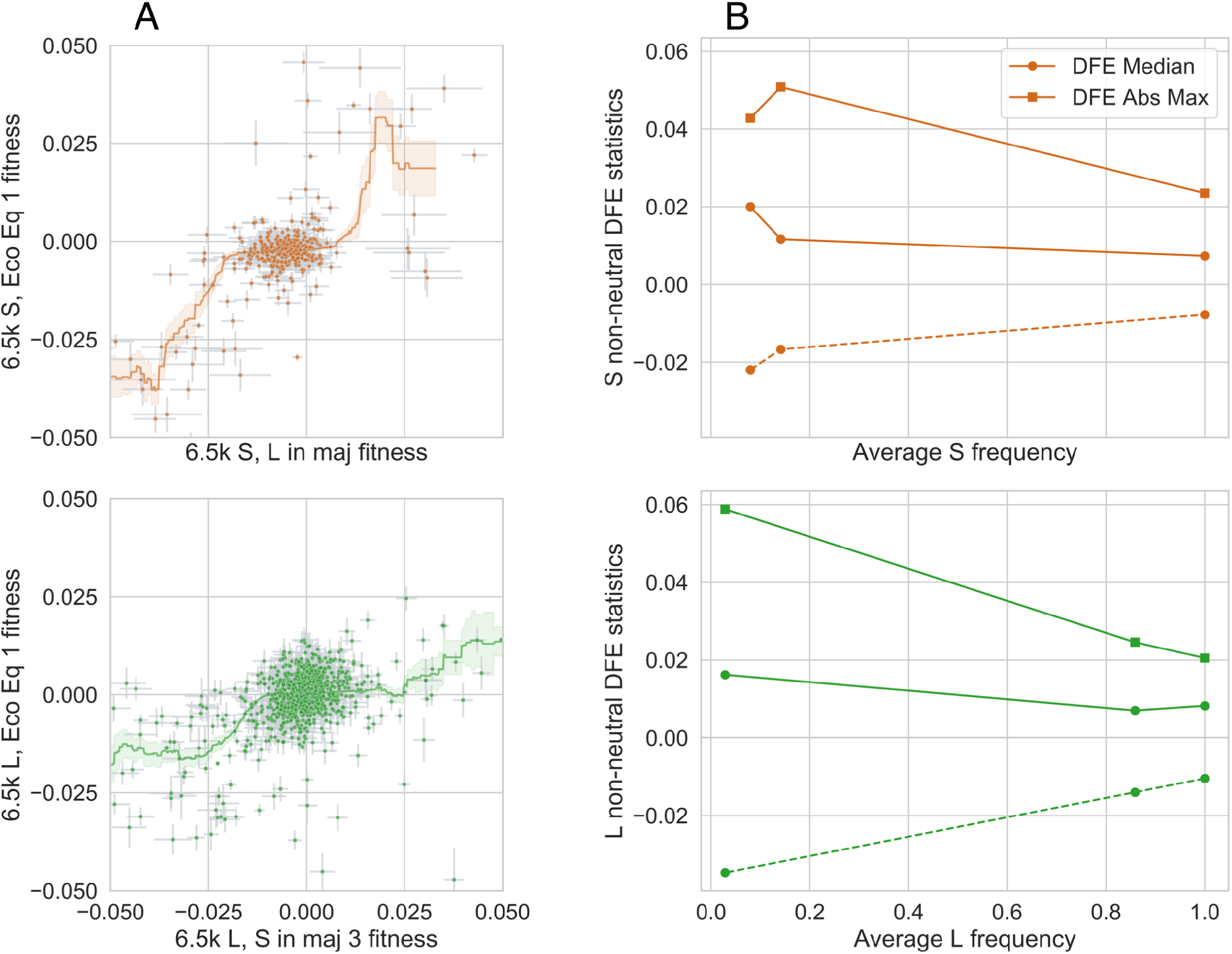
Frequency-dependent knockout fitness effects for both 6.5k S and L. (**A**) Similar to Fig 2A in main text, except comparing fitness at ecological equilibrium to fitness when the ecotype is in the minority. (**B**) Changes in summary statistics of the DFE as a function of ecotype frequency. Solid lines represent the beneficial side of the DFE, while dashed lines represent the deleterious side.

**Fig. S12.**
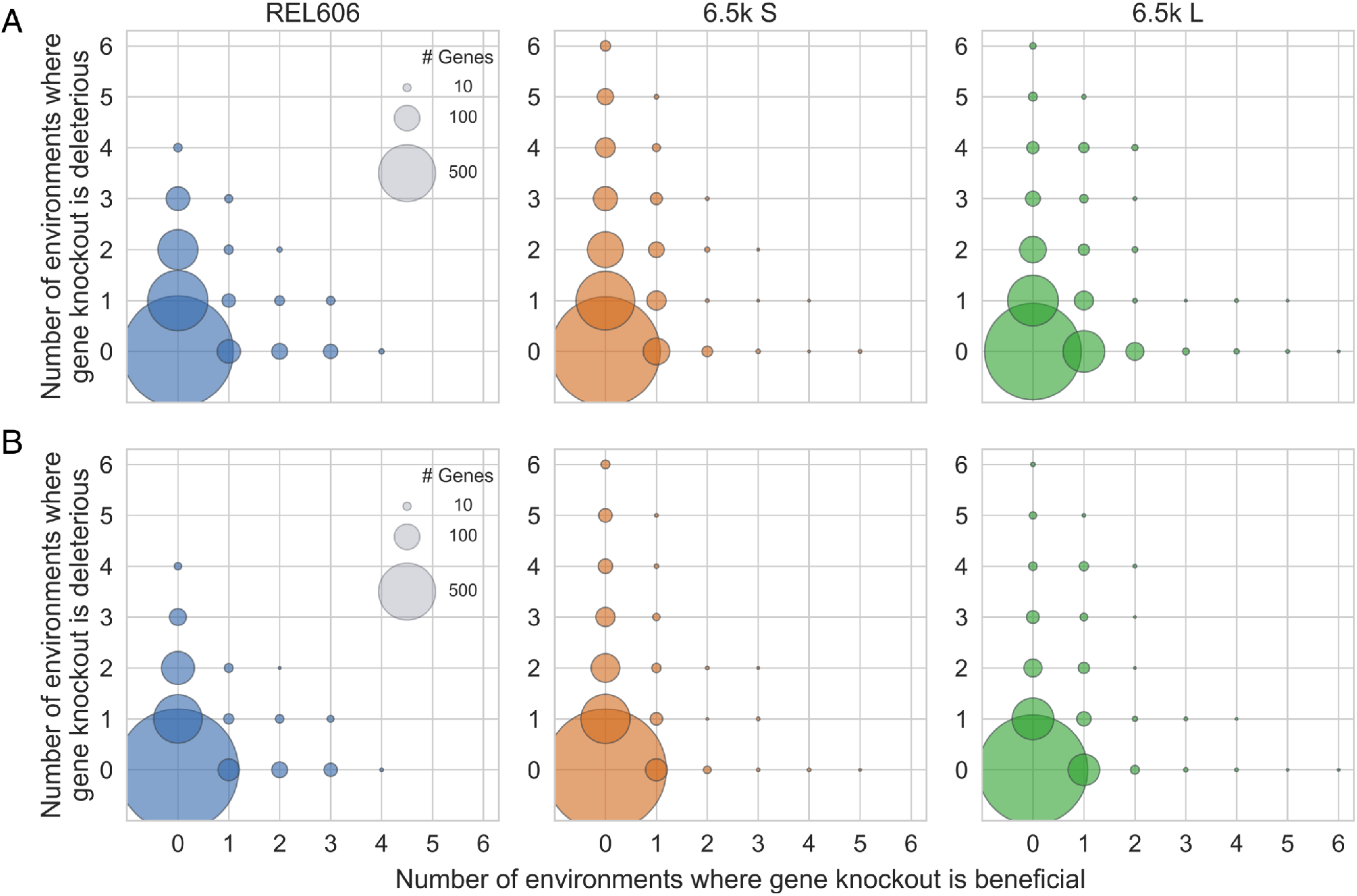
Fitness effect sign-flipping across environments. Same as Figure 2C (main text), but (post-FDR correction) p-value cutoff is reduced from 0.05 to (**A**) 10^−3^ or (**B**) 10−5

**Fig. S13.**
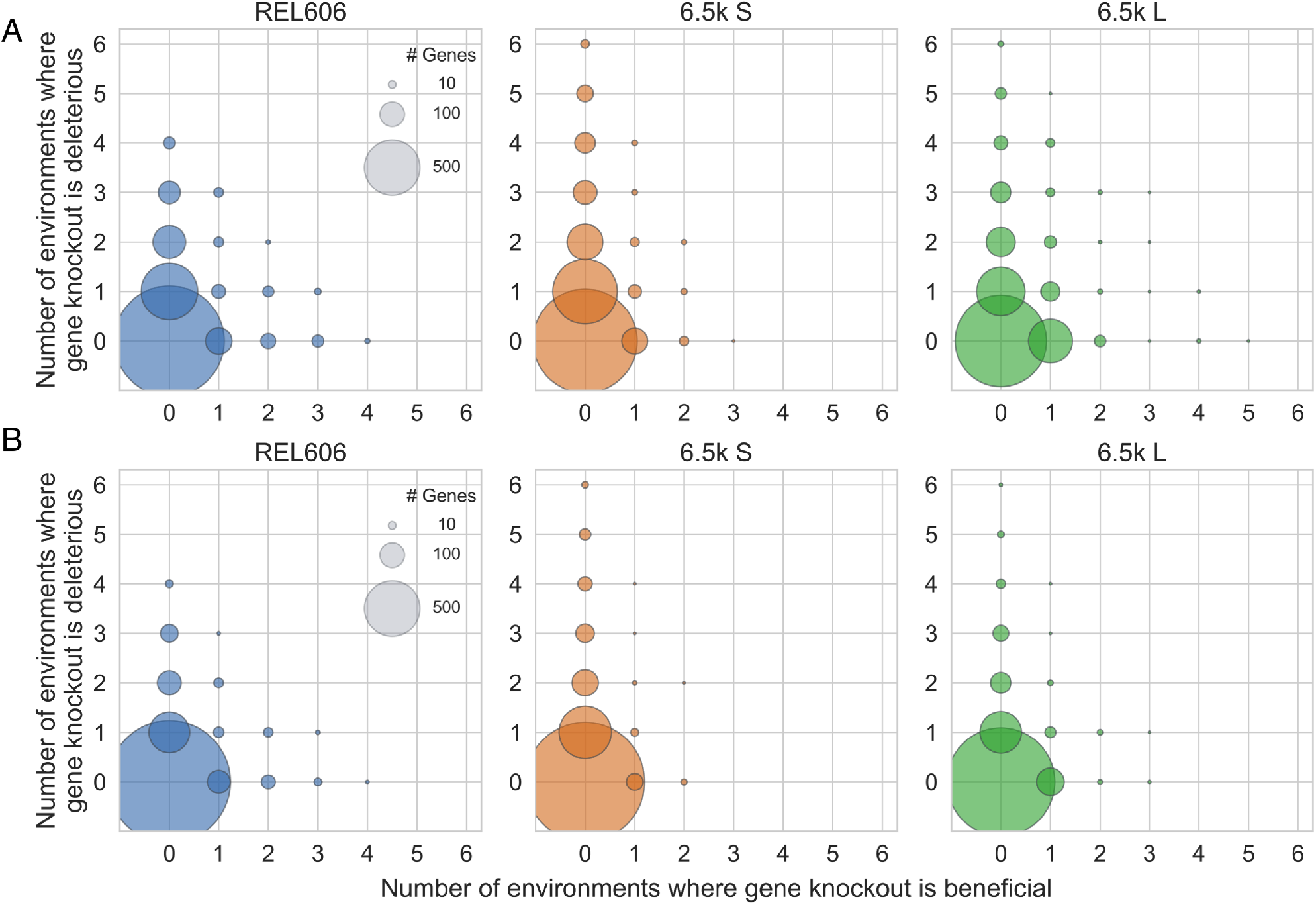
Fitness effect sign-flipping across environments. Same as Figure 2C (main text), but we only consider genes non-neutral with fitness (**A**) |*s*| *>* 1% or (**B**) |*s*| *>* 2%

**Fig. S14.**
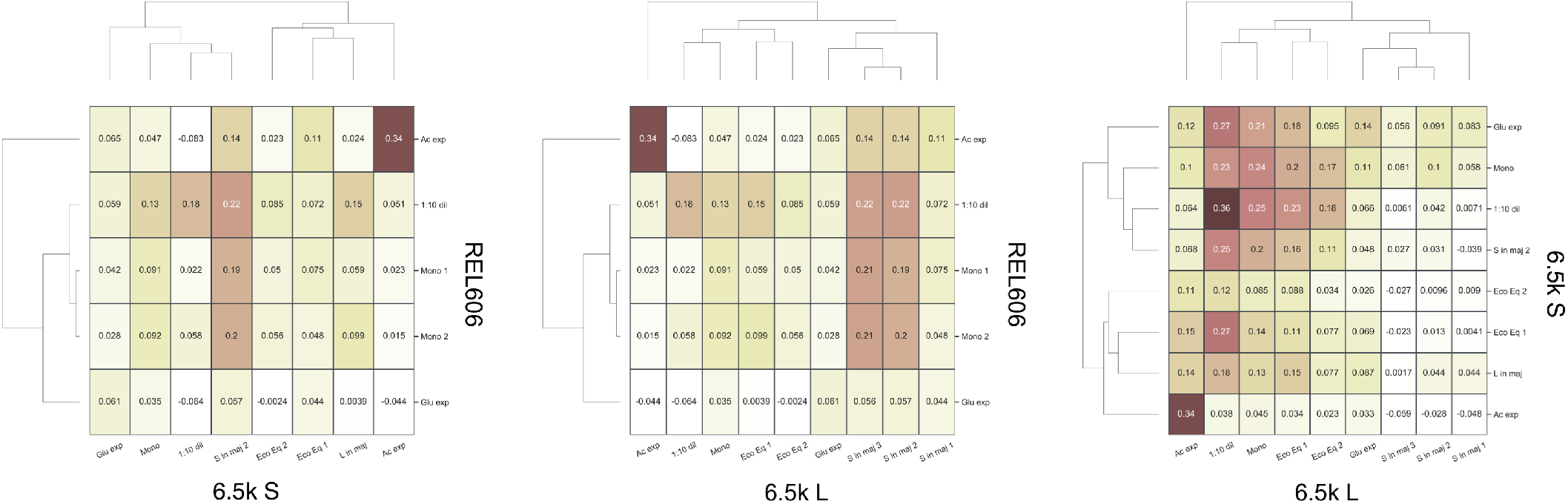
Fitness effect correlations between strains.

**Fig. S15.**
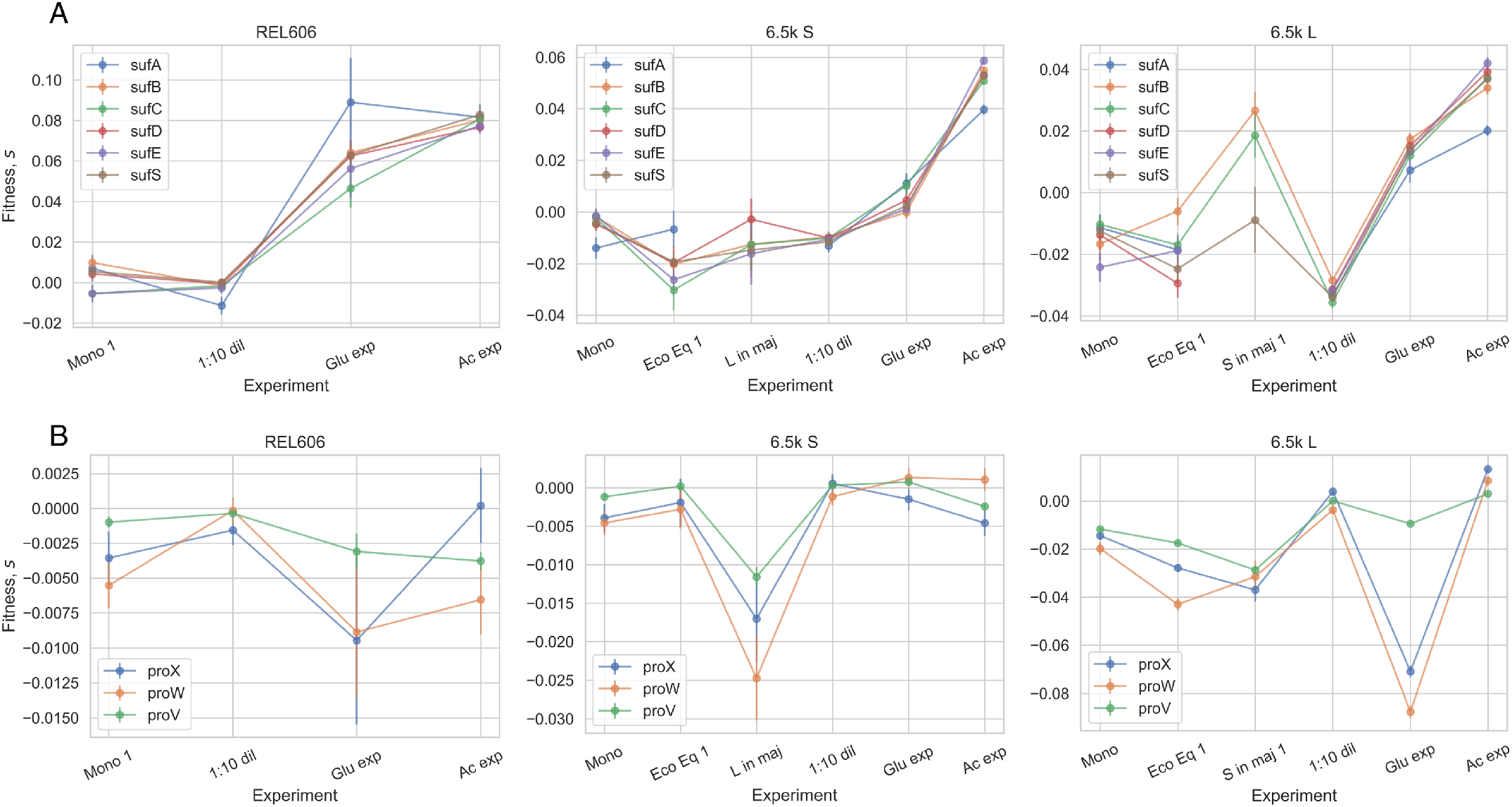
Fitness effects of (**A**) *sufABCDSE* and (**B**) *proVWX* operons in REL606 and 6.5k L/S.

**Fig. S16.**
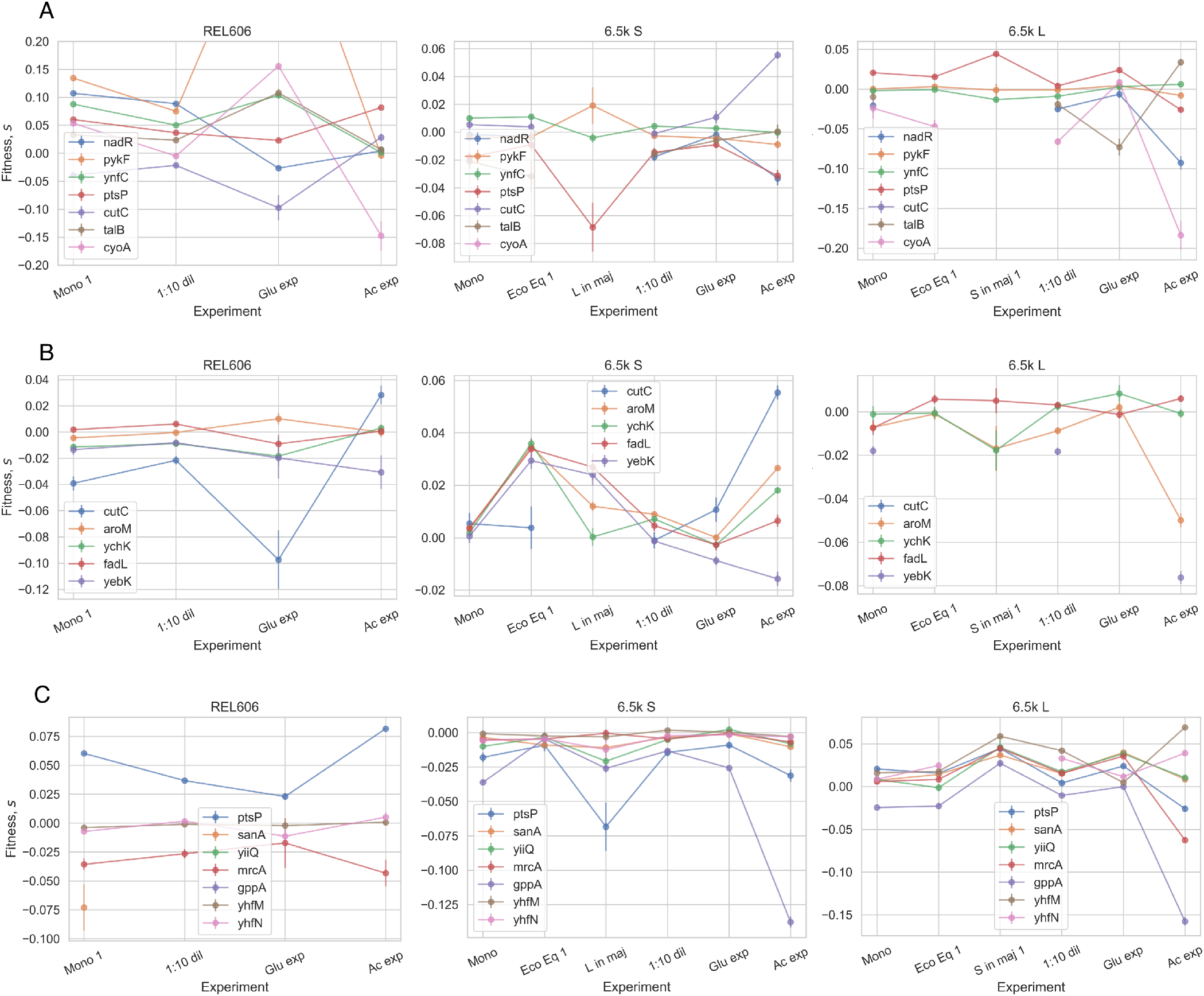
Fitness effects of knockouts across environments, where knockouts are beneficial in at least one condition on the (**A**) REL606, (**B**) 6.5k S, (**C**) 6.5k L background.

**Fig. S17.**
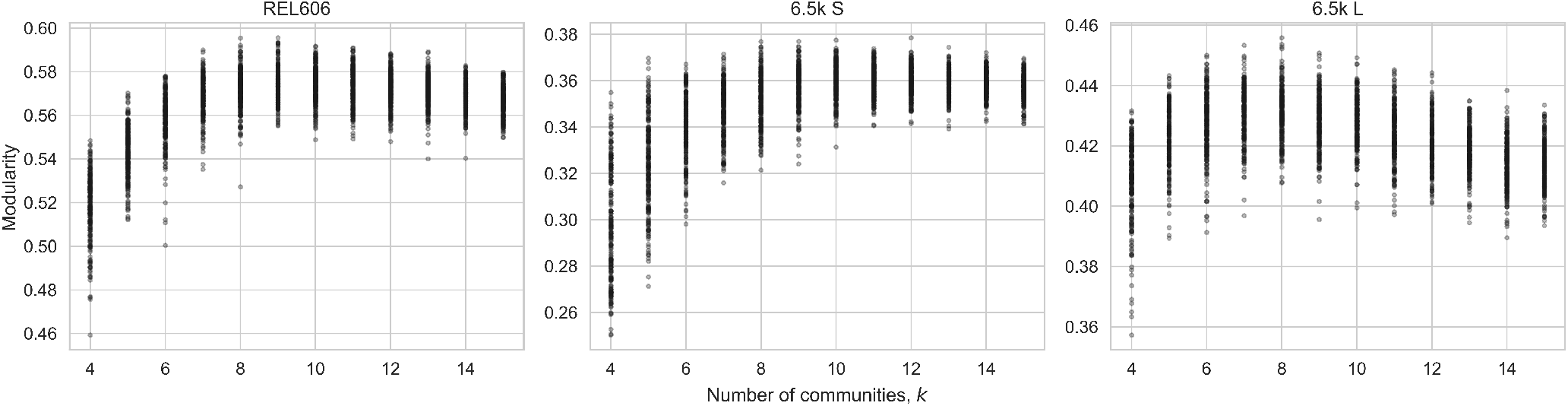
Modularity of cofitness clusters, across 200 (stochastic) initializations for different numbers of communities from 4 − 15.

**Fig. S18.**
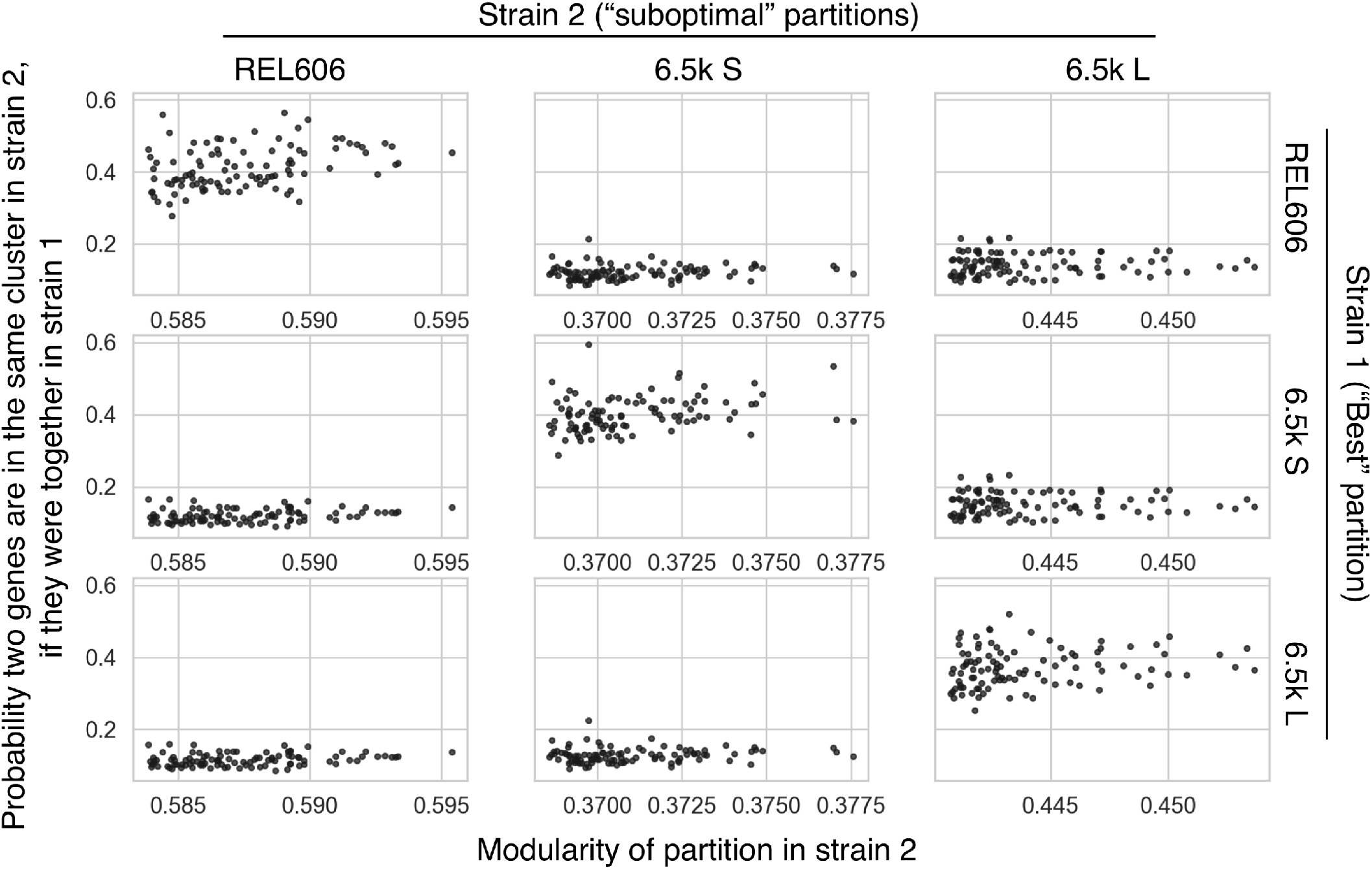
We compared the optimal partition of the REL606/S/L cofitness networks to the next 100 best (but suboptimal) partitions, also shown in Figure S17. For each suboptimal partition, we asked if two genes were in the same cluster in the optimal partition, what is the probability that they are also in the same cluster in the suboptimal partition. We can see that if we compare partitions in the same genetic background, this probability is around 40%, while it is around 10% when comparing partitions across background. This suggests that different reasonable partitions of the cofitness networks are much more similar within genetic backgrounds than between backgrounds.

**Fig. S19.**
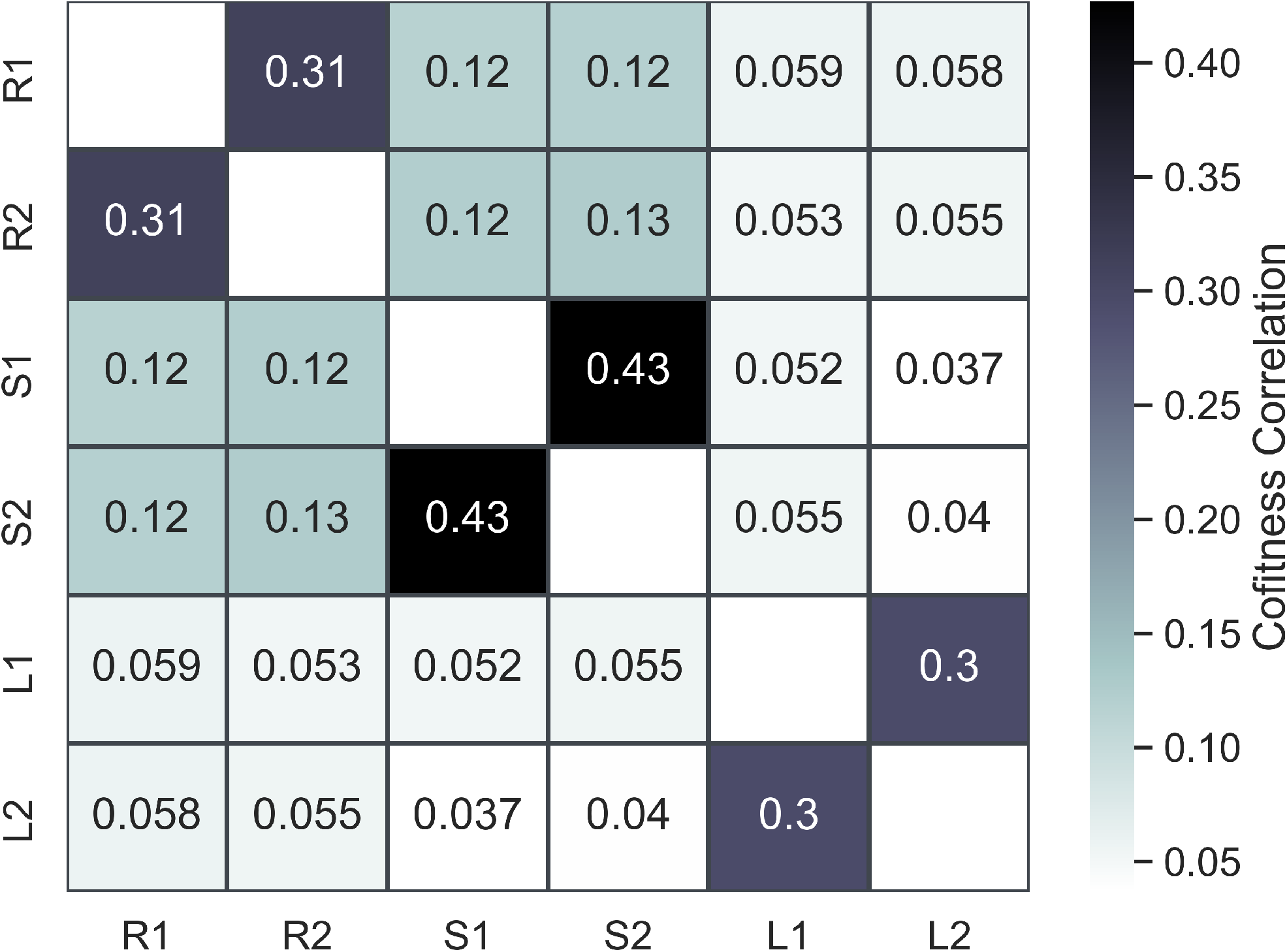
In order to better understand the extent to which the structure of our cofitness networks is driven by measurement noise, we re-computed the cofitness networks, only using one of the biological replicates per experiment for every experiment. We then computed the correlation of all cofitness values across all networks. We can see that even when the data is independently split, the cofitness networks within a genetic background are more similar than between backgrounds. In the figure, R, S and L refer to REL606, 6.5k S/L libraries, respectively, and 1 and 2 refer to using only biological replicates “1” and “2” from each experiment.

**Fig. S20.**
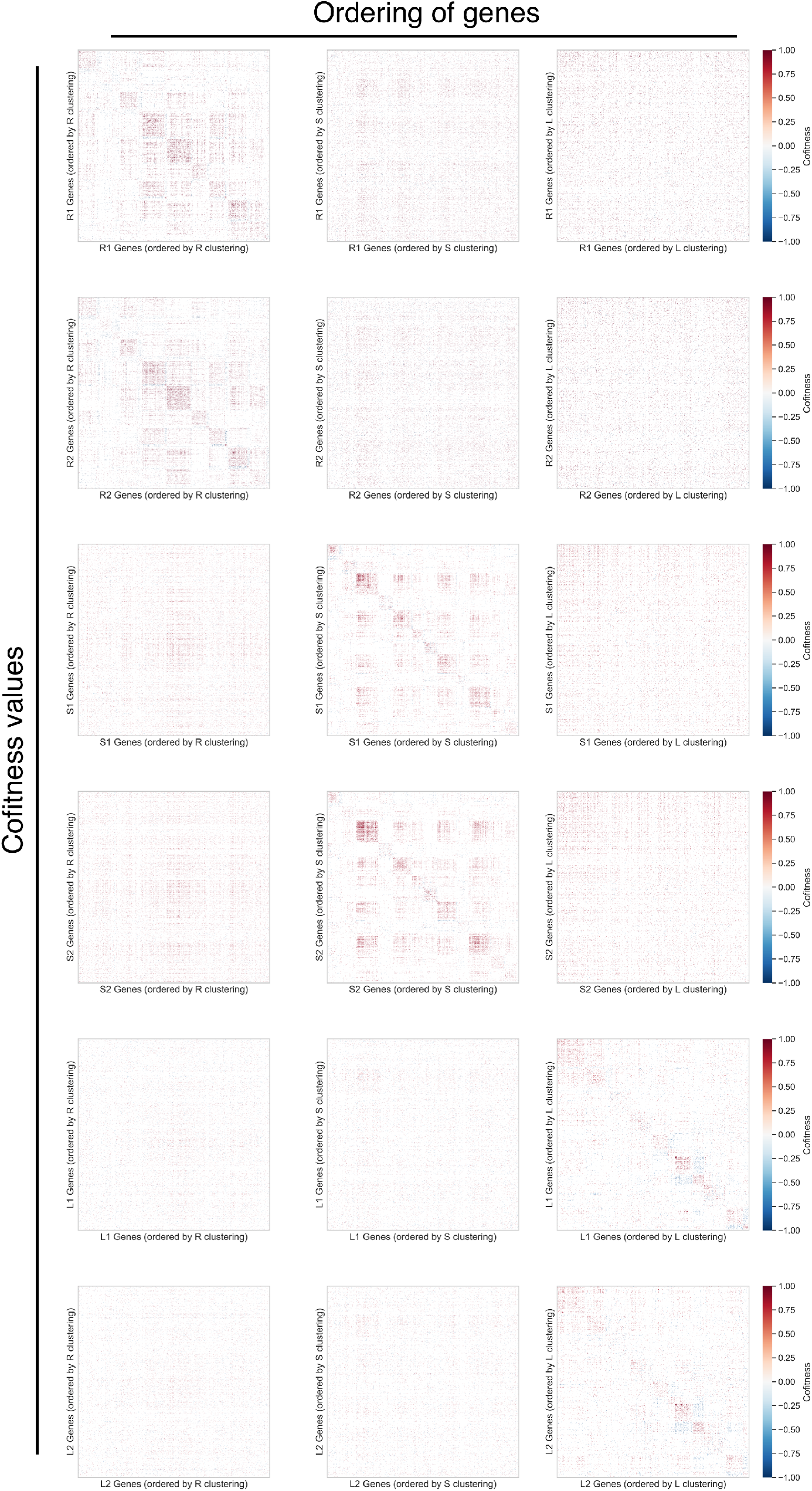
We repeated the cofitness clustering, as done in Figure 4B, using cofitness networks computed using only one of the biological replicates per experiment for every experiment (as done in Figure S19). We see similar results to Figure 4B, where clusters are visibly preserved only when clustered on the same background, albeit to a weaker extent. In the figure, R, S and L refer to REL606, 6.5k S/L libraries, respectively, and 1 and 2 refer to using only biological replicates “1” and “2” from each experiment.

**Fig. S21.**
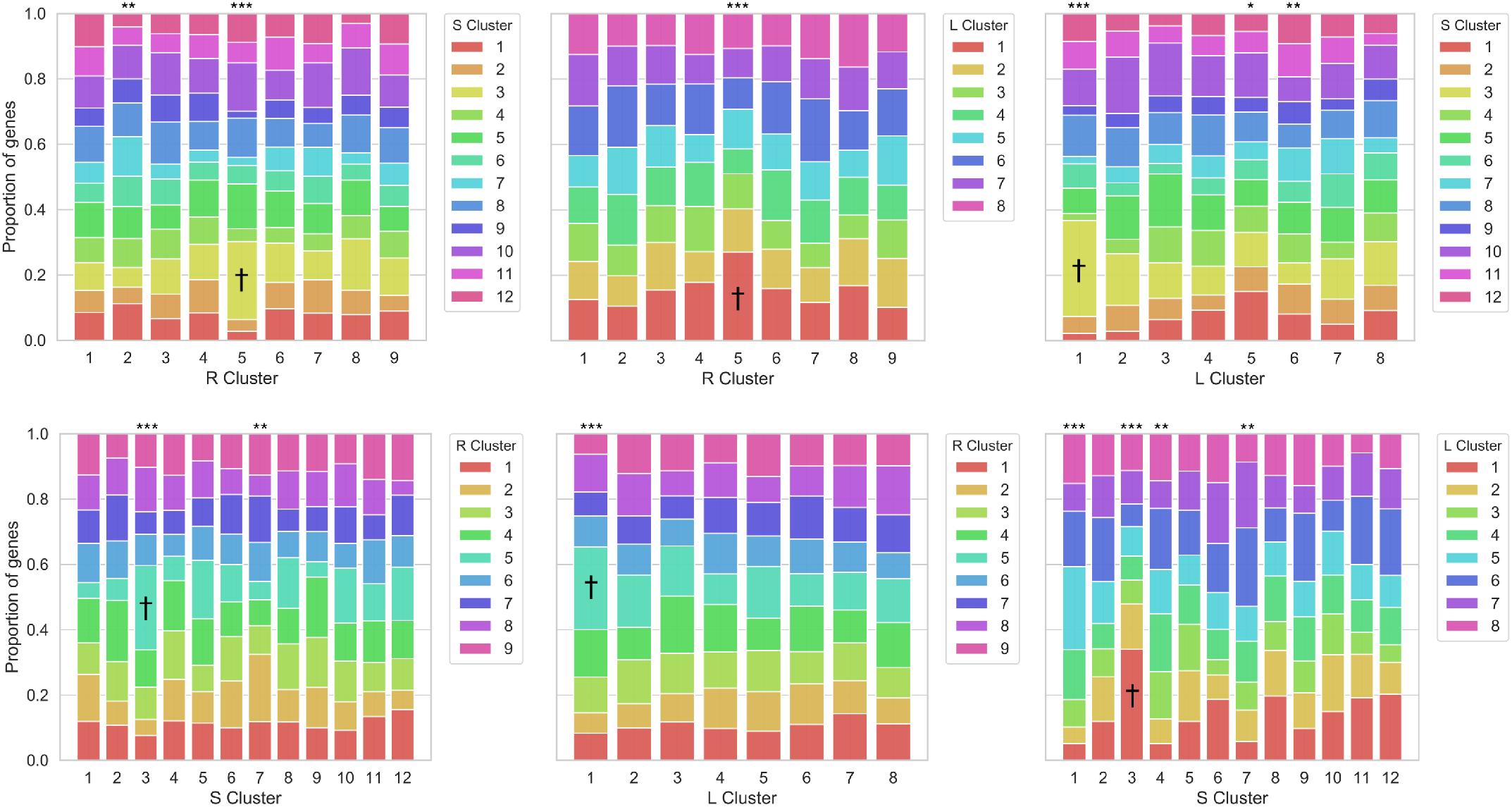
In order to explore how clusters of genes changed across genetic background, we calculated the fraction of genes in a given cluster that belong to a cluster in a different genetic background. We see that clusters are mostly not preserved between genetic backgrounds, with the exception of the clusters marked by asterisks, which show non-random sampling across genetic backgrounds (* *p <* 0.05, ** *p <* 0.005, *** *p <* 10^−4^; Pearson’s chi-squared test). ^†^ set of clusters across all three genetic backgrounds which share more genes than expected, driven primarily by adhesion-related genes.

**Fig. S22.**
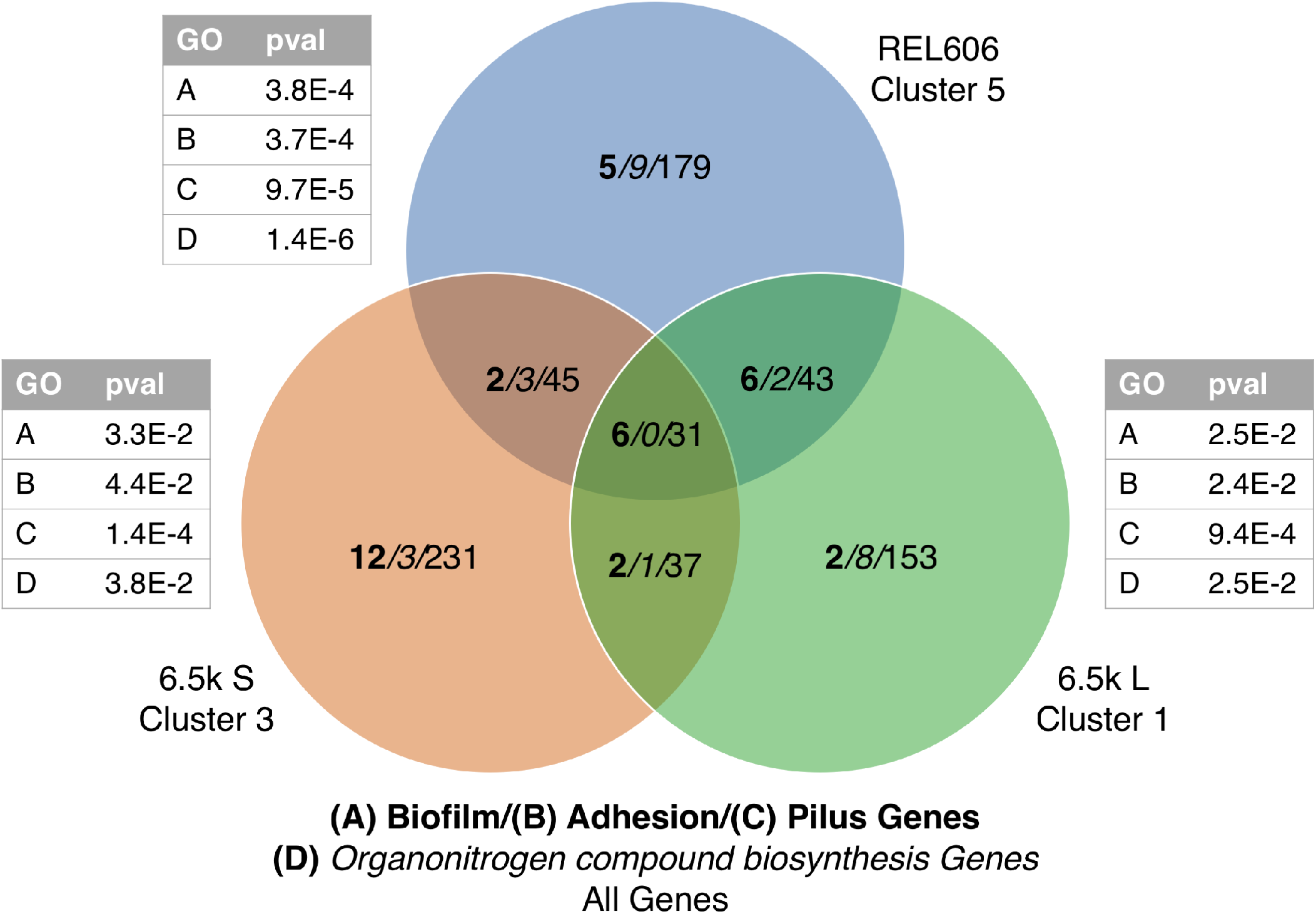
Biofilm (GO:0043708)/adhesion (GO:0022610)/ pilus organization (GO:0043711)/ organonitrogen compound biosynthesis (GO:1901566) genes tend to appear in the same clusters across genetic backgrounds. P-values are post-FDR correction.

**Fig. S23.**
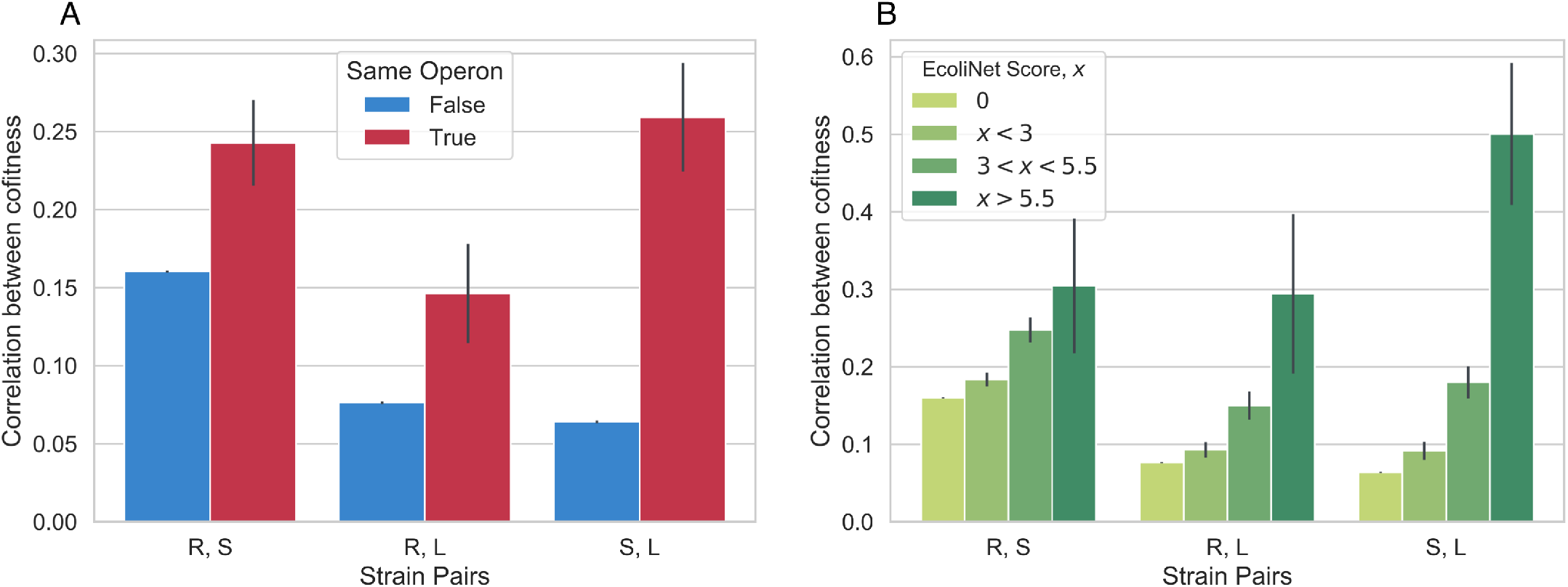
Correlations between cofitness increase (**A**) when genes are in the same operon and (**B**) with EcoliNet (58) score. A score of 0 indicates that the gene pair is not connected in EcoliNet, i.e. a node distance greater than 1.

**Fig. S24.**
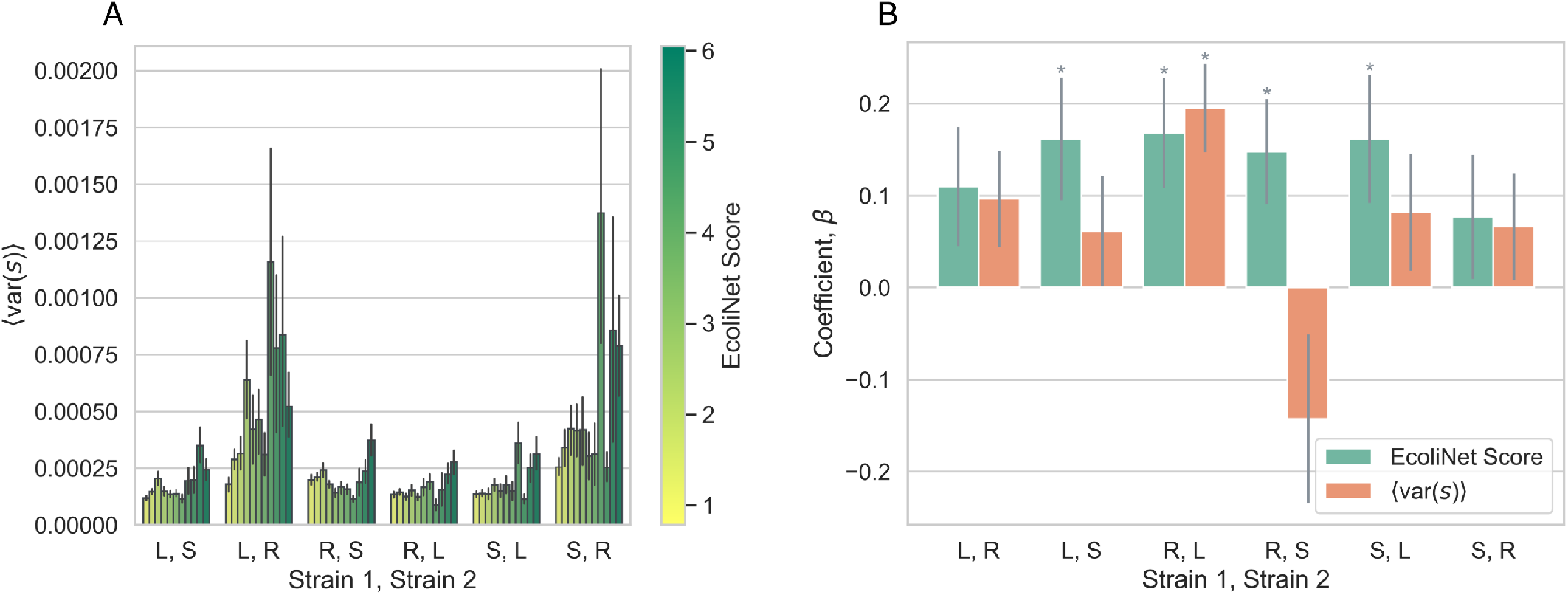
Variance in fitness effect across environment does not fully explain correlation between EcoliNet score and probability that two genes will be in the same cluster across strains. (**A**) Covariation of EcoliNet scores and variations in fitness effects in some strain pairs. The observed covariation is interesting in and of itself, as it suggests that more strongly interacting genes tend to have a larger variation in fitness effects across environments. (**B**) A standard multiple logistic regression with both fitness variance (in strain 2) and EcoliNet score as covariates, with response variable as the probability two genes are in the same cluster in strain 2, if they were together in strain 1. For most strain pairs, the regression reveals that the correlation reported in Figure 4D still holds after controlling for variation in fitness effects. * *p <* 0.05. See section S4.2 for model details.

**Fig. S25.**
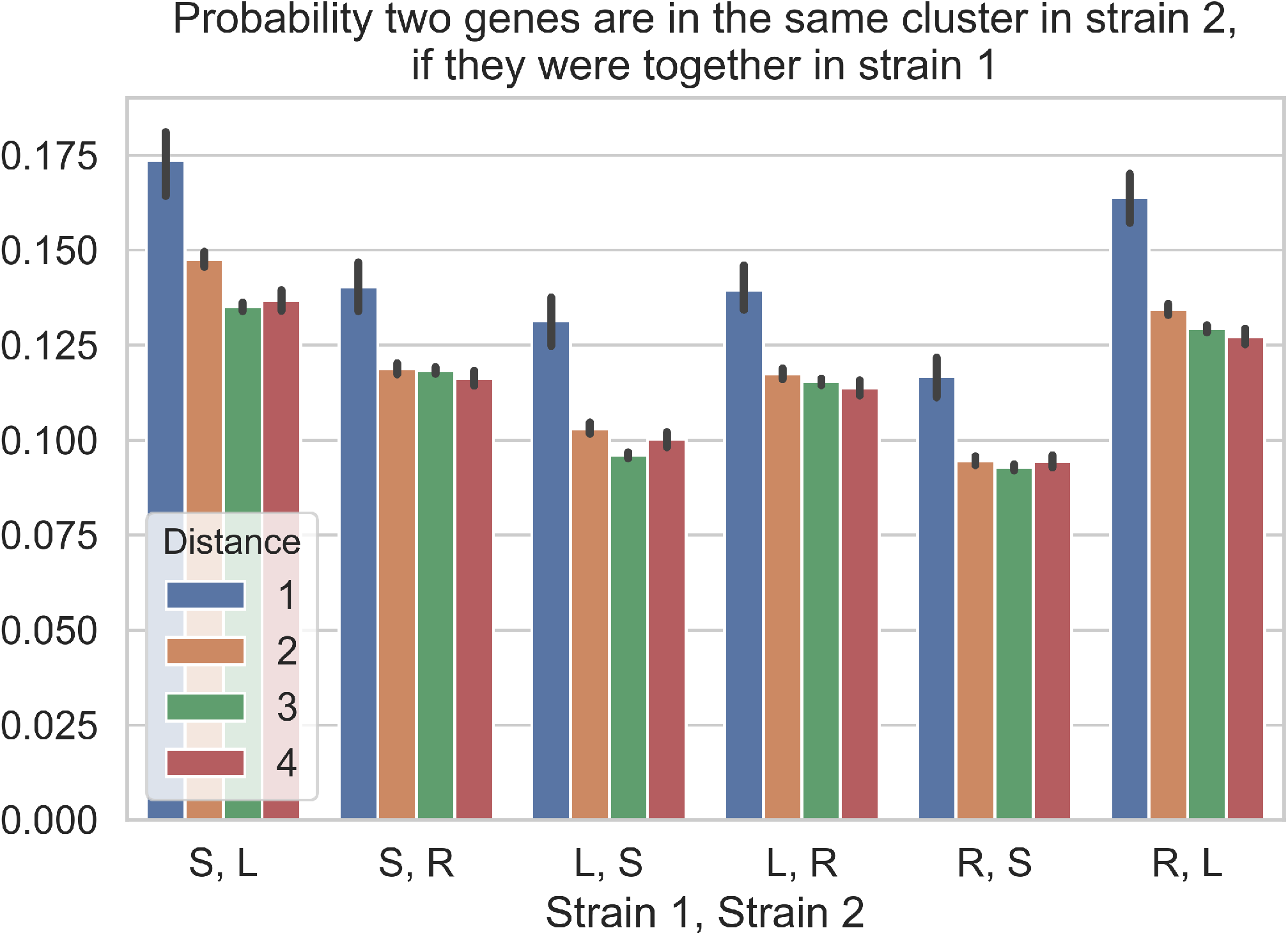
Shortest distance between genes in EcoliNet (58) predicts if genes stay correlated across genetic background

**Fig. S26.**
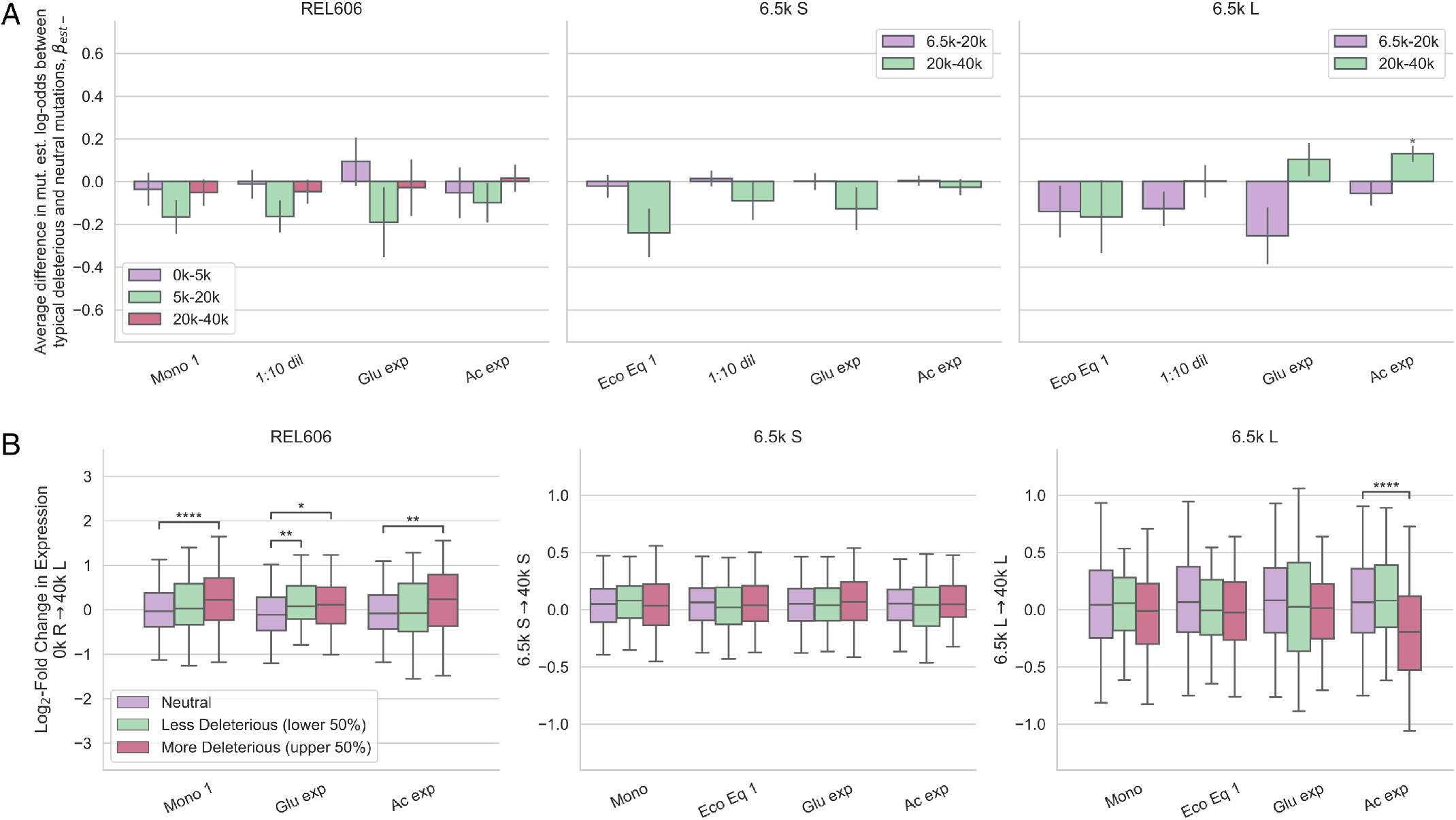
Relationship between deleterious mutations and evolutionary outcomes. (**A**) Deleterious fitness effects generally do not predict which genes will mutate, with the one exception that L seems more likely than random to get mutations in genes with deleterious acetate knockout fitness. (**B**) In REL606, deleterious knockout fitness effects are predictive of increased gene expression across all tested environments. Asterisks denote coefficients/comparisons that are significantly different than 0 (* *p <* 0.05, ** *p <* 0.01, *** *p <* 0.001).

**Fig. S27.**
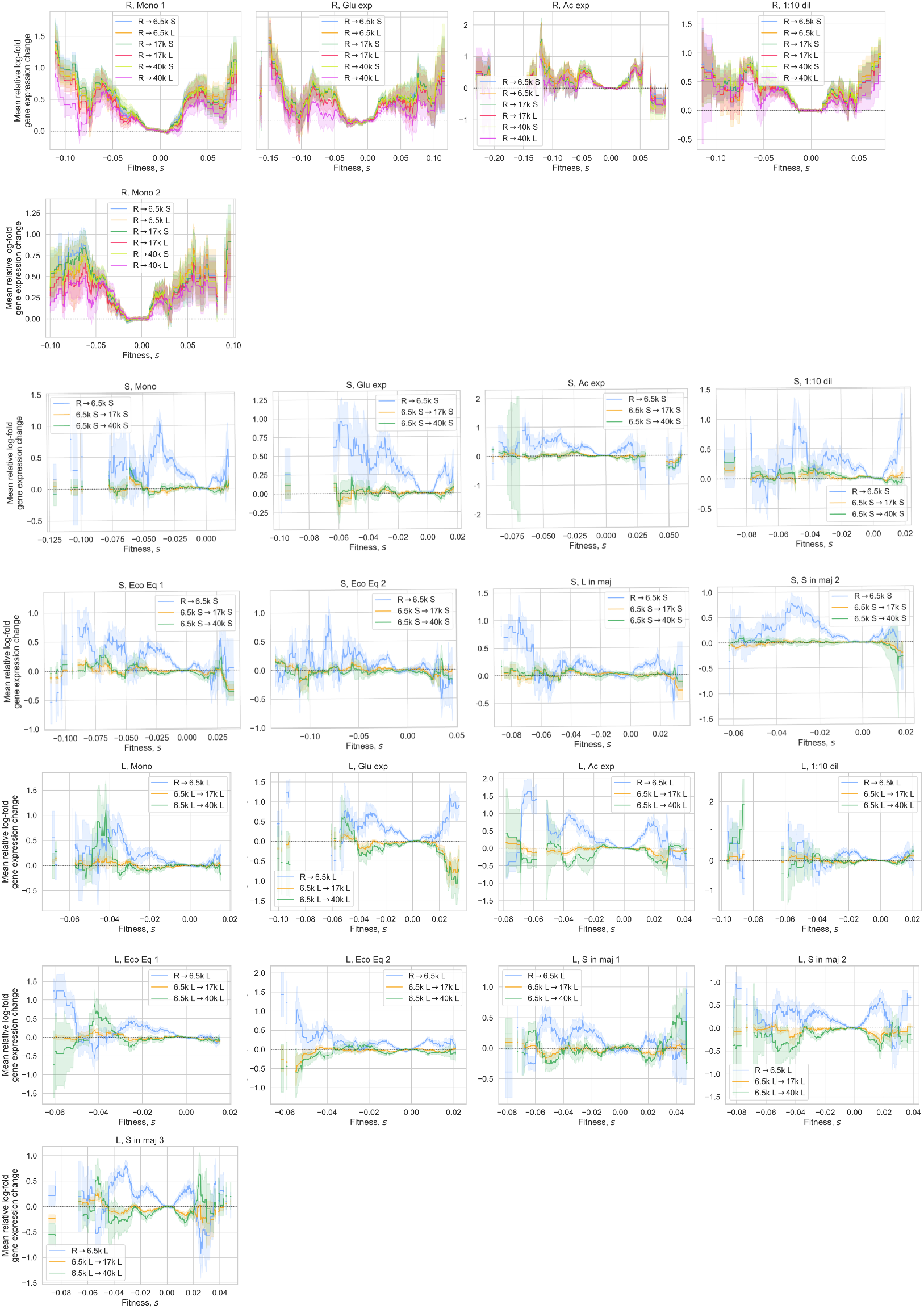
Relationship between fitness effects and log-fold gene expression change for all experiments, relative to the average change for neutral knockouts. Lines show mean gene expression change as a function of fitness effect (± standard error), with a ±0.01 moving average.

**Fig. S28.**
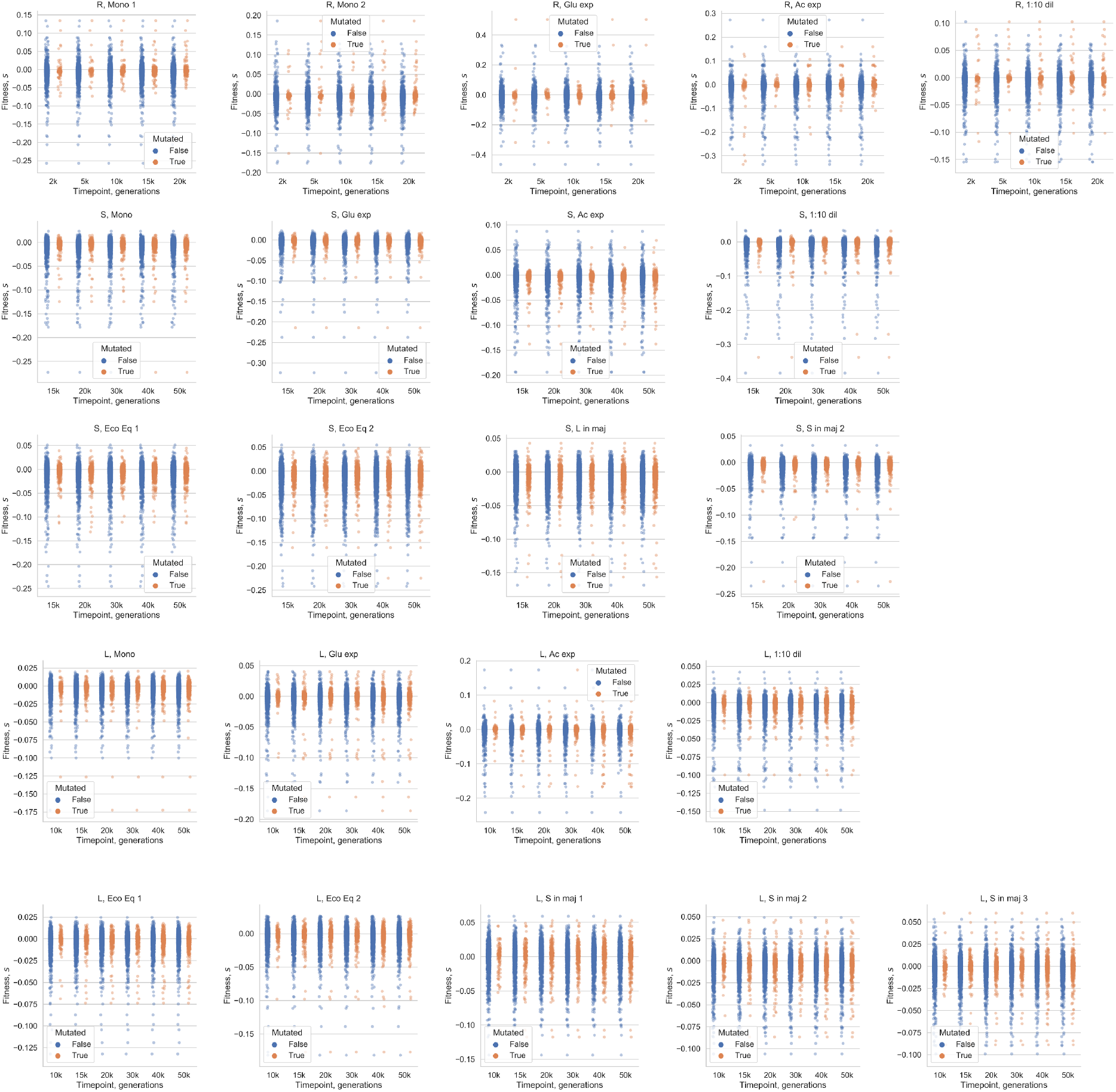
Establishment of a mutation in a gene by its knockout fitness.

**Fig. S29.**
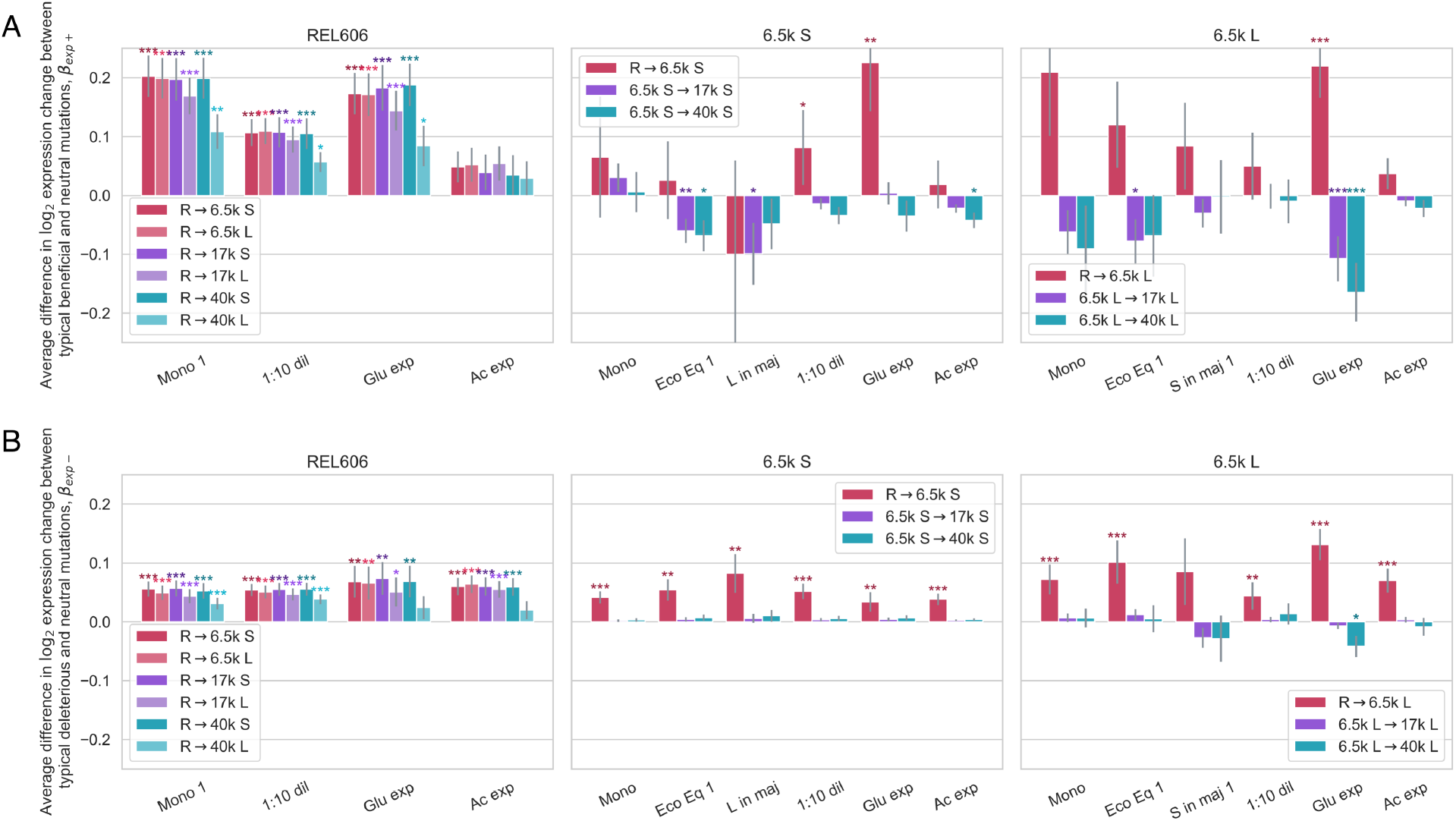
Linear model to explore correlation of gene expression changes with knockout fitness effects, comparing neutral to (a) beneficial and (b) deleterious knockouts. Asterisks denote coefficients that are significantly different than 0 (FDR correction; * *p <* 0.05, ** *p <* 0.01, *** *p <* 0.001).

**Fig. S30.**
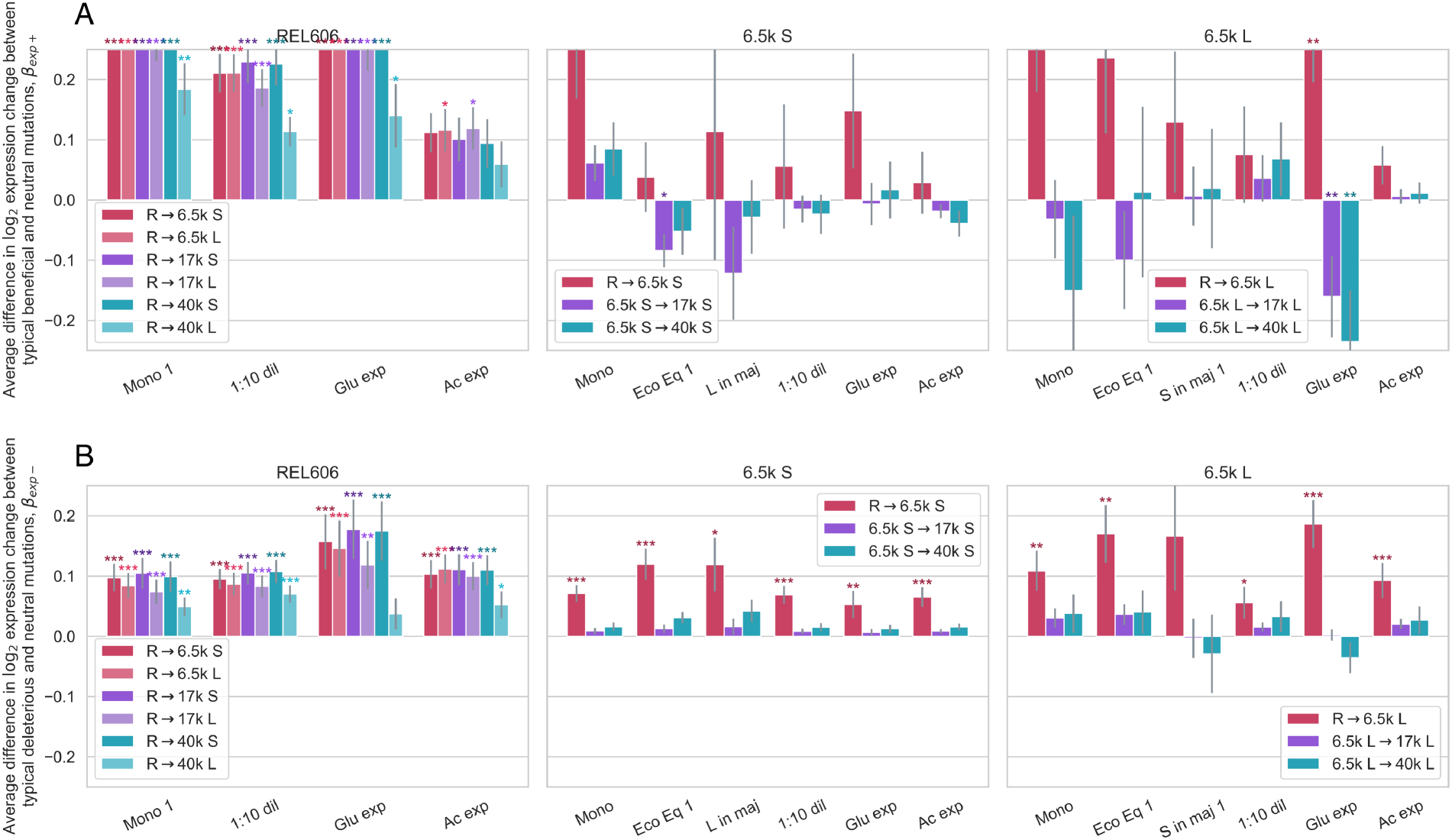
Restricting analysis of Figure S29 to exclude poorly expressed genes (bottom 50%) does not qualitatively change results of analysis, when comparing neutral to (a) beneficial and (b) deleterious knockouts.

**Fig. S31.**
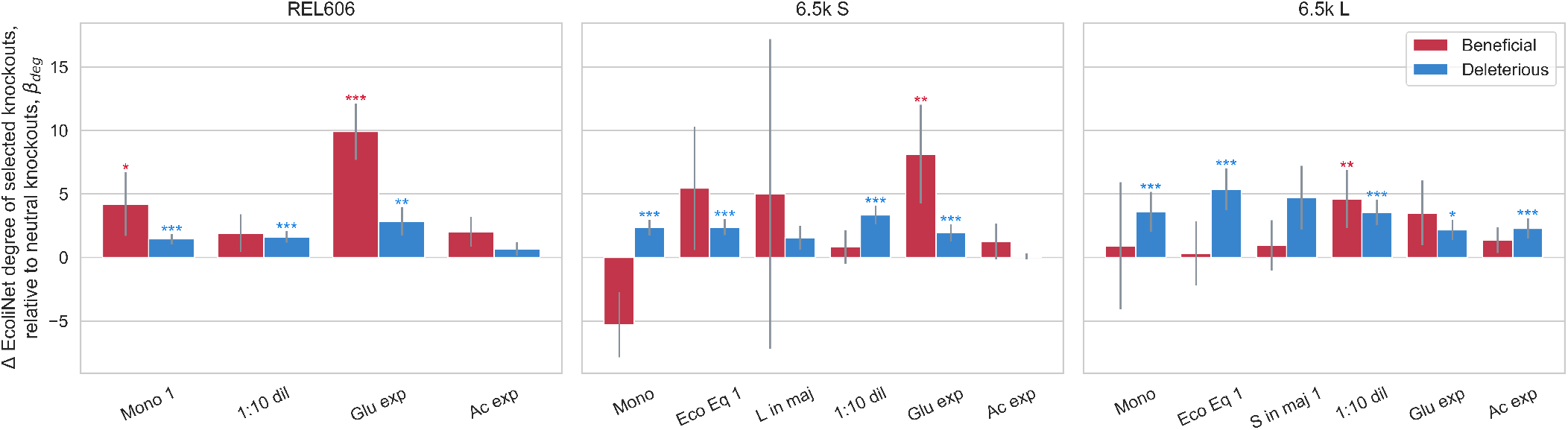
Fitness effects predict EcoliNet node degree. Deleterious knockouts across environments are more likely to have a high degree compared to neutral knockouts. The same general pattern appears for beneficial knockouts, although less clearly. Linear model fit with ordinary least squares; normalized analogous to model in section S4.4. Asterisks denote coefficients that are significantly different than 0 (* *p <* 0.05, ** *p <* 0.01, *** *p <* 0.001).

## Notes

### Competing Interest Statement

The authors have declared no competing interest.

### Summary of Updates

Figures 3/5 revised; introduction significantly changed; additional controls added

